# Accounting for epistasis improves genomic prediction of phenotypes with univariate and bivariate models across environments

**DOI:** 10.1101/2020.10.08.331074

**Authors:** Elaheh Vojgani, Torsten Pook, Johannes W.R. Martini, Armin C. Hölker, Manfred Mayer, Chris-Carolin Schön, Henner Simianer

## Abstract

We compared the predictive ability of various prediction models for a maize dataset derived from 910 doubled haploid lines from European landraces (Kemater Landmais Gelb and Petkuser Ferdinand Rot), which were tested in six locations in Germany and Spain. The compared models were Genomic Best Linear Unbiased Prediction (GBLUP) as an additive model, Epistatic Random Regression BLUP (ERRBLUP) accounting for all pairwise SNP interactions, and selective Epistatic Random Regression BLUP (sERRBLUP) accounting for a selected subset of pairwise SNP interactions. These models have been compared in both univariate and bivariate statistical settings within and across environments. Our results indicate that modeling all pairwise SNP interactions into the univariate/bivariate model (ERRBLUP) is not superior in predictive ability to the respective additive model (GBLUP). However, incorporating only a selected subset of interactions with the highest effect variances in univariate/bivariate sERRBLUP can increase predictive ability significantly compared to the univariate/bivariate GBLUP. Overall, bivariate models consistently outperform univariate models in predictive ability. Over all studied traits, locations, and landraces, the increase in prediction accuracy from univariate GBLUP to univariate sERRBLUP ranged from 5.9 to 112.4 percent, with an average increase of 47 percent. For bivariate models, the change ranged from −0.3 to +27.9 percent comparing the bivariate sERRBLUP to the bivariate GBLUP. The average increase across traits and locations was 11 percent. This considerable increase in predictive ability achieved by sERRBLUP may be of interest for “sparse testing” approaches in which only a subset of the lines/hybrids of interest is observed at each location.

**Key Message:** The prediction accuracy of genomic prediction of phenotypes can be increased by only including top ranked pairwise SNP interactions into the prediction models.

## Introduction

Genomic prediction of phenotypes has been widely explored for crops (Crossa *et al*., 2010), livestock (Daetwyler *et al*., 2013) and clinical research (de los Campos *et al*., 2013). Broad availability and cost effective generation of genomic data had a considerable impact on plant (Bernardo and Yu, 2007; de los Campos *et al*., 2009; Crossa *et al*., 2010, 2011; de Los Campos *et al*., 2010; Pérez *et al*., 2010) and animal breeding programs (de los Campos *et al*., 2009; Hayes and Goddard, 2010; Daetwyler *et al*., 2013). Genomic prediction relates a set of genome wide markers to the variability in the observed phenotypes and enables the prediction of phenotypes or genetic values of genotyped but unobserved material (Meuwissen *et al*., 2001; Jones, 2012; Windhausen *et al*., 2012). This approach has been positively evaluated in most major crop and livestock species (Albrecht *et al*., 2011; Daetwyler *et al*., 2013; Desta and Ortiz, 2014) and is becoming a routine tool in commercial and public breeding programs (Stich and Ingheland, 2018). In plant breeding, phenotyping is one of the major current bottlenecks and the optimization or minimization of phenotyping costs within breeding programs is needed (Akdemir and Isidro-Sánchez, 2019). Therefore, the maximization of genomic prediction accuracy can be directly translated into reduced phenotyping costs (Akdemir and Isidro-Sánchez, 2019; Jarquin *et al*., 2020).

Genomic selection and the corresponding prediction of breeding values is based on a covariance matrix describing the (additive) relationship between the considered individuals (Wolc *et al*., 2011; Burgueño *et al*., 2012). This matrix can be constructed from pedigree information, from marker information (Haley and Visscher, 1998; Hayes and Goddard, 2008) or from a combination of pedigree and available genotypic information in a single step approach (Aguilar *et al*., 2010; Legarra *et al*., 2014). It has been broadly demonstrated that marker based relationship matrices enhance the reliability of breeding value estimation on average across traits and compared to pedigree based approaches (Meuwissen *et al*., 2001; VanRaden, 2007; Hayes and Goddard, 2008; Crossa *et al*., 2010).

One of the first broad applications of genomic prediction was to select young sires with high breeding values earlier in their life span, without the need for information of the performance of their progeny in dairy cattle breeding (Schaeffer, 2006; VanRaden, 2007). Since breeding values are additive by definition (Falconer and Mackay, 1996), the early development of prediction models exclusively accounted for the additive effects (Filho *et al*., 2016).

Concerning additive models, genomic best linear unbiased prediction (GBLUP, Meuwissen *et al*., 2001; VanRaden, 2007) is a widely-used linear mixed model (Da *et al*., 2014; Rönnegård and Shen, 2016; Covarrubias-Pazaran *et al*., 2018). The computational steps involved in GBLUP are much faster than Bayesian methods and it has been difficult to find a method which consistently outperforms GBLUP when predicting complex traits (Wang *et al*., 2018). Daetwyler *et al*. (2010) showed that BayesB can yield higher accuracy than GBLUP for traits controlled by a small number of quantitative trait nucleotides, emphasizing that the genetic architecture of the trait has an important impact on which method may predict better (Wimmer *et al*., 2013; Momen *et al*., 2018). Moreover, the training set size was shown to play a role. For instance, human height prediction using BayesB and BayesC methods in a small reference population (<6,000 individuals) had no advantage over GBLUP. Only when increasing the size of the reference population (>6,000 individuals), these methods outperformed GBLUP (Karaman *et al*., 2016).

Understanding how genetic variation causes phenotypic variation in quantitative traits is still a major challenge of contemporary biology. It has been proven that epistasis as a statistical interaction between two or more loci (Falconer and Mackay, 1996) contributes substantially to the genetic variation of quantitative traits (Wright, 1931; Carlborg and Haley, 2004; Hill *et al*., 2008; Huang *et al*., 2012; Mackay, 2014). On the one hand, models which incorporate epistasis have the potential to increase predictive ability (de Los Campos *et al*., 2010; Hu *et al*., 2011; Wang *et al*., 2012; Mackay, 2014). On the other hand, accounting for epistasis by modeling interactions explicitly was considered to be computationally challenging (Mackay, 2014). In this context, the extended genomic best linear unbiased prediction (EG-BLUP) as an epistasis marker effect model was proposed to reduce the computational load by constructing marker-based epistatic relationship matrices (Jiang and Reif, 2015; Martini *et al*., 2016). While EG-BLUP is potentially beneficial for genomic prediction, its performance depends on the marker coding (Martini *et al*., 2017, 2019), and the Hadamard products of the additive genomic relationship matrices provide only an approximation for the interaction effect model based on interactions between individual loci (Martini *et al*., 2020). Moreover, it has been shown that the superiority of epistasis models over the additive GBLUP in terms of predictive ability may vanish when the number of markers increases (Schrauf *et al*., 2020).

Another downside of epistasis models is that, due to the high number of interactions, a large number of unimportant variables can be introduced into the model (Rönnegård and Shen, 2016). This ‘noise’ might prevent a gain in predictive ability. In this regard, Martini *et al*. (2016) showed that selecting just a subset of the largest epistatic interaction effects has the potential to improve predictive ability. Therefore, reducing the full epistasis model to a model based on a subnetwork of ‘most relevant’ pairwise SNP interactions may be beneficial for prediction performance (Martini *et al*., 2016).

In addition to the extension from additive effect models to models including epistatic interactions, genomic prediction models can be extended from univariate models to multivariate models. Univariate models consider each trait separately, while multivariate models treat several traits simultaneously with the objective to exploit the genetic correlation between them to increase predictive ability. Multivariate models which have been first proposed for the prediction of genetic values by Henderson and Quaas (1976) were shown to be potentially beneficial for prediction accuracy when the correlation between traits is strong (He *et al*., 2016; Covarrubias-Pazaran *et al*., 2018; Schulthess *et al*., 2018; Velazco *et al*., 2019). A situation of dealing with multiple environments can also be considered in the framework of a multivariate model by simply considering a trait-in-environment combination as another correlated trait. Therefore, prediction accuracy could be potentially enhanced through borrowing information across environments by utilizing multi-environment models (Burgueño *et al*., 2012). In addition to multi-environment models, Martini *et al*. (2016) showed that predictive ability of a univariate model can be increased in one environment by variable selection in the other environment under the assumption of a relevant correlation of phenotypes in different environments. This, however, was only demonstrated with a data set of limited size and especially a limited set of markers and, thus, marker interactions.

In the present study, we use a data set of two doubled haploid populations derived from two European landraces, to investigate how beneficial the use of subnetworks of interactions in the proposed sERRBLUP framework can be. This was compared in the context of univariate and bivariate models. We assess the optimum proportion of SNP interactions to be kept in the model in the variable selection step. The underlying phenotypic data was generated in multi-location trials, and we assessed whether the different univariate and bivariate models have a potential to provide benefits in across location predictions. The development of efficient selection strategies which could mitigate costly and time consuming phenotyping of a large number of selection candidates in multiple environments has been a particular focus of research in plant breeding (Jarquin *et al*., 2020). A successful application of our models may reduce costs of phenotyping by reducing the number of test locations per line.

## Materials and Methods

### Data used for analysis

We used a set of 501 / 409 doubled haploid lines of the European maize landraces Kemater Landmais Gelb / Petkuser Ferdinand Rot genotyped with 501,124 markers using the Affymetrix ^®^ Axiom Maize Genotyping Array (Unterseer *et al*., 2014), out of which 471 and 402 lines were phenotyped for Kemater and Petkuser, respectively.

The lines were phenotyped in 2017 for a series of traits in six different environments which were Bernburg (BBG, Germany), Einbeck (EIN, Germany), Oberer Lindenhof (OLI, Germany), Roggenstein (ROG, Germany), Golada (GOL, Spain) and Tomeza (TOM, Spain). The weather data which were obtained from 15th of April to 30th of September 2017 are provided by Hölker *et al*. (2019).

The phenotypic trait description and the mean, standard deviation, maximum and minimum values of each trait across all six locations in 2017 and both landraces are provided in Table 1. Moreover, these values are shown in the supplementary for each environment and for each landrace separately (Table S1). The vegetative growth stage is the corn growth stage based on the leaf collar method representing the number of visible leaf collars. To illustrate this, V4 indicates the growth stage at which four leaf collars are fully developed (Abendroth *et al*., 2011). In our study, we consider PH_V4 as the main trait for evaluating our methods, since it is a metric quantitative trait for early plant development which is suitable to use for testing our methods. The phenotypic correlations of PH_V4 across all environments are provided in Table 2.

**Table 1.** Phenotypic traits description and the mean, minimum, maximum and standard deviation for all landraces.

| Trait | Definition | Mean | Minimum | Maximum | Standard deviation |
| --- | --- | --- | --- | --- | --- |
| **EV_V3** | Early vigour at V3 stage scored on scale from 1 (very poor early vigour) to 9 (very high early vigour) | 5.22 | 0.79 | 9.01 | 1.32 |
| **EV_V4** | Early vigour at V4 stage scored on scale from 1 (very poor early vigour) to 9 (very high early vigour) | 5.12 | 0.7 | 8.49 | 1.3 |
| **EV_V6** | Early vigour at V6 stage scored on scale from 1 (very poor early vigour) to 9 (very high early vigour) | 5.36 | 0.4 | 8.86 | 1.25 |
| **PH_V4** | Mean plant height of three plants of the plot at V4 stage in cm | 35.34 | 6.67 | 95.35 | 14.65 |
| **PH_V6** | Mean plant height of three plants of the plot at V6 stage in cm | 65.57 | 6.73 | 130.37 | 20.04 |
| **PH_final** | Final plant height after flowering in cm | 131.73 | 29.33 | 241.96 | 27.05 |
| **FF** | Days after sowing until female flowering (days until 50% of the plot showed silks) | 79.71 | 59.12 | 102.22 | 6.46 |
| **RL** | Root lodging score from 1 to 9 (1 belonged to no lodging and 9 belonged to severe lodging) | 2.83 | 0.27 | 9.38 | 2.29 |

**Table 2.** Phenotypic correlation across all environments for the trait PH_V4.

| Location | EIN | OLI | ROG | GOL | TOM |
| --- | --- | --- | --- | --- | --- |
| BBG | 0.817 | 0.660 | 0.674 | 0.689 | 0.575 |
| EIN | - | 0.707 | 0.767 | 0.747 | 0.662 |
| OLI | - | - | 0.711 | 0.649 | 0.503 |
| ROG | - | - | - | 0.703 | 0.575 |
| GOL | - | - | - | - | 0.693 |

Among the phenotypic traits, root lodging (RL) and female flowering (FF) were not phenotyped in all the environments: RL was only scored in BBG, ROG, OLI and EIN. FF was phenotyped in all environments except GOL.

### Quality control, coding and imputing

In total, 948 DH lines from the landraces Kemater (KE) and Petkuser (PE) were genotyped using the 600k Affymetrix^®^ Axiom^®^ Maize Array (Unterseer et al. 2014). After stringent quality filtering, as described in Hölker *et al*. (2019), the dataset was reduced to 501,124 SNPs and 910 DH lines from KE and PE. Remaining heterozygous calls were set to missing. Genotype calls were coded according to the allele counts of the B73 AGPv4 reference sequence (Jiao *et al*., 2017) (0 = homozygous for the reference allele, 2 = homozygous for the alternative allele). Imputation of missing values was performed separately for each landrace, using BEAGLE version 4.0 with parameters buildwindow=50, nsamples=50 (Browning and Browning, 2007; Pook *et al*., 2020). As the dataset only included doubled haploid lines and heterozygous calls were not expected, the DS (dosage) information of the BEAGLE output was used to recode remaining heterozygous calls. The genotype was then coded as 0 if DS < 1 and as 2 if DS >= 1.

### Linkage disequilibrium pruning

Linkage disequilibrium based SNP pruning with PLINK v1.07 was used to generate a subset of SNPs which are in approximate linkage equilibrium with each other. The parameters: indep 50 5 2 were used, in which 50 was the window size in SNPs, 5 was the number of SNPs to shift the window at each step and 2 was the variance inflation factor *VIF* = 1/(1 − *r*^2^), where *r*^2^ was the squared correlation between single SNPs and linear combinations of all SNPs in the window. All variants in the 50 SNP window which had a VIF > 2 were removed. Then, the window was shifted 5 SNPs forward and the procedure was repeated (Purcell et al. 2007; Chang et al. 2015).

In our study, LD pruning was done separately for each landrace, resulting in data panels containing 25’437 SNPs for KE and 30’212 SNPs for PE.

### Univariate statistical models for phenotype prediction

We used three different statistical models to predict phenotypes, which are all based on the same linear mixed model (Henderson 1975). We assume that we have in total *n* lines which are genotyped, and phenotypes are available for a subset of *n*_1_ lines. These *n*_1_ lines are used to train the model and missing phenotypes for the remaining *n*_2_ = *n* − *n*_1_ lines are predicted by using the genotypes of these lines. The basic univariate model is

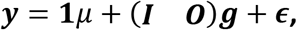

where ***y*** is an *n*_1_ × 1 vector of phenotypes, **1** is an *n*_1_ × 1 vector with all entries equal to 1, *μ* is a scalar fixed effect, ***I*** is an identity matrix of dimension *n*_1_ × *n*_1_ and ***O*** is a matrix of dimension *n*_1_ × *n*_2_ of zeros. The design matrix (***I O***) is the *n*_1_ × (*n*_1_ + *n*_2_) matrix resulting from the concatenation of ***I*** and ***O***. Moreover, 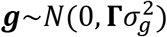 is an *n* × 1 vector of random genetic effects, and 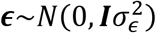 is a random error vector, where **Γ** and ***I*** are the respective dispersion matrices and 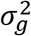 and 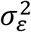 are the corresponding variance components.

With this model, the population mean and the genetic effects ***g*** for all lines, including those without phenotypes, are estimated using

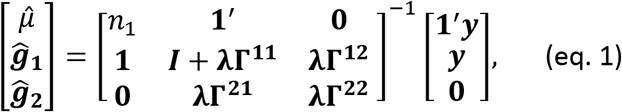

where 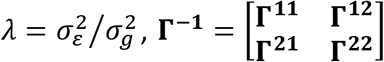 and 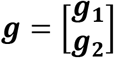 and the indices pertain to the subset of individuals with (index 1) or without (index 2) phenotypes, respectively.

With these solutions, the phenotypes for the set of unphenotyped individuals can be predicted as 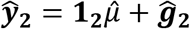, where 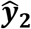 is the *n*_2_ × 1 vector of predicted phenotypes and **1**_2_ is an *n*_2_ × 1 vector of ones.

For *n* = *n*_1_ and *n*_2_ = 0 the solution of eq. 1 provides estimates of genetic effects when all lines are phenotyped and genotyped.

### Bivariate statistical models for phenotype prediction

Besides univariate models, we also used bivariate models, where the two variables represent the same trait measured in two different environments.

The basic bivariate model is

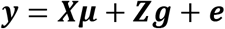

or, in more detail,

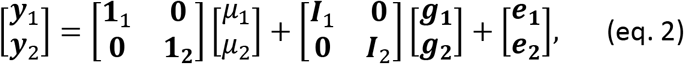

where, 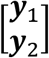 is the phenotype vector of length *m* = *m*_1_ + *m*_2_ for environment 1 (*m*_1_) and 2 (*m*_2_), **1_1_** and **1_2_** are respectively *m*_1_ × 1 and *m*_2_ × 1 vectors with all entries equal to 1, 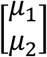 is the vector of population means for environment 1 and 2, ***I***_1_ and ***I***_2_ are identity matrices of dimension *m*_1_ × *m*_1_ and *m*_2_ × *m*_2_, respectively assigning genomic values to phenotypes. Moreover, 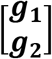 is the vector of random genomic values which is assumed to have a multivariate normal distribution with mean zero and variance ***G*** = ***H*** ⊗ **Γ**, where 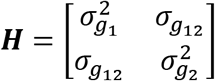, **Γ** is the dispersion matrix of genetic effects and ⊗ is the Kronecker product; 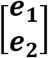 is the vector of random errors which is assumed to have a multivariate normal distribution with mean zero and variance ***R*** = ***R***_0_ ⊗ ***I***, where 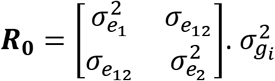 and 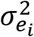 represent the genetic and residual variance of environment *i* = 1,2, and 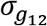 and 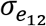 are the genetic and residual covariance between the environment 1 and 2 (Guo *et al*., 2014). In this model, the phenotypes have to be ordered in the same way in both environments. In the case that the number of observations in environment 1 and environment 2 not being identical (i.e. in general terms *m*_1_ ≠ *m*_2_) or not having the same lines in both environments in the model, the incidence matrices have to be adapted accordingly.

With this model, the vector of environment specific population means and the vector of genetic effects for all lines are estimated using the standard mixed model equations

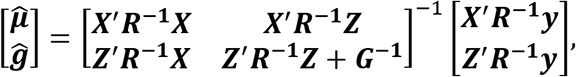

In analogy to the procedure described in the univariate setting, we consider a setting in which the last *l* phenotypes for environment 2 are masked and predicted from all observations in environment 1 and the first *k* = *m*_2_ − *l* non-masked observations in environment 2.

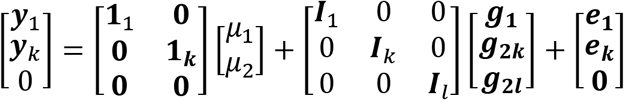

From the solutions obtained with this model, the phenotypes for the set of unphenotyped individuals in environment 2 can be predicted as 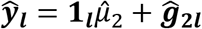, where 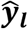 is the *l* × 1 vector of predicted phenotypes and **1**_*l*_ is an *l* × 1 vector of ones.

The three models compared in this study only differ in the choice of the dispersion matrix **Γ** of the genetic effects.

### Model 1: Genomic Best Linear Unbiased Prediction (GBLUP)

In this additive model we use as **Γ** the genomic relationship matrix which is calculated according to VanRaden (2008) as

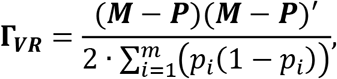

where ***M*** is the *n* × *m* marker matrix which gives *m* marker values for *n* lines under the assumption of having *n* genotyped lines in total. ***P*** is a matrix of equal dimension as ***M*** with 2 · *p*_*i*_ in the *i*^*th*^column, and *p*_*i*_ is the allele frequency of the minor allele of SNP *i*.

### Model 2: Epistatic Random Regression BLUP (ERRBLUP)

This model accounts for all possible SNP interactions in the prediction model. With m markers and fully inbred lines, we have two possible genotypes at a single locus, i.e. 0 or 2 when coded as the counts of the minor allele. For each pair of loci, we have four different possible genotype combinations: {00, 02, 20, 22}. The total number of pairs of loci is 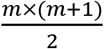 if we allow for interaction of a locus with itself. Since for each of these pairs we have four possible genotype combinations, the total number of combinations to be considered as dummy variables is 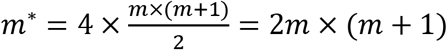.

We define a marker combination matrix ***M***\* of dimension *n* × *m*\* whose element *i*, *j* is 1 if genotype combination *j* is present in individual *i* and is 0 otherwise. We further define for column *i* of this matrix the average value 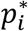, giving the frequency of the respective genotype combination in the population, and a matrix ***P***\* being of equal dimension as ***M***\* with 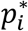 in the *i*^*th*^column.

Then, the relationship matrix based on all SNP interactions was calculated according to VanRaden (2008) as

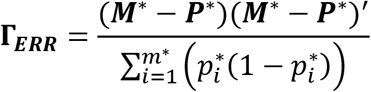

and this matrix was used in ERRBLUP as dispersion matrix for the genetic effects, which now are based on epistatic interaction effects. It should be noted that including the interaction of each locus with itself replaces the additive effect, so that it is not necessary to use a model that separately accounts for additive and epistatic effects. This model had been introduced earlier as “categorical epistasis model” (Martini *et al*., 2017).

### Model 3: selective Epistatic Random Regression BLUP (sERRBLUP)

sERRBLUP is based on the same approach as ERRBLUP, but here the **Γ** -matrix is constructed from a selected subset of genotype interactions. We decided to use those interactions with the highest estimated marker effects variances. Selection based on highest absolute effects (as used by Martini *et al*. (2016) in the framework of the EGBLUP epistasis model) was also considered, but lead to similar to slightly worse results. For this, it was necessary to backsolve interaction effects 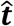 and effects variances 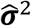 from the ERRBLUP model using (Mrode, 2014)

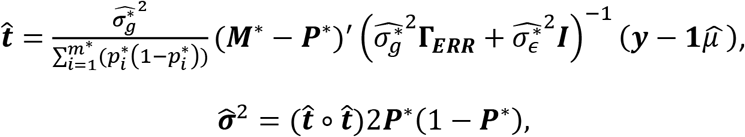

with ∘ denoting the Hadamard product.

After estimating SNP interaction effects in 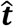 and effects variances in 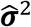, we selected those interactions whose absolute estimated effects or effect variances were in the top π = 0.05, 0.01, 0.001, 0.0001, 0.00001 or 0.000001 proportion of all interactions, respectively. These proportions were chosen since it was observed in preliminary analyses that they cover the most relevant range. For each of these subsets, we generated reduced matrices 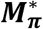 and 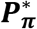 of dimension *n* × π*m*\*, containing only those columns of ***M***\* and ***P***\* pertaining to the selected subset of genotype interactions, and then set up the dispersion matrix in analogy to VanRaden (2008) as

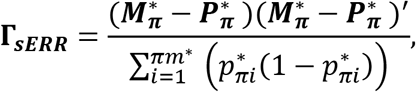

where 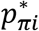 are the mean frequencies of the selected genotype combinations.

Note here that even for the univariate model, information from another environment is used for the prediction, namely for variable selection and the definition of **Γ**_***sERR***_. However, having used the information from another environment to define the subset of interactions and to derive the relationship matrix **Γ**_***sERR***_, the actual prediction is within the considered environment from the training to the test set.

We used the miraculix package (Schlather, 2020) to efficiently calculate **Γ**_***ERR***_, 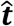 and **Γ**_***sERR***_.

### Assessment of predictive ability via 5-fold random cross validation with 5 replicates

In a 5-fold cross validation, the original sample is randomly partitioned into five subsamples of equal size. Out of the five subsamples, each subsample is subsequently considered as the test set for validating the model, and the remaining four subsamples are considered as training data. The training set is used to predict the test (validation) set. By this, all observations are used for both training and validation and each observation is only used once for validation (Utz *et al*., 2000). We repeated the cross-validation procedure 5 times, using random partitions of the original sample. The results of the 25 repetitions were then averaged (Erbe *et al*., 2010). We used the Pearson correlation between the predicted genetic value and the observed phenotype in the test set as measure for predictive ability. In our study, predictive ability was assessed for PE and KE for all phenotypic traits separately. In addition, the trait’s prediction accuracy was calculated by dividing the obtained predictive ability by the square-root of the respective trait’s heritability (Dekkers, 2007). The numbers of KE and PE lines which are available for all combinations of environments are summarized in Table 3. For some traits these numbers can be smaller or even zero for of some environment combinations.

**Table 3.**
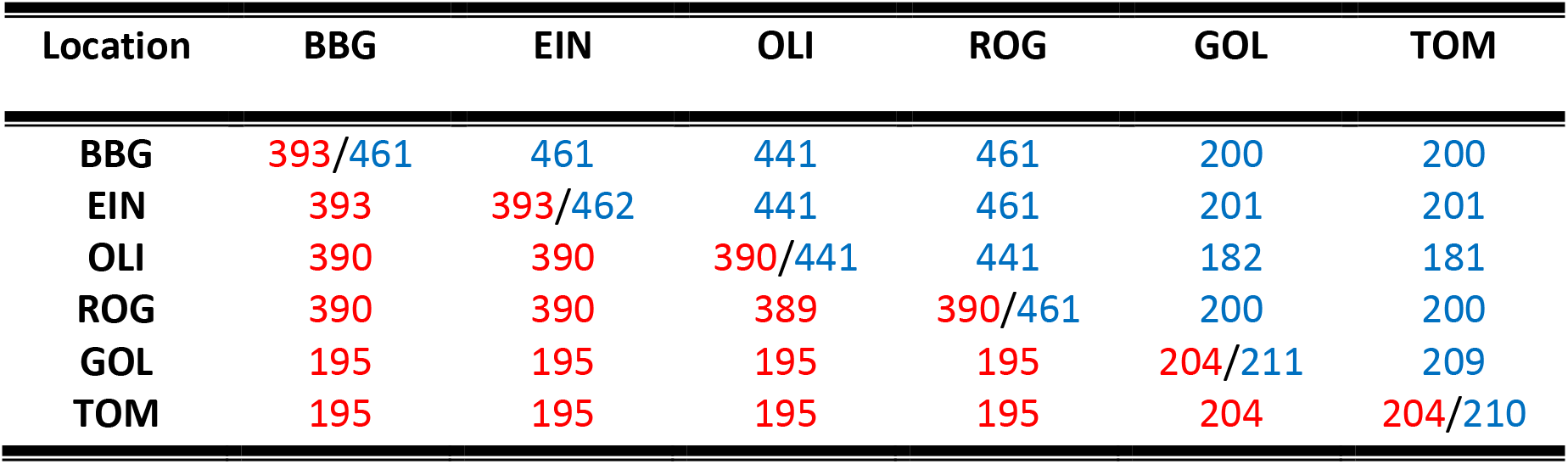
Number of Kemater (blue numbers above diagonal) and Petkuser (red numbers below diagonal) lines phenotyped in each pair of environments for trait PH_V4.

We evaluated our univariate and bivariate models as follows:

### Assessment of univariate GBLUP and ERRBLUP predictive abilities within environments

GBLUP and ERRBLUP within environments were evaluated by training the model in the same environment as the test set was sampled from.

### Assessment of univariate sERRBLUP predictive ability across environments

The basic strategy for sERRBLUP across environments is illustrated in Fig. 1: first, all pairwise SNP interaction effects and their variances are estimated from all data in environment 1 and effects are ordered either by absolute effect size or effect variance (A). Next, an epistatic relationship matrix for all lines is constructed from the top ranked subset of interaction effects (B). Then, this matrix is used in environment 2 (C) to predict phenotypes of the test set (green) from the respective training set (red) (D). This approach henceforth is termed ‘sERRBLUP across environments’.

**Fig. 1.**
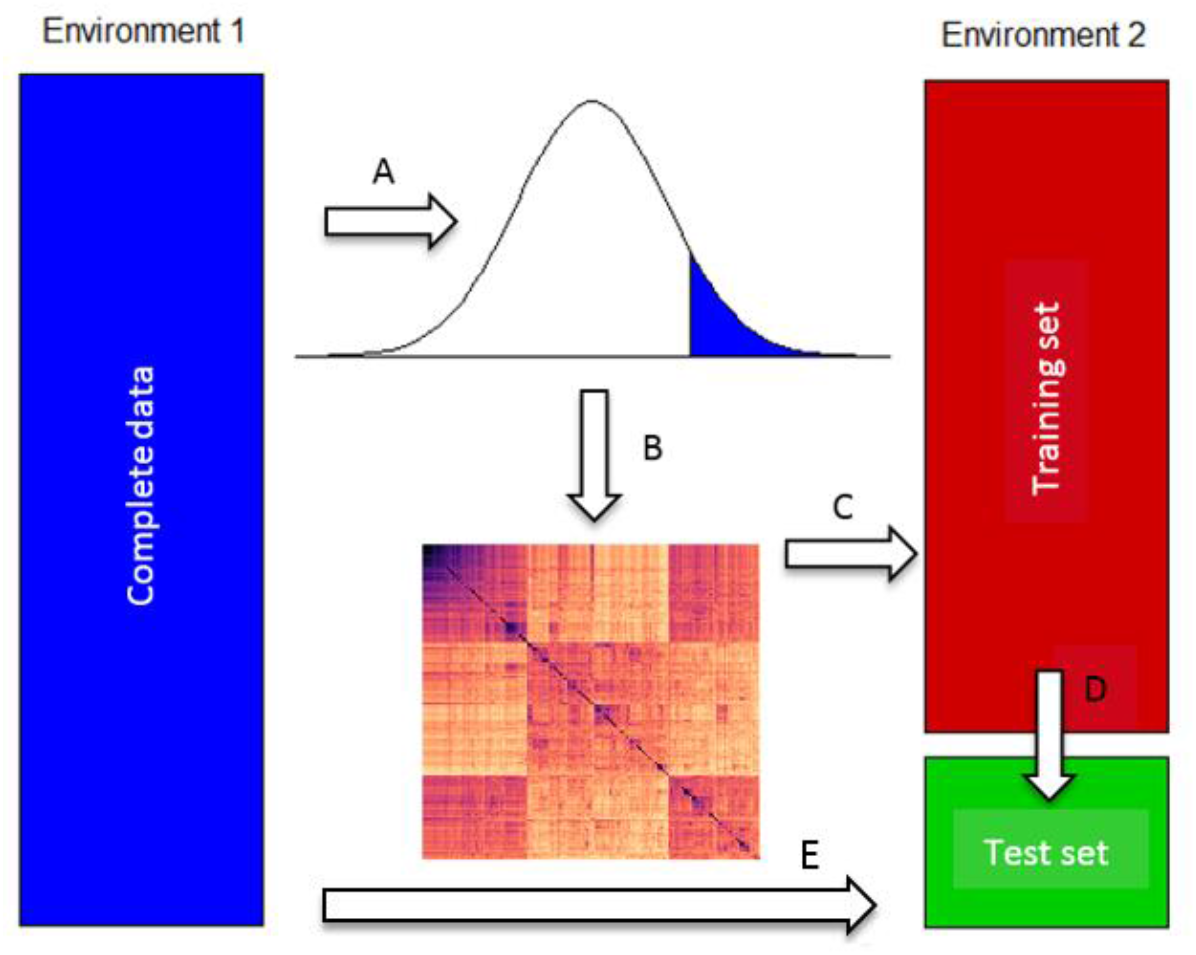
Basic scheme of uni- and bivariate sERRBLUP across environments. All pairwise SNP interaction effects and their variances are estimated from all data in environment 1, and effects are ordered either by absolute effect size or effect variance (A). Then, an epistatic relationship matrix for all lines is constructed from the top ranked subset of interaction effects (B) which in the univariate model is used in environment 2 (C) to predict phenotypes of the test set (green) from the respective training set (red, D). In the bivariate model, this information is combined with the complete data from environment 1 (blue, E) to predict the test set.

### Assessment of bivariate GBLUP, ERRBLUP predictive abilities

The basic strategy for bivariate GBLUP and ERRBLUP is illustrated in Fig. 2: The model is trained jointly on the complete dataset of environment 1 (A) and the training set of environment 2 (B). The test set of environment 2 is predicted, using as dispersion matrix for the genetic effects either **Γ**_***VR***_ or **Γ**_***ERR***_.

**Fig. 2.**
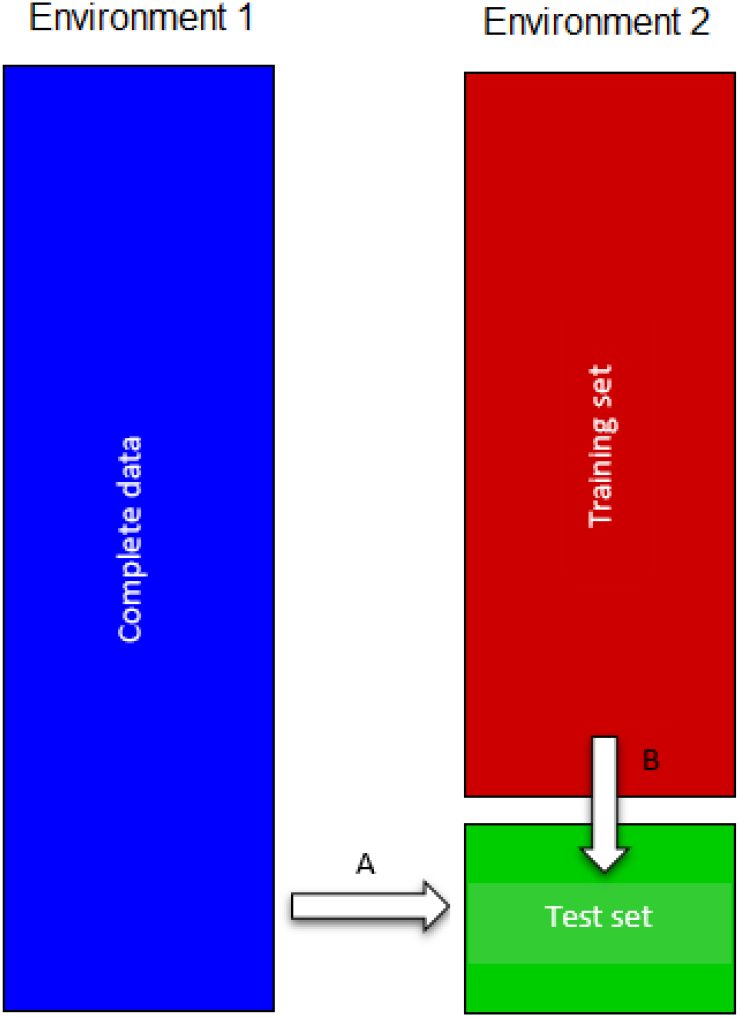
Basic scheme of bivariate GBLUP and ERRBLUP illustrating that the model is trained in both the complete data set of environment 1 (A) and the training set of environment 2 (B) to predict the test set in environment 2.

### Assessment of bivariate sERRBLUP predictive ability

The basic strategy of bivariate sERRBLUP is also illustrated in Fig. 1: first, all pairwise SNP interaction effects variances are estimated from univariate ERRBLUP model based on all data in environment 1, and effects are ordered by effect variance (A). Then, an epistatic relationship matrix **Γ**_***sERR***_ for all lines is constructed from the top ranked subset of interaction effects (B) This relationship matrix **Γ**_***sERR***_ is then used to predict the phenotypes of the test set (green) of environment 2 jointly from the training set of environment 2(red) (D) and the complete data set of environment 1 (blue) (E).

### Assessment of univariate sERRBLUP and bivariate GBLUP, ERRBLUP and sERRBLUP predictive ability across five environments jointly

In addition to assessing the predictive ability of univariate sERRBLUP when borrowing information from a single alternative environment, we considered five environments jointly for borrowing information to predict the sixth environment. Based on this approach, we combined five environments’ phenotypes by considering the mean of all five centered environments’ mean phenotypic values. The combined phenotypes for predicting BBG from EIN, OLI, ROG, GOL and TOM were jointly calculated as

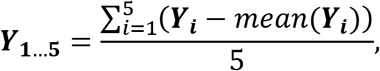

where ***Y***_**1**…**5**_ is the averaged phenotypes of EIN (***Y***_**1**_), OLI (***Y***_**2**_), ROG (***Y***_**3**_), GOL (***Y***_**4**_), TOM (***Y***_**5**_).

Borrowing information for variable selection across five environments jointly for univariate sERRBLUP follows the same strategy as borrowing information from a single environment for univariate sERRBLUP. Therefore, the newly generated phenotypes based on five environments (i.e. ***Y***_**1**…**5**_) were used to estimate the interaction effects and their variances, then the corresponding relationship matrices were constructed to be used for a prediction within the sixth (i.e. BBG (***Y***_**6**_)) environment. This approach henceforth is termed ‘univariate sERRBLUP across multiple environments jointly’.

This approach was also applied to bivariate GBLUP, ERRBLUP and sERRBLUP by using the combined phenotypes of the other five environments as the additional environment instead of a single environment’s phenotypes which are then termed ‘bivariate GBLUP across multiple environments jointly’, ‘bivariate ERRBLUP across multiple environments jointly’ and ‘bivariate sERRBLUP across multiple environments jointly’.

### Estimation of variance and covariance components

Since we aimed at estimating variance components in each replicate of the cross-validation from the training data, but variance component estimation with ASREML has a certain risk of non-convergence, we needed to specify a strategy to deal with such cases in an automated manner, given the huge number of analyses we ran. In univariate analyses, variance components were estimated using EMMREML (Akdemir and Godfrey, 2015) in each run of a 5-fold cross validation based on the training set. In bivariate analyses, the variance components were estimated using ASReml-R (Butler *et al*., 2018). In the bivariate ERRBLUP and sERRBLUP models, the genetic and residual variance and covariance were estimated first from the full data set in a bivariate ASReml-R model for each combination of environments in each trait. If the estimation of variance components didn’t converge after 100 iterations, then the computation was stopped and the genetic and residual variance and covariance estimates at the last iteration (100) were extracted. These estimates were defined as the initial starting values of the bivariate ASReml-R model in each run of a 5-fold cross validation, followed by a re-estimation of the variance and covariance components based on the training set in the cross validation. If the estimation of variance components did not converge at 50 iterations in each fold, the pre-estimated variance and covariance components based on the full dataset, which was defined as the initial start values of the model, were used as fixed values, so that the breeding values were estimated based on these pre-estimated parameters. It was verified from converged estimates that variance and covariance components estimated from the training set deviated only little from the variances and covariances from the full set (see Fig. 3). Also, the mean result obtained from just the converged replicates and the mean results of all replicates including the ones where variance and covariance components were fixed were rather similar (Fig. 4), only when the majority (>20) of replicates failed to converge, substantial random fluctuation was observed. Thus we argue that this strategy appears justifiable, but still the number of cases where estimates did not converge in 5-fold cross validation with 5 replicates and the combinations whose pre estimation of variance components also did not converged in 100 iterations are detailed in the supplementary (Table S2 – S9).

**Fig. 3.**
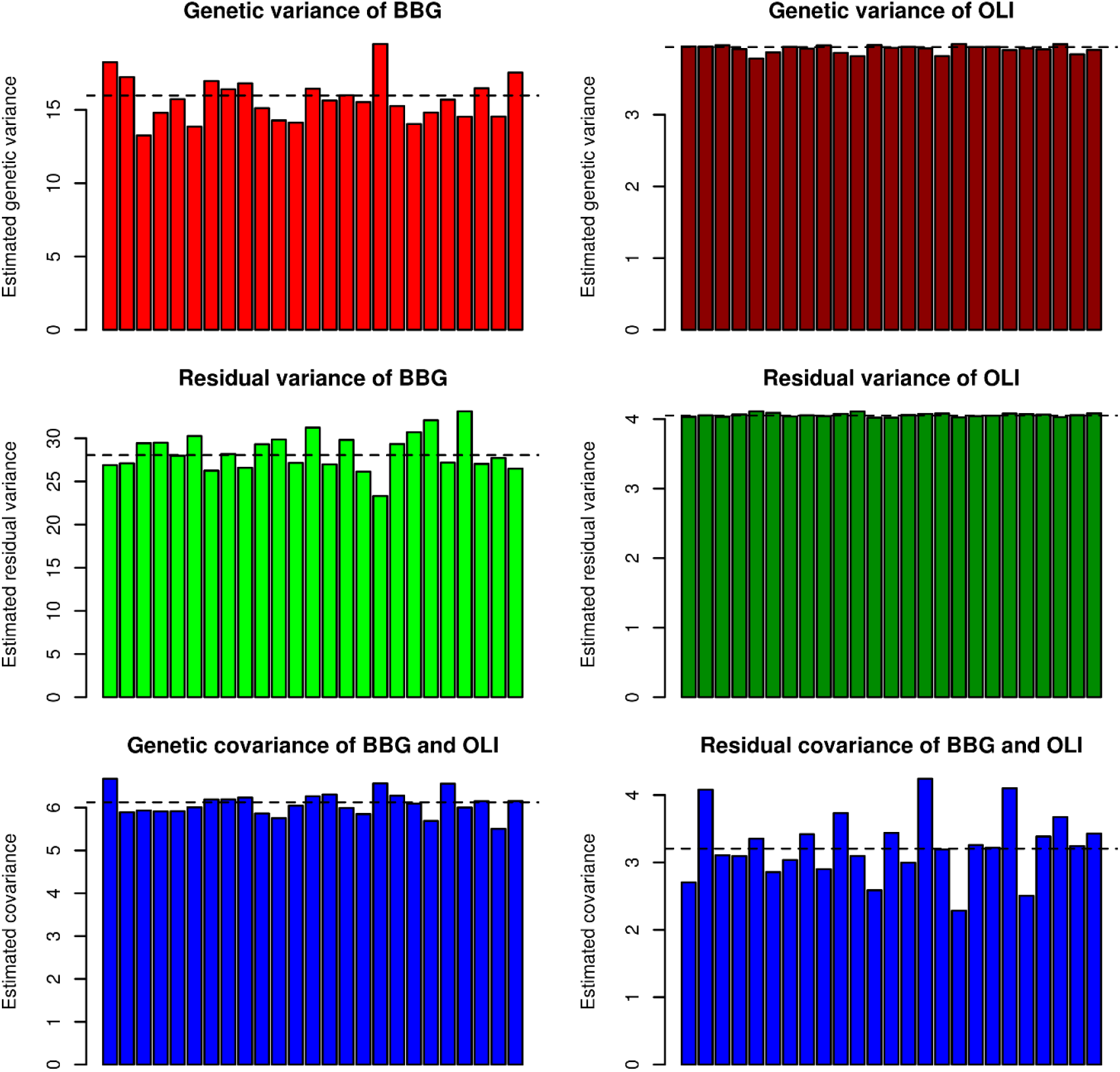
Comparison of pre estimated genetic and residual variances and covariances of converged bivariate sERRBLUP model (top 5%) based on the full dataset (dashed horizontal lines) and estimated genetic and residual variances and covariances of converged bivariate sERRBLUP (top 5%) based on training set in each run of 5-fold cross validation with 5 replicates (colored bars) for predicting BBG when the additional environment is OLI in Kemater for trait PH-V4.

**Fig. 4.**
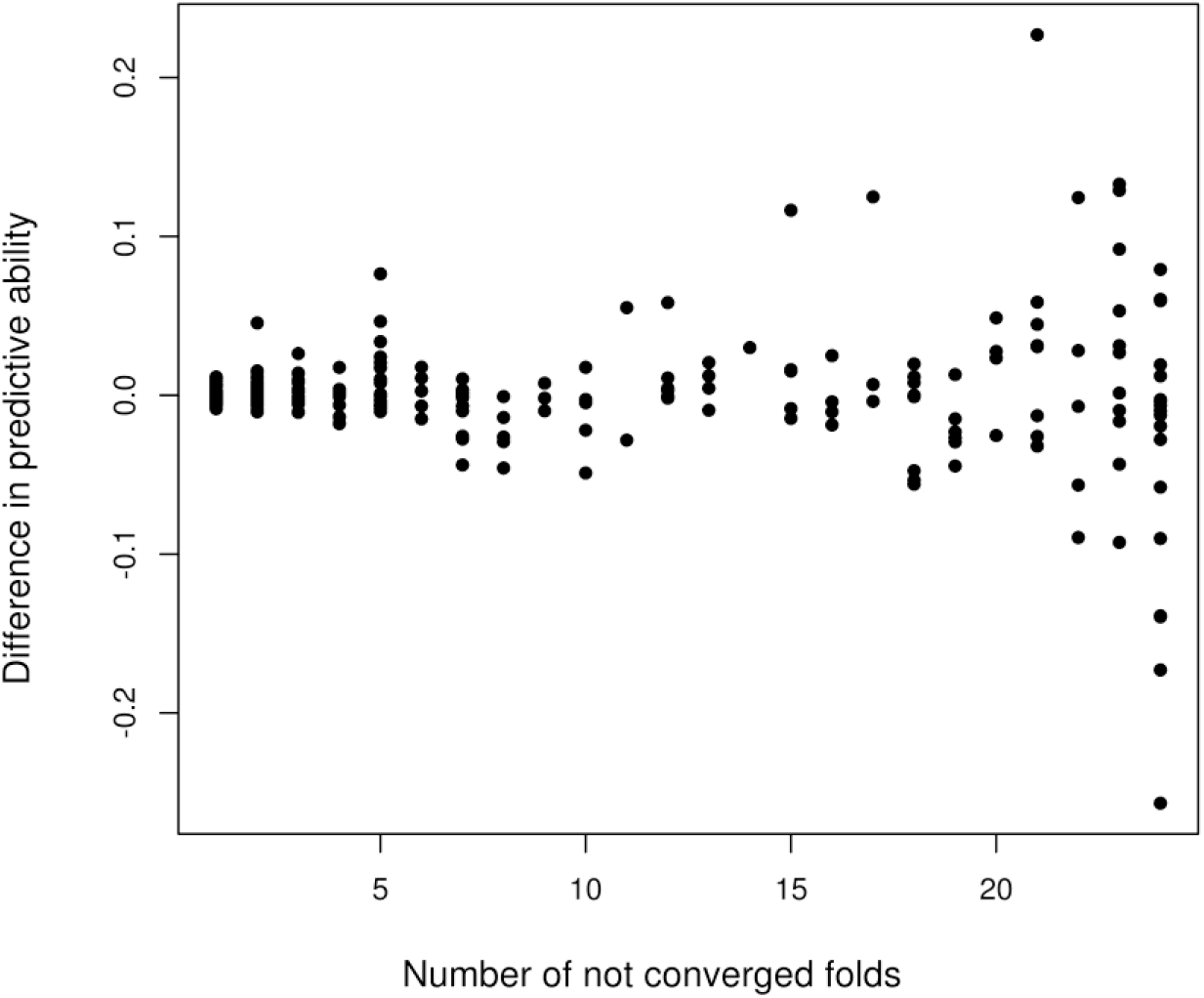
The difference between the mean predictive ability of only the converged folds and the mean predictive ability of all folds in 5-fold cross validation with 5 replicates vs. the number of the folds (1 to 24) which did not converge across all traits in all combinations for both Kemater and Petkuser.

## Results

Predictive abilities of univariate sERRBLUP across environments compared to univariate ERRBLUP and univariate GBLUP within environments for the trait PH_V4 are shown in Fig. 5 and 6 for KE and PE, respectively. Univariate GBLUP within the environment is used as a reference and is compared to results obtained with univariate ERRBLUP within environments and univariate sERRBLUP when the top 5, 1, 0.1, 0.01, 0.001 and 0.0001 percent of pairwise SNP interactions are maintained in the model. Fig. 5 and 6 show that the predictive abilities of univariate GBLUP and univariate ERRBLUP within the environment are almost identical (the highest deviation observed was 0.004). A considerable increase in predictive ability was observed when the top 1 or 0.1 percent of SNP interactions were kept in the univariate sERRBLUP model regardless of the interaction selection strategy. A more stringent selection, i.e. by considering only the top 0.01, 0.001 and 0.0001 percent of SNP interactions in the model, often led to a reduction in predictive ability. For the most stringent selection of 0.001 and 0.0001 percent, the predictive ability was sometimes even below the univariate GBLUP reference. This pattern is observed over all environments which is more pronounced in KE then PE. Moreover, the predictive abilities of sERRBLUP when the selection of pairwise SNP interaction was based on estimated absolute effect sizes (left hand side of the panel) were compared to the predictive abilities of sERRBLUP when SNP interactions were selected based on estimated effects variances (right hand side of the panel). This comparison reveals that the interaction selection based on the effect variances is more robust than interaction selection based on the absolute effect sizes, especially when the top 0.001 percent of interactions are maintained in the model for KE. However, this comparison in Fig. 6 does not display a considerable difference for the interaction selection strategy for PE. Due to the higher robustness of the approach based on interaction variances as selection criterion for KE, we used this strategy for the series of all other traits in both landraces, for which results are shown in the supplementary (Fig. S1a – S7a).

**Fig. 5.**
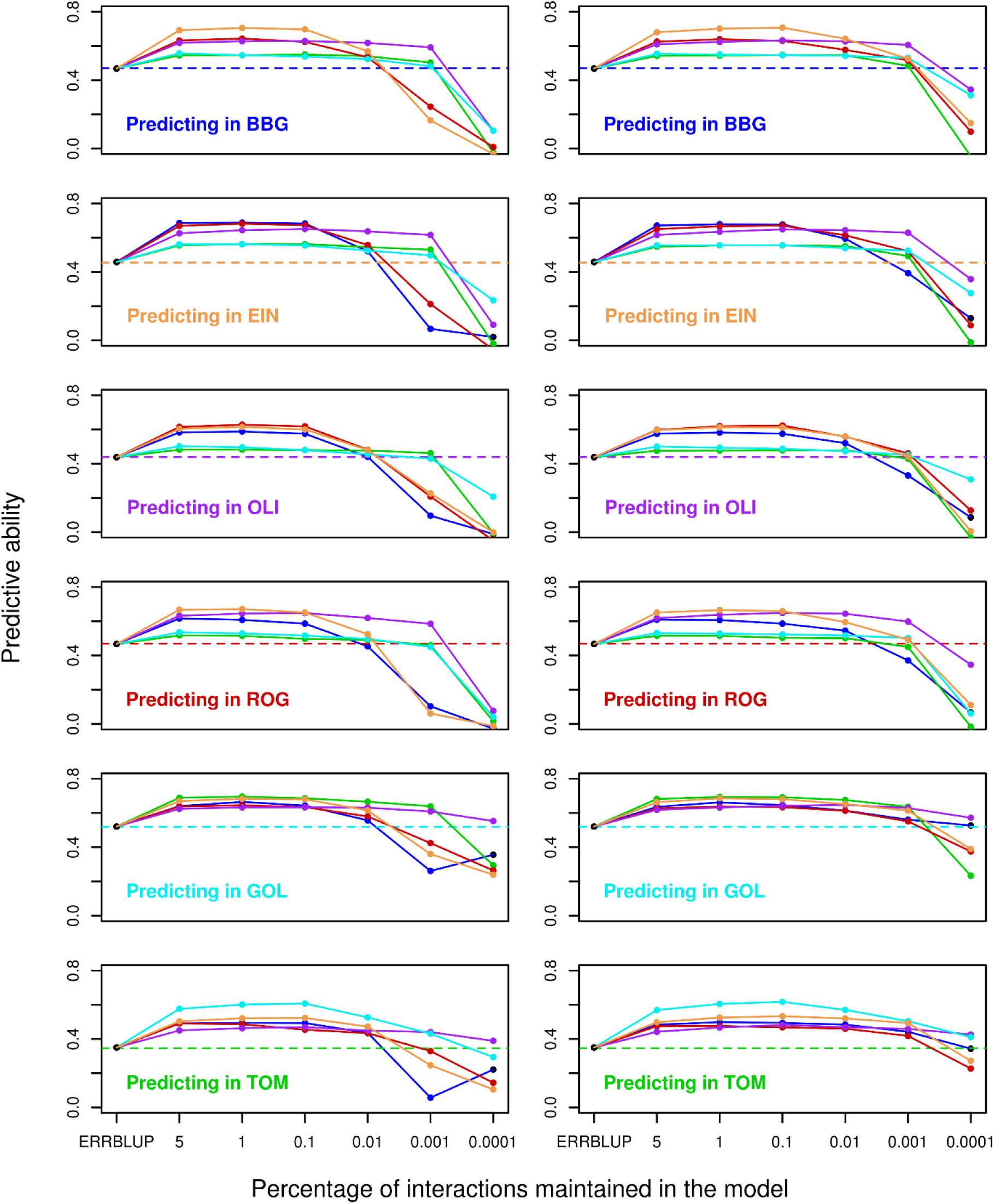
Predictive ability for univariate GBLUP within environment (dashed horizontal line), univariate ERRBLUP within environment (black filled circle) and univariate sERRBLUP across environments (solid colored lines) when SNP interaction selections are based on estimated effects sizes (left side) and estimated effects variances (right side) for trait PH-V4 in Kemater. In each panel, the solid lines’ color indicates the environment in which the relationship matrices were determined by variable selection.

**Fig. 6.**
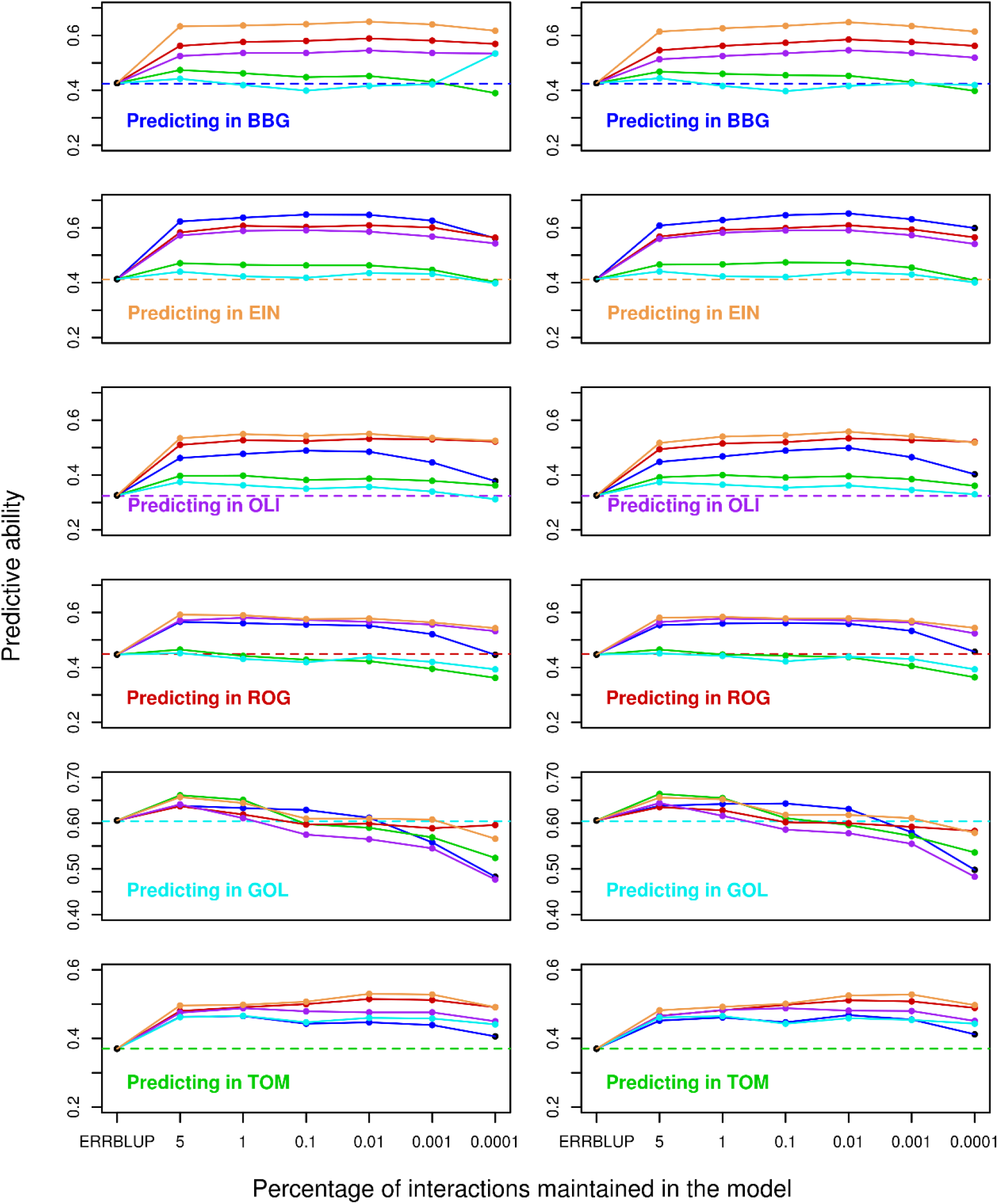
Predictive ability for univariate GBLUP within environment (dashed horizontal line), univariate ERRBLUP within environment (black filled circle) and univariate sERRBLUP across environments (solid colored lines) when SNP interaction selections are based on estimated effects sizes (left side) and estimated effects variances (right side) for trait PH-V4 in Petkuser. In each panel, the solid lines’ color indicates the environment in which the relationship matrices were determined by variable selection.

In the context of univariate models, we also investigated the predictive ability of univariate sERRBLUP when variable selection was based on the training set of the same environment as the test set was sampled from. This was exemplarily done within Bernburg for the trait PH-V4 (see Fig. S8), illustrating that the predictive ability obtained from univariate sERRBLUP is marginally higher than univariate GBLUP only when the top 0.01 percent of interactions are kept in the model. When selection of effects is too strict, with only 0.001 percent of interactions used, predictive ability of univariate sERRBLUP within Bernburg falls short of the one obtained with GBLUP, especially if selection is based on effect sizes. Consequently, we did not include this model variant in our model comparison due to its poor performance which was obtained at a cost of high computing time, required for setting up a different univariate sERRBLUP relationship matrix for each training set.

The predictive abilities of bivariate GBLUP, ERRBLUP and sERRBLUP when SNP interactions were selected based on estimated effect variances are compared for trait PH_V4 in KE and PE in Fig. 7. Fig. 7 shows that the bivariate ERRBLUP increases the predictive ability slightly compared to bivariate GBLUP with the maximum absolute increase of 0.03 in KE and 0.02 in PE across all environments’ combinations. A considerable increase in predictive ability is obtained in bivariate sERRBLUP mostly when the top 5 or 1 percent of interactions are maintained in the model. However, the bivariate sERRBLUP predictive abilities decrease dramatically for too stringent selection of pairwise SNP interactions such as 0.01, 0.001 or 0.0001 percent. Moreover, the reduction in predictive ability with too stringent factor selection is more severe for KE than for PE. This pattern is observed for the majority of environments for both landraces and the results for other traits are shown in the supplementary (Fig. S1b – S7b).

**Fig. 7.**
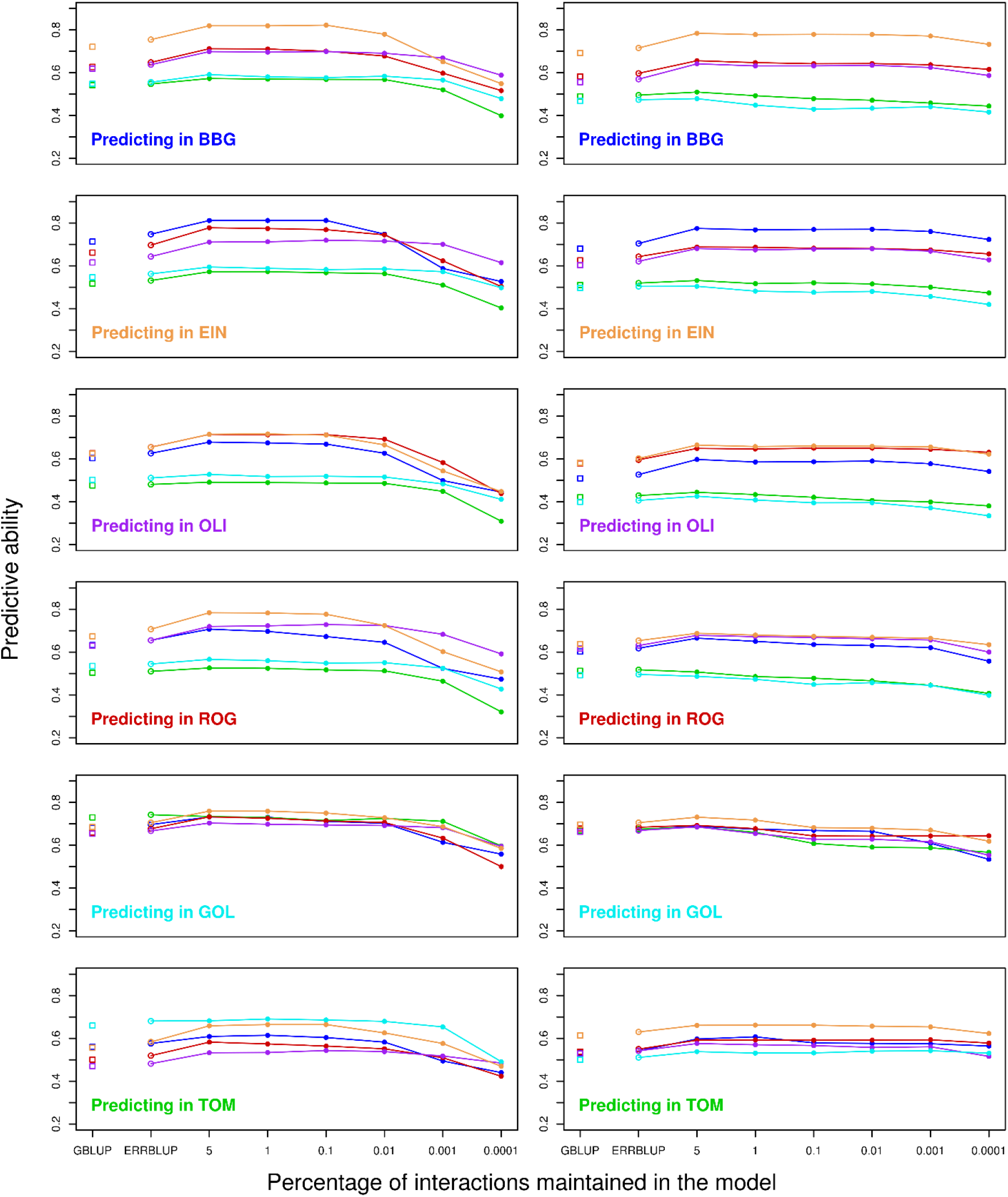
Predictive ability for bivariate GBLUP (open squares), bivariate ERRBLUP (open circles) and bivariate sERRBLUP (filled circles and solid lines) when SNP interaction selections are based on estimated effects variances in Kemater (left side) and Petkuser (right side) for trait PH-V4. In each panel, the solid lines’ color indicates the additional environment used to predict the target environment.

Fig. 8 provides a comparison of predictive ability obtained from univariate GBLUP within environments and the maximum predictive ability obtained from univariate sERRBLUP across environments based on both strategies of interaction selection for trait PH_V4. Both panels in Fig. 8 demonstrate that for both landraces in each environment univariate sERRBLUP across environments outperforms univariate GBLUP regardless of the selection strategy. It also demonstrates that the maximum predictive ability obtained when selecting interactions based on absolute effect sizes and their variances are almost identical.

**Fig. 8.**
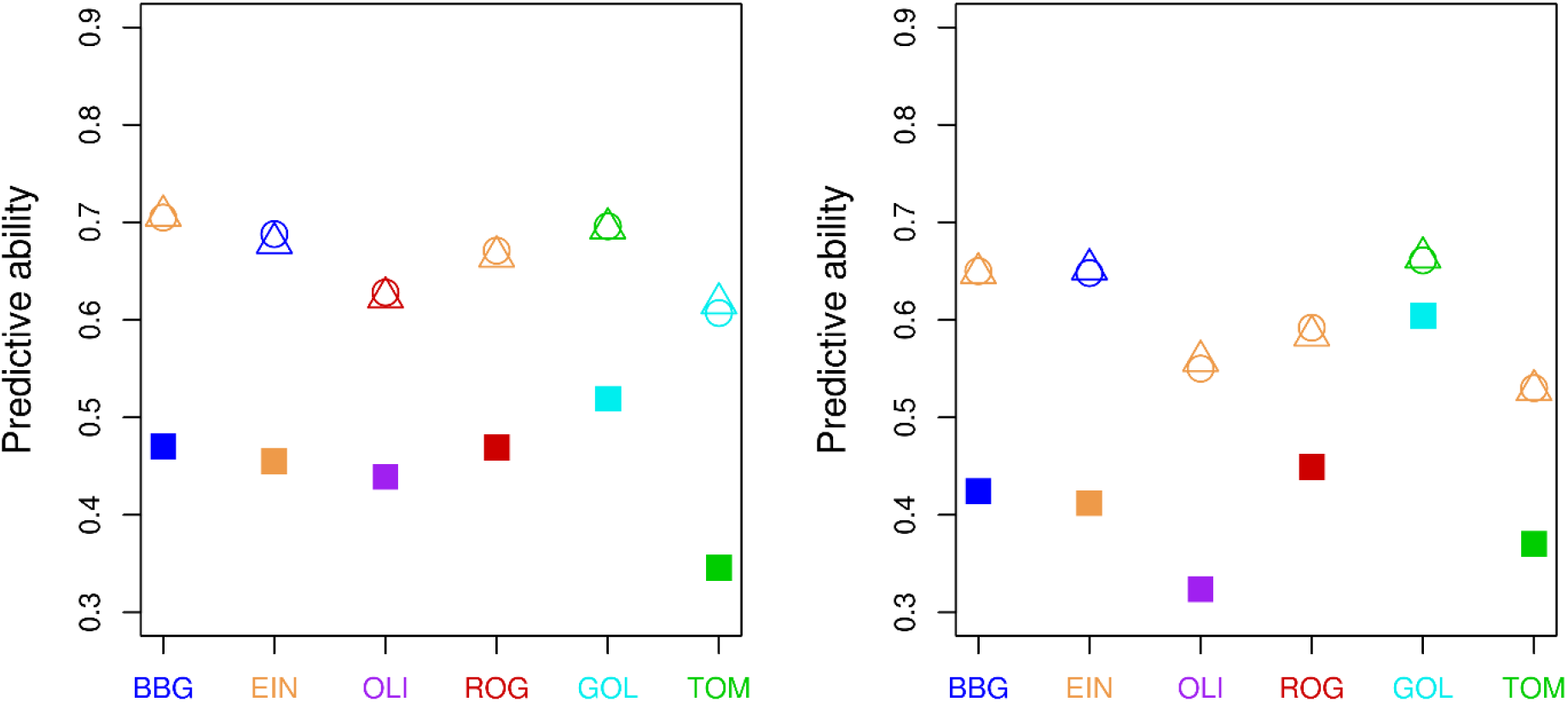
Comparison of predictive ability of univariate GBLUP within environments (filled squares) and the maximum predictive ability of univariate sERRBLUP across environments when the SNP interaction selections are based on estimated effects sizes (circles) and estimated effects variances (triangles) for trait PH-V4 in Kemater (left side plot) and in Petkuser (right side plot). The colors dark blue, green, red, purple, light blue and orange represent the environments BBG, EIN, OLI, ROG, GOL and TOM, respectively. The circles’ and triangles’ colors indicate the environment which had the maximum predictive ability for this respective target environment

The maximum predictive ability obtained from bivariate GBLUP is also compared with the maximum predictive ability obtained from bivariate sERRBLUP when pairwise SNP interaction selections were based on estimated effects variances in Fig. 9 for the trait PH_V4. Both panels of Fig. 9 illustrate that the predictive ability in each environment is enhanced from bivariate GBLUP to bivariate sERRBLUP in both landraces.

**Fig. 9.**
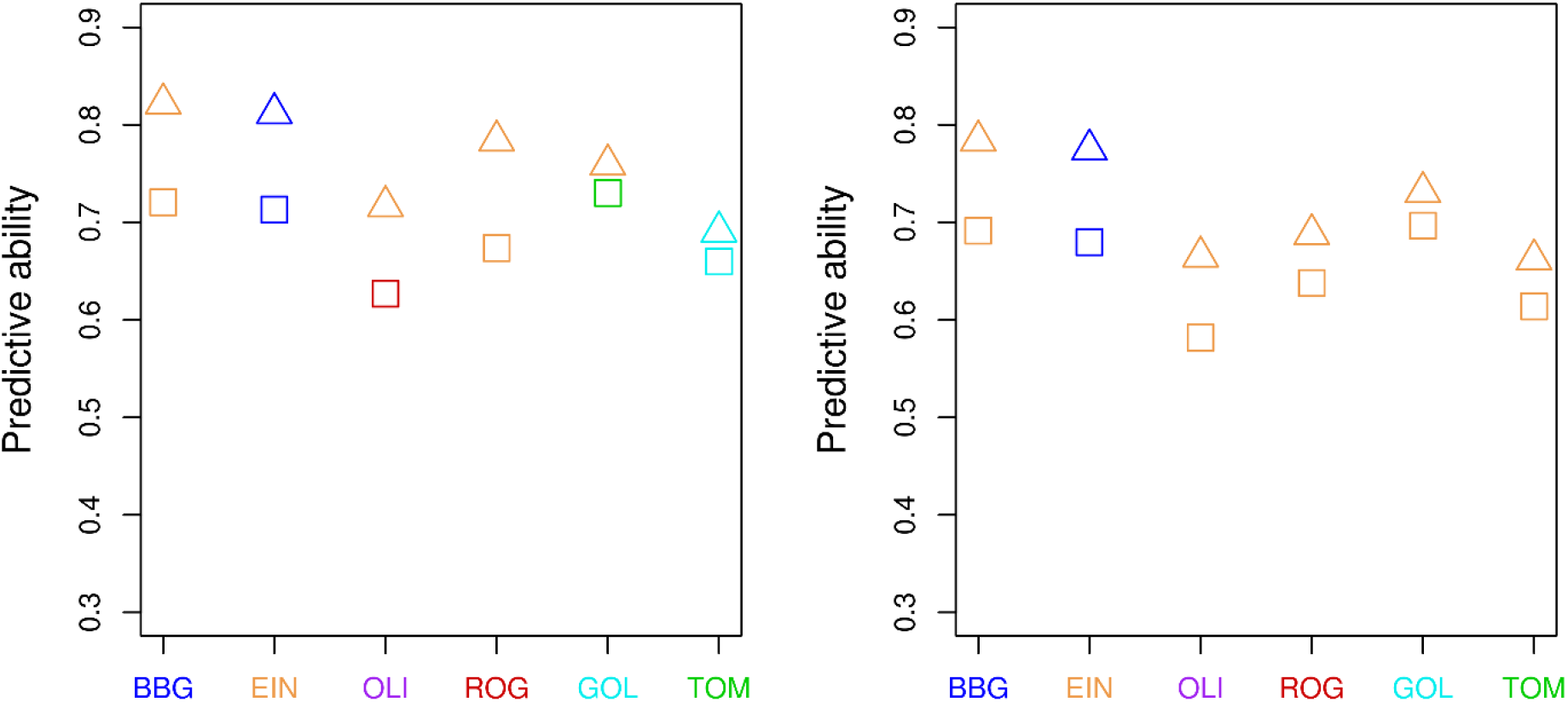
Comparison of maximum predictive ability of bivariate GBLUP (squares) to the maximum predictive ability of bivariate sERRBLUP (triangles) when the SNP interactions are selected based on estimated effects variances for trait PH-V4 in Kemater (left side plot) and in Petkuser (right side plot). The colors dark blue, green, red, purple, light blue and orange represent the environments BBG, EIN, OLI, ROG, GOL and TOM, respectively. The colors of the squares and triangles indicate the environment leading to the maximal predictive ability for this respective target environment.

The relative increase in prediction accuracy of the best univariate sERRBLUP across environments compared to univariate GBLUP within environments for all traits and all locations is shown in form of a heat map in Fig. 10 for both landraces. The maximum relative increase in prediction accuracy among all traits and all environments in KE is 85.6 percent (PH_V6 in OLI) and in PE it is 112.4 percent (EV_V3 in EIN). Those highest increases in accuracy were found in traits and environments combinations where the univariate GBLUP prediction accuracy was particularly low. An increase is observed in each studied trait by location combination, with the smallest increase in both landraces for PH_final in BBG (20.1 percent in KE) or in GOL (5.9 percent in PE). In general, both plots in Fig. 10 demonstrate that for the majority of traits and environments, there is more than 30 percent increase in prediction accuracy from univariate GBLUP within environments to the best univariate sERRBLUP across environments. Average increase across all combinations in KE is 47.1 percent and in PE is 46.7 percent. Fig. 10 also shows the average increase in prediction accuracy for each environment across series of all traits and for each trait across series of all environments. The maximum average increase in prediction accuracy among all traits was found for EV_V4 in KE (59.7 percent) and EV_V3 in PE (75.0 percent). Regarding environments, the maximum average increase in prediction accuracy obtained for univariate sERRBLUP was in TOM for KE (63.3 percent) and in OLI for PE (71.9 percent). Additionally, the absolute increase in prediction accuracy is shown as a heat map in supplementary (Fig. S9a) which indicates the average absolute increase of 0.204 in KE and 0.181 in PE.

**Fig. 10.**
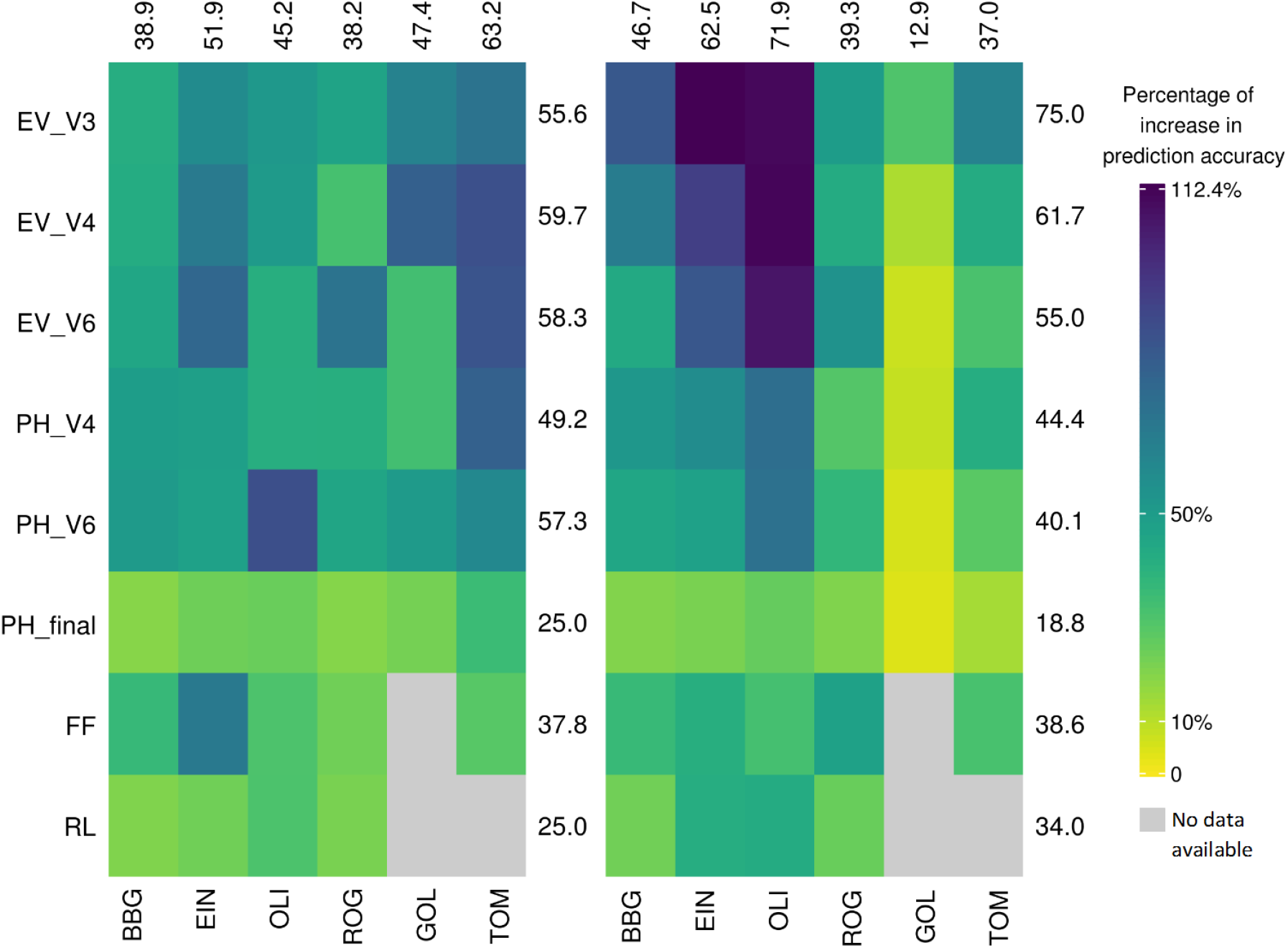
Percentage of increase in prediction accuracy from univariate GBLUP within environments to the maximum prediction accuracy of univariate sERRBLUP across environments when the SNP interaction selections are based on estimated effects variances in Kemater (left side plot) and in Petkuser (right side plot). The average percentage of increase in prediction accuracy for each trait and environments are displayed in rows and columns, respectively.

Fig. 11 also shows the relative increase in prediction accuracy from the best bivariate GBLUP to the best bivariate sERRBLUP for all traits and all locations in a form of heat map. The maximum increase in prediction accuracy among all traits and all environments is 21.1 percent (EV_V6 in ROG) in KE and 27.9 percent (EV_V3 in BBG) in PE. There is an increase across all studied traits in all environments except for the trait PH_final in PE which shows a relative decrease of 0.257 percent. The minimum increase in prediction accuracy in KE was also observed for PH_final (1.7 percent). In general, Fig. 11 shows that the relative increase in prediction accuracy from the best bivariate GBLUP to the best bivariate sERRBLUP is more than 7 percent for the majority of trait by location combinations in both landraces with an average increase of 10.9 percent in KE and 10.5 in PE across all combinations. The maximum average increase in prediction accuracy among all traits was observed for EV_V3 in both landraces (16.3 percent in KE and 20.3 percent in PE). Regarding environments, the maximum average increase across all environments was found in EIN for KE (13.4 percent) and in OLI for PE (14.6 percent). In addition, the absolute increase in prediction accuracy of bivariate models is shown as a heat map in supplementary (Fig. S9b) indicating an average absolute increase of 0.1 across all traits, environments combinations, and landraces.

**Fig. 11.**
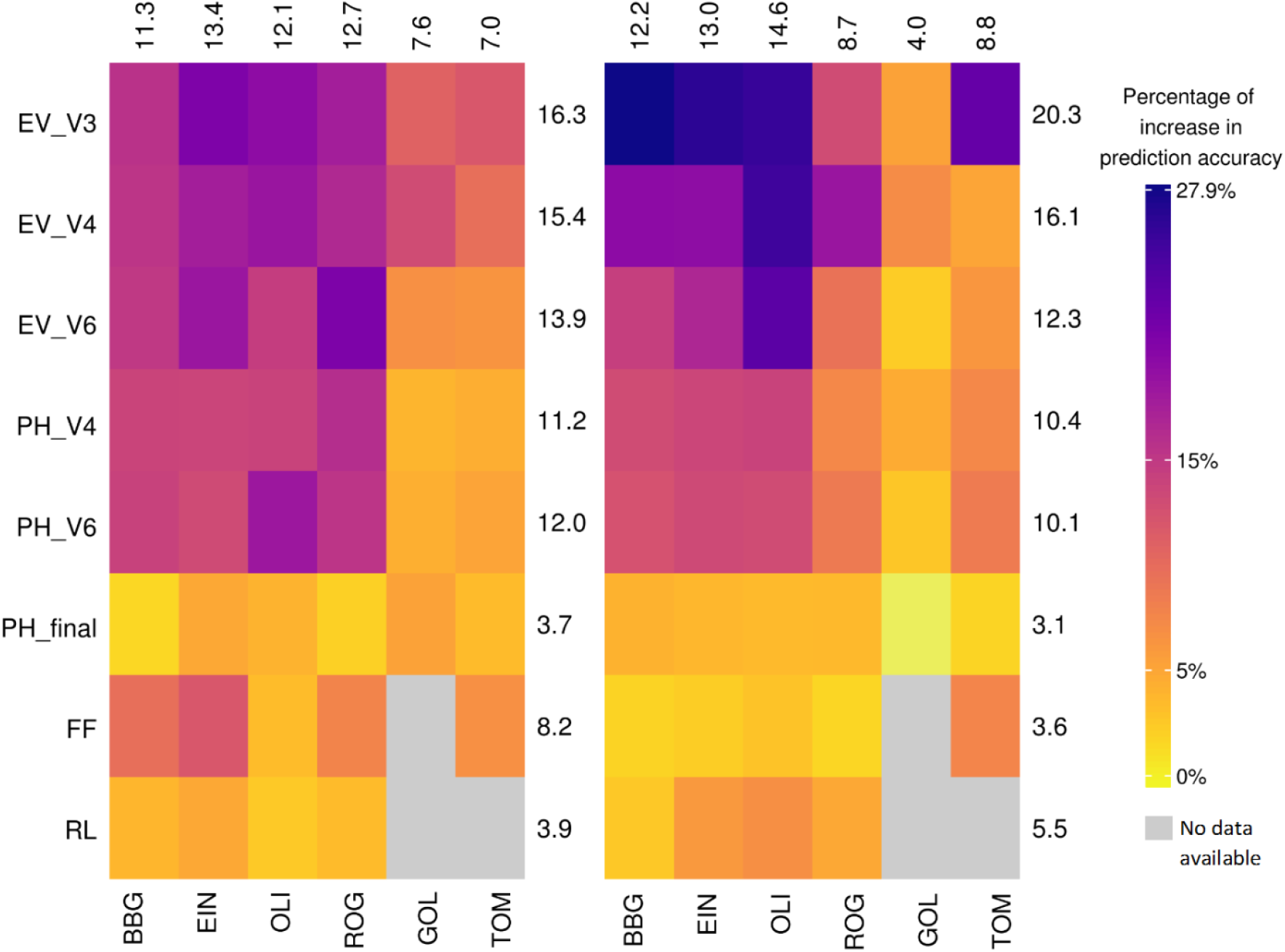
Percentage of increase in prediction accuracy from the maximum bivariate GBLUP to the maximum prediction accuracy of bivariate sERRBLUP when the SNP interaction selections are based on estimated effects variances in Kemater (left side plot) and in Petkuser (right side plot). The average percentage of increase in prediction accuracy for each trait and environments are displayed in rows and columns, respectively.

In general, we assume that predictive ability for phenotypes should be higher with higher heritability. This is confirmed by the results in both Fig. 12 and 13, depicting the correlation between the heritability of all traits and the predictive ability obtained from univariate and bivariate models, respectively. The traits’ heritabilities were calculated on an entry-mean basis within each KE and PE landraces (Hallauer *et al*., 2010) over all six environments in the year 2017. Fig. 12 shows the correlation between the heritability of all traits and the univariate GBLUP within environments and maximum predictive ability of univariate sERRBLUP across environments. It is shown that this correlation increased from 0.296 with the univariate GBLUP model to 0.543 with the univariate sERRBLUP model. Likewise, Fig. 13 shows the correlation between the heritability of all traits and the maximum predictive ability of bivariate GBLUP and maximum predictive ability of bivariate sERRBLUP. It is shown that this correlation in bivariate GBLUP is 0.537 which is lower than respective correlation in univariate sERRBLUP and it is increased to 0.647 in bivariate sERRBLUP.

**Fig. 12.**
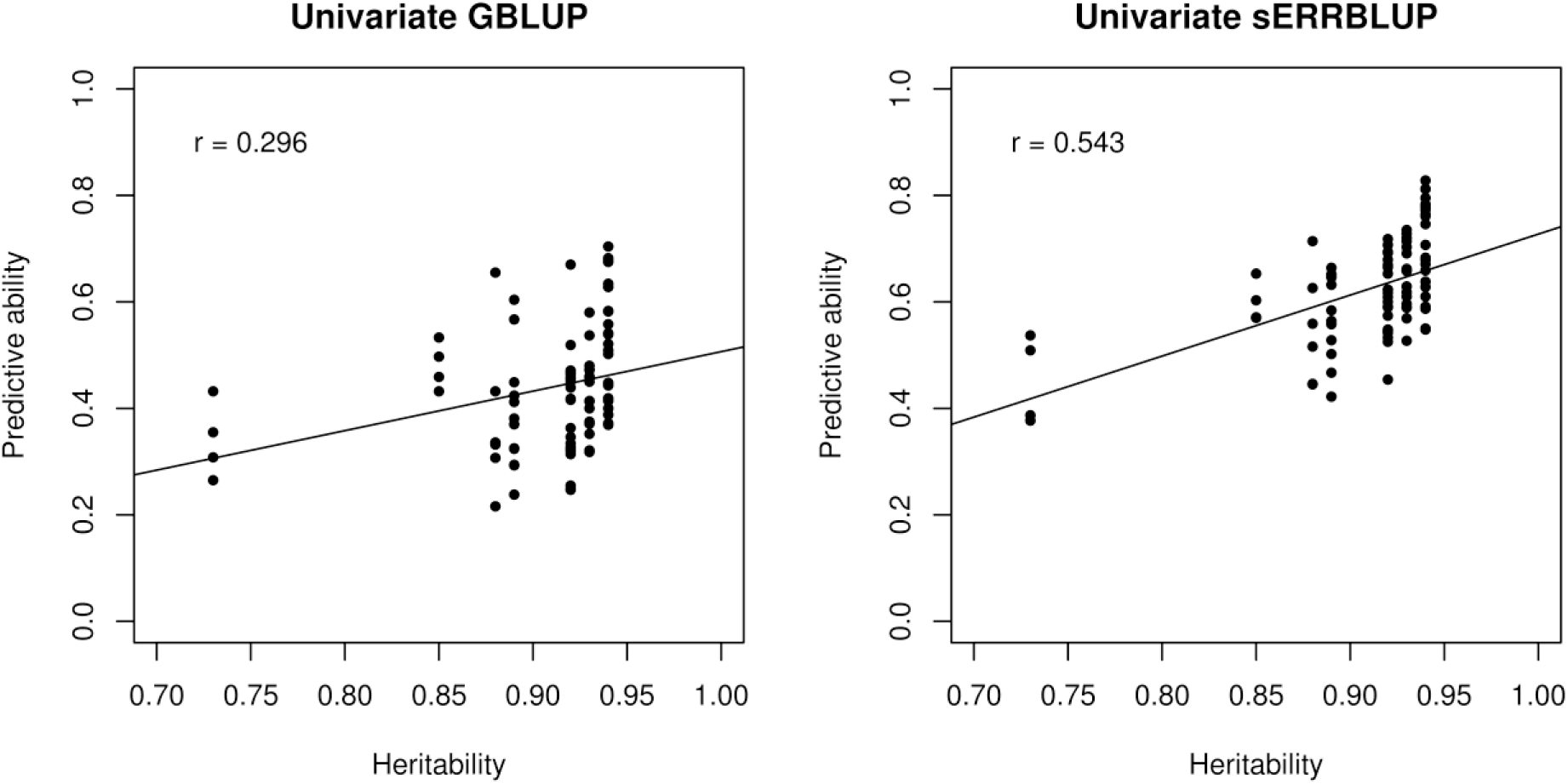
The correlation between all eight traits’ heritabilities and predictive abilities of univariate GBLUP within all six environments (left side plot) and maximum predictive abilities of univariate sERRBLUP across environments (right side plot) for all traits in both landraces. The black lines indicate the overall linear regression lines.

**Fig. 13.**
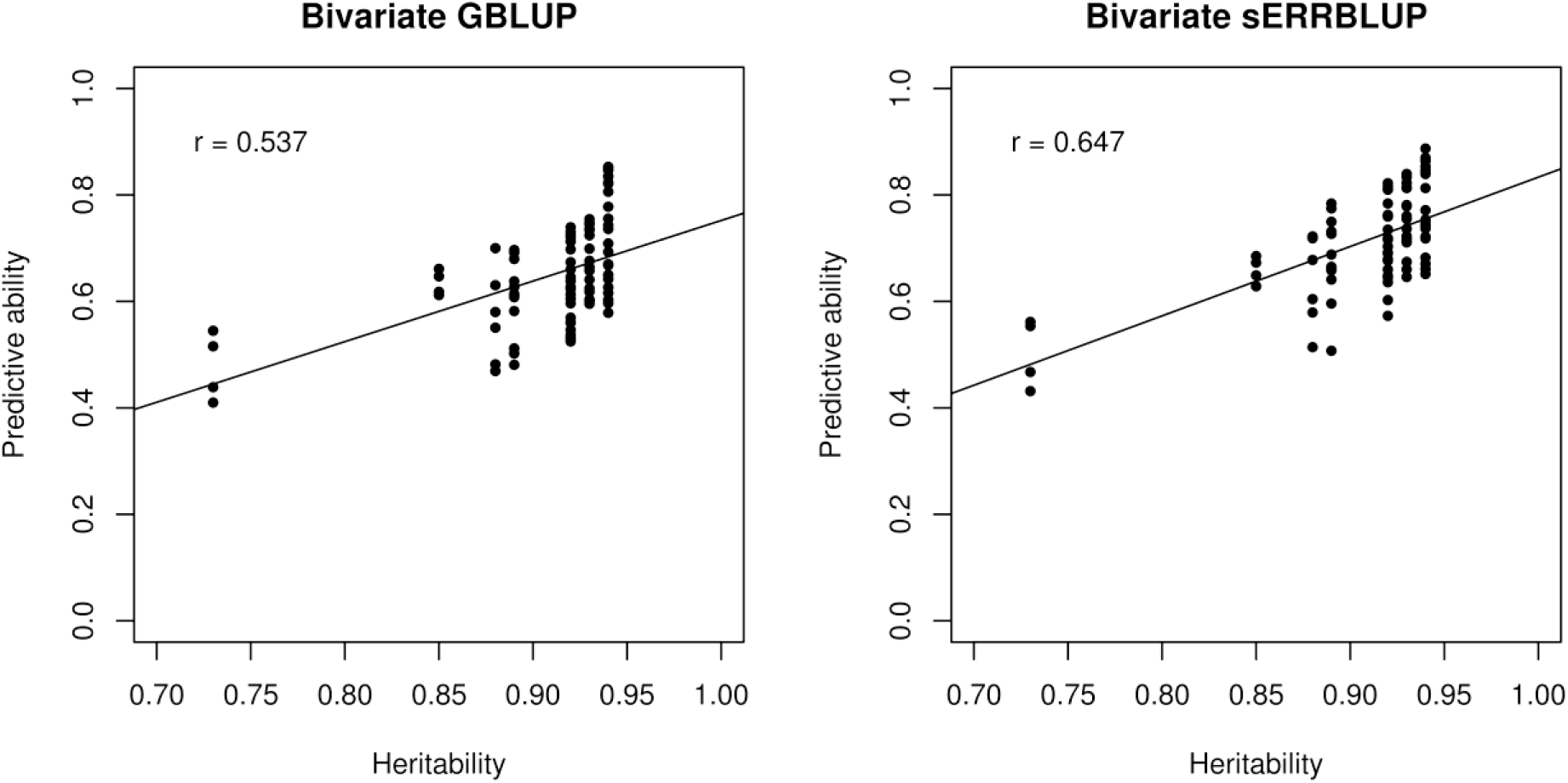
The correlation between all eight traits’ heritabilities and the maximum predictive abilities of bivariate GBLUP (left side plot) and maximum predictive abilities of bivariate sERRBLUP (right side plot) for all six environments in both landraces. The black lines indicate the overall linear regression lines.

In addition to assessing the predictive ability of univariate sERRBLUP based on a single environment, Fig. 14 and 15 display the comparison between the predictive ability obtained from univariate GBLUP and univariate ERRBLUP within environments, and univariate sERRBLUP across multiple environments jointly for trait PH-V4 in KE and PE, respectively. In each Figure, the left hand side of the panel represent predictive abilities of univariate sERRBLUP across multiple environments jointly when pairwise SNP interaction selections were based on estimated absolute effect sizes, and the right hand side of the panel represent predictive abilities of univariate sERRBLUP across multiple environments jointly when pairwise SNP interaction selections were based on estimated effects variances. It is demonstrated that univariate sERRBLUP has a higher predictive ability than univariate GBLUP when interactions are selected based on all the other five environments jointly. Fig. 14 also reveals the robustness of the selection strategy based on the effects variance compared to selection strategy based on the absolute effects sizes in KE, while Fig. 15 does not show a significant difference for the interaction selection strategy for PE. Fig. 14 and 15 demonstrate that the predictive ability of univariate sERRBLUP across multiple environments jointly is as good as or better than using a single environment with few exceptions when selection of effects is not too strict. With less than 0.1 percent of interactions used, predictive abilities deteriorate (especially so in KE) and selection from combined environments turns out to be worse than selection from single environments. The comparison between the maximum predictive ability of univariate sERRBLUP across a single environment and multiple environments jointly are shown in the supplementary (Fig. S10).

**Fig. 14.**
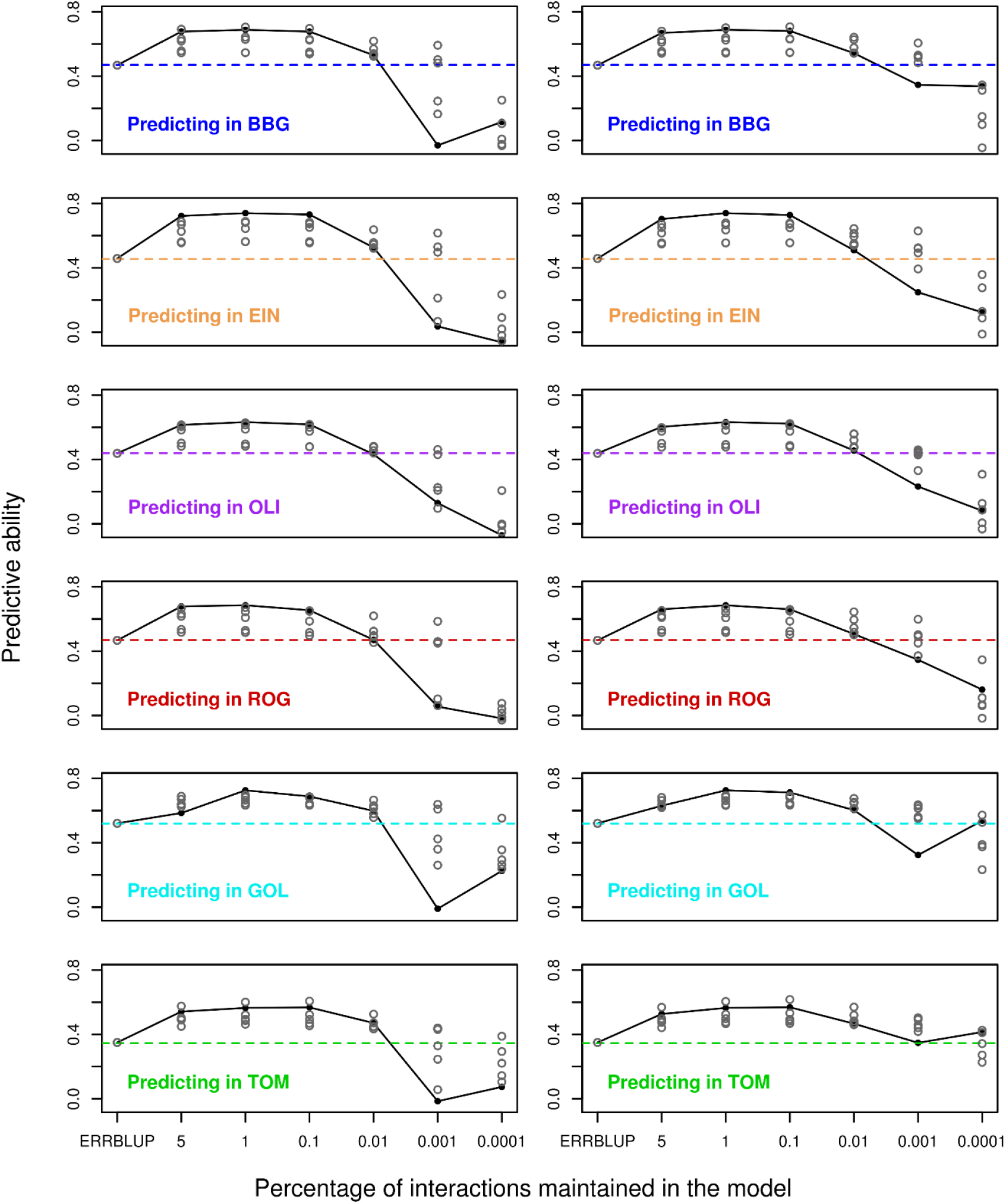
Predictive ability for univariate GBLUP within environment (dashed horizontal line), univariate ERRBLUP within environment (gray open circle), univariate sERRBLUP using a single environment for selecting the SNP interactions (gray open circles) and univariate sERRBLUP using all 5 environments jointly (filled black circles and solid line) for the SNP interaction selection based on estimated effects sizes (left side) and estimated effects variances (right side) for trait PH-V4 in Kemater.

**Fig. 15.**
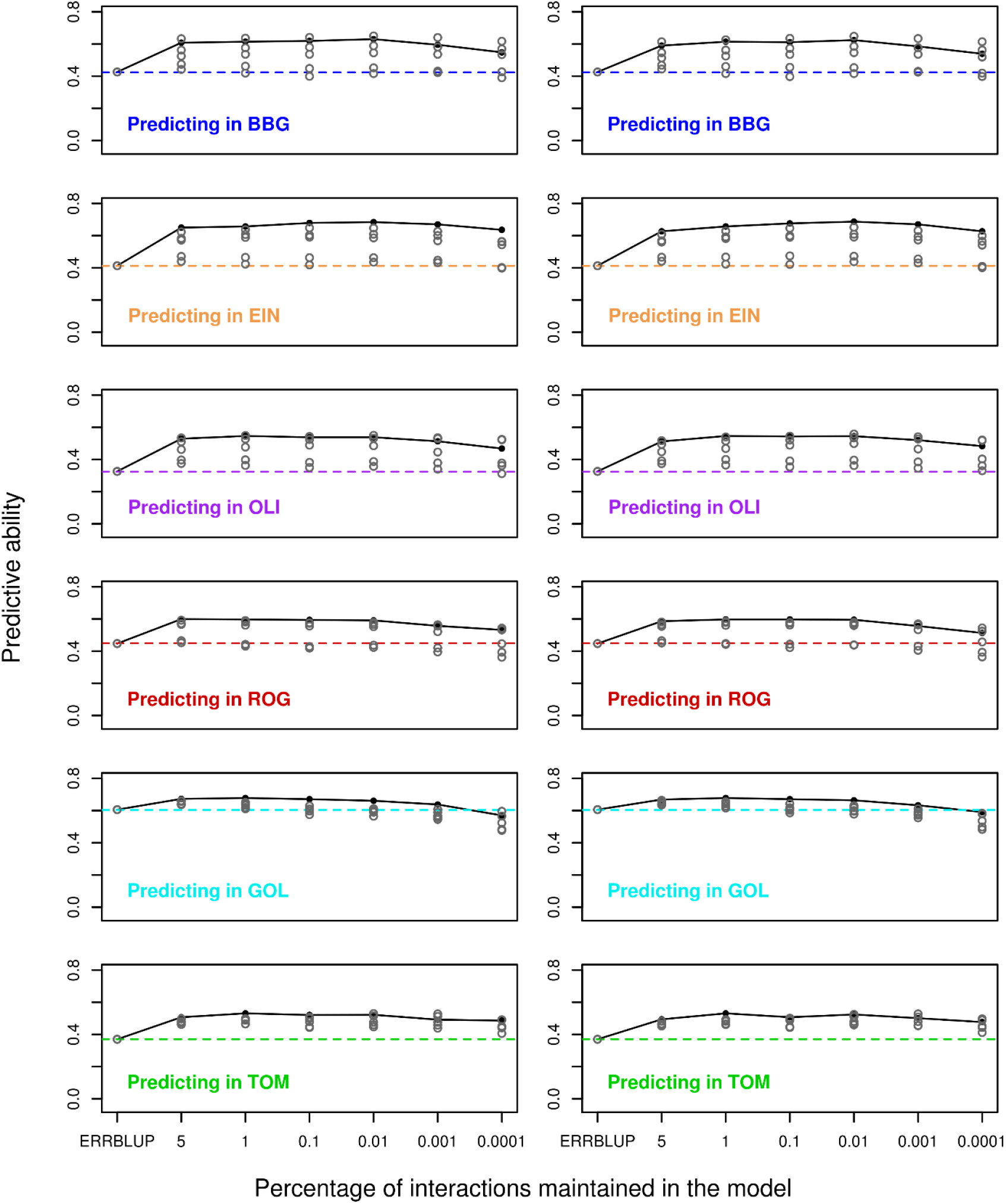
Predictive ability for univariate GBLUP within environment (dashed horizontal line), univariate ERRBLUP within environment (gray open circle), univariate sERRBLUP using a single environment for selecting the SNP interactions (gray open circles) and univariate sERRBLUP using all 5 environments jointly (filled black circles and solid line) for the SNP interaction selection based on estimated effects sizes (left side) and estimated effects variances (right side) for trait PH-V4 in Petkuser.

Fig. 16 illustrates the comparison between the predictive ability of bivariate GBLUP, ERRBLUP and sERRBLUP across multiple environments jointly and the maximum predictive ability of bivariate GBLUP and ERRBLUP and all the predicative abilities of sERRBLUP when a single environment is considered as an additional environment for the trait PH_V4 in both KE and PE. The results indicate that bivariate sERRBLUP across multiple environments jointly increases the predictive ability compared to bivariate GBLUP and ERRBLUP across multiple environments jointly. Moreover, in most cases bivariate GBLUP, ERRBLUP and sERRBLUP across multiple environments jointly are as good as or better than bivariate GBLUP, ERRBLUP and sERRBLUP (when the selection of effects is not too strict especially in KE) joint with a single environment, respectively. With less than 0.1 percent of pairwise SNP interactions used in the model, the predictive ability of the bivariate sERRBLUP across multiple environments jointly decreases considerably in KE.

**Fig. 16.**
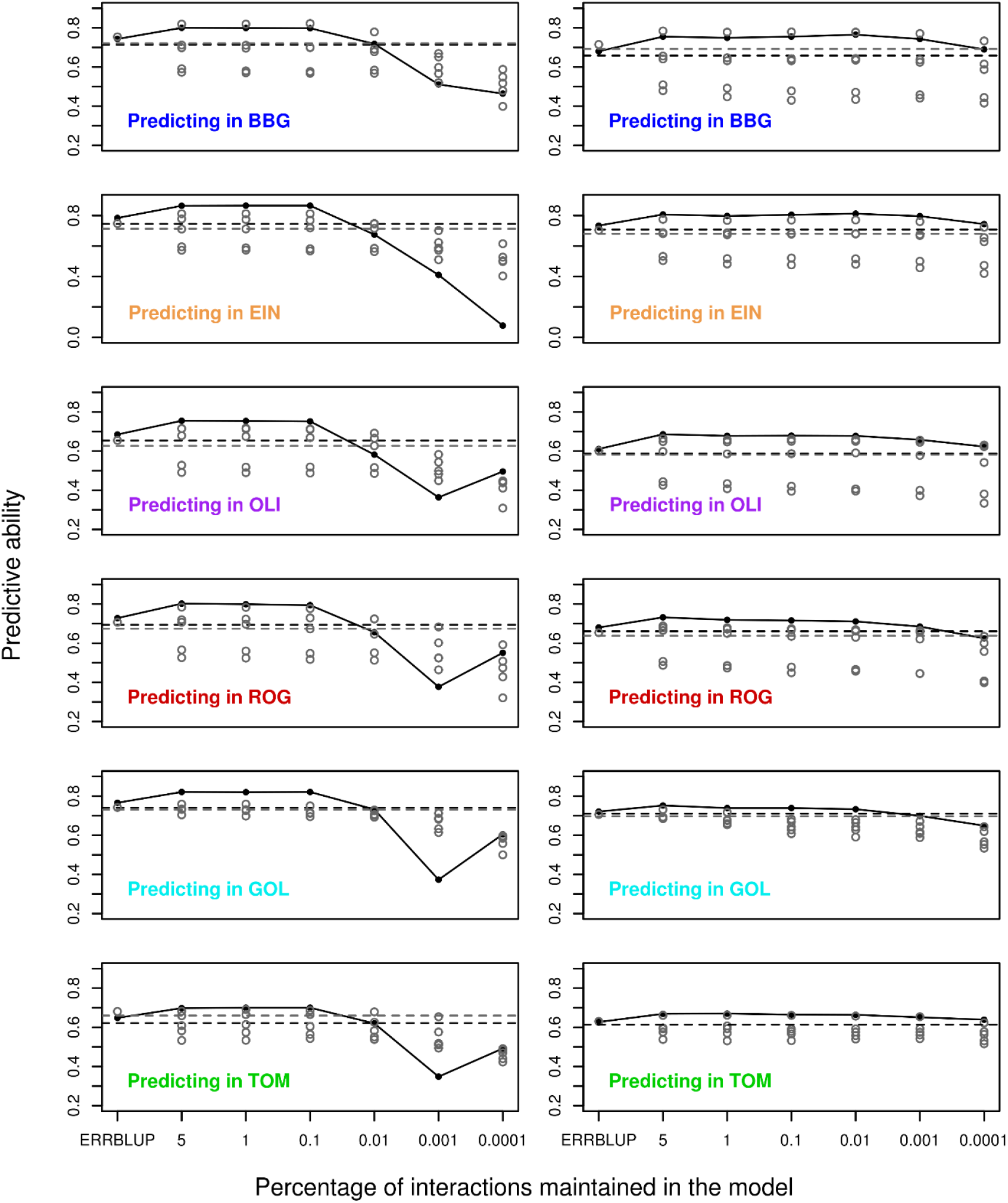
Predictive ability for bivariate GBLUP (black dashed horizontal line), bivariate ERRBLUP and bivariate sERRBLUP (filled black circles) for the SNP interaction selection based on estimated effects variances using all 5 environments jointly for trait PH-V4 in Kemater (left side) and Petkuser (right side). In each panel, gray horizontal line and first gray open circles refer to maximum bivariate GBLUP and maximum bivariate ERRBLUP, and the gray open circles at the top 5, 1, 0.1, 0.01, 0.001, 0.0001 quantiles refer to bivariate sERRBLUP using a single environment as an additional environment.

## Discussion

The accuracy of the genomic prediction when incorporating epistasis interactions in the model compared to prediction models with only main effects have been widely discussed over the last years. In particular, it was found that accounting for epistasis can increase predictive ability (Carlborg and Haley, 2004; Hu *et al*., 2011; Huang *et al*., 2012; Wang *et al*., 2012; Mackay, 2014; Jiang and Reif, 2015; Ober *et al*., 2015; Rönnegård and Shen, 2016).

The major concern in utilizing epistasis models has been the high computational load (Mackay, 2014) which has been reduced for the full model including all interactions by utilizing marker based epistasis relationship matrices (Jiang and Reif, 2015; Martini *et al*., 2016). In this work, epistasis relationship matrices were constructed by using the package miraculix (Schlather, 2020) which carries out matrix multiplications about 15 times faster than regular matrix multiplications on genotype data in R. In the analyzed datasets containing up to 30’212 SNPs, the computing time required to set up the sERRBLUP relationship matrix was about 810 minutes out of which around 330 minutes were required to estimate all pairwise SNP interaction effects and 480 minutes were required to set up the sERRBLUP relationship matrix for selected proportion of interactions by utilizing the R-package miraculix with 15 cores on a server cluster with Intel E5-2650 (2X12 core 2.2GHz) processors. Computing times for sERRBLUP scale approximately quadratic in the number of markers considered. The EpiGP R-package has been released for genomic prediction of phenotypes based on ERRBLUP and sERRBLUP (Vojgani *et al*., 2019a).

The predictive ability of EG-BLUP depends on marker coding and the symmetric coding seems to perform best. In contrast, the ERRBLUP (called “categorical epistasis (CE)” by Martini *et al*. (2017)) does not rely on marker coding and performs as good as EG-BLUP with symmetrically coded markers (Martini *et al*., 2017). Although the marker matrix generated with ERRBLUP and sERRBLUP is 4 times as large than the EG-BLUP interaction marker matrix, the computing time is comparable to EG-BLUP. Still, our proposed epistasis models eventually can generate a considerable prohibitive computational load if the number of SNPs grows to hundreds of thousands (Vojgani, *et al*., 2019). The computing time for sERRBLUP exhibits quadratic growth with increasing number of SNPs. Potential strategies to overcome these limitations are to achieve a feature reduction by SNP pruning, as was implemented in our maize dataset (Purcell *et al*., 2007; Chang *et al*., 2015). Another option might be the use of haplotype blocks (Pook *et al*., 2019).

In this study, we showed that the predictive ability obtained from univariate GBLUP and from a univariate full epistasis model with all pairwise SNP interactions included (ERRBLUP) was almost identical. In contrast, it was shown that the univariate sERRBLUP across environments increases predictive ability when the most relevant SNP interactions are taken into account. Based on the preliminary analysis in our datasets, univariate sERRBLUP across environments did not provide a considerable increase in predictive ability until 95 percent of pairwise SNP interactions were removed which means around 30 million interactions still remained in the model. It was also demonstrated in a wheat dataset (Crossa *et al*., 2010; Pérez and de los Campos, 2014) that the highest gain in predictive ability was obtained when over 80 percent were removed (Martini *et al*., 2016). Ober *et al*. (2015) concluded that the improvement of predictive ability when selecting only the top interaction - as we did in sERRBLUP - is likely a result of enriching for true causal variants among the list of variants used to construct the genetic covariance matrix. In our study, the maximum predictive ability with univariate sERRBLUP was obtained by incorporating the top 1 or 0.1 percent of pairwise SNP interactions, while a too strict selection of SNP interactions such as the top 0.01, 0.001 and 0.0001 percent reduced the predictive ability. A similar loss in predictive ability with a too strict selection of interactions to be included in the model was also observed by Ober et al. (2015). The difference in interaction selection may be explained by the absolute number of interaction effects in the model being more important than the percentage as well as potential differences in linkage leading to different redundancy patterns of interactions. Here we also saw the only major systematic difference between the two selection criteria: when SNP interactions were selected based on the magnitude of their estimated (absolute) effects, the loss in predictive ability when selecting too few interactions was much more severe than when SNP interactions were selected based on the variance associated with them (see Fig. 5). This phenomenon has been more prevalent in KE than in PE (Fig. 5 vs. Fig. 6), and is valid in both scenarios, using information either from a single environment or from the average of all other environments (Fig. 14 and Fig. 15). Conceptually, the SNP interaction effect variance appears to be the better choice, since extreme estimates of SNP interaction effects can result in cases where some interaction classes are just represented by very few lines. This is balanced by taking into account the frequency in the SNP interaction variance. Due to the conceptual advantage and the more robust performance we recommend the latter to be used as selection criterion in sERRBLUP applications.

The bivariate models exhibited a considerably higher predictive ability than univariate models, in consequence the maximum bivariate GBLUP performed slightly better than the maximum univariate sERRBLUP in most cases (Fig. S11), and the predictive ability increased further in the bivariate sERRBLUP when the top 5 or 1 percent of pairwise SNP interactions are selected based on the effects variances in most cases. Over all studied traits, the increase in prediction accuracy from GBLUP to sERRBLUP displays a similar pattern in both univariate and bivariate models. It should be noted, though, that the increase in prediction accuracy is limited by the prediction accuracy obtained with GBLUP in relation to the trait heritability, which is illustrated with the following example: for the trait final plant height (PH_final) the GBLUP prediction accuracy is 0.73, while the maximum possible value is the square root of the trait heritability, which is 0.97 in this case. Thus, the maximum possible absolute increase would be 0.24 which is 32.9 per cent of the prediction accuracy with GBLUP. So, the higher the GBLUP prediction accuracy is, the less room for improvement remains. However, the increase in predictive ability from bivariate GBLUP to bivariate sERRBLUP is only caused by the modelling of epistasis.

Cross validation in multi-trait genomic prediction models utilizing the secondary trait’s full dataset for prediction of the test set in the focal trait was shown to bias prediction accuracy (Runcie and Cheng, 2019), the main explanation being the existence of non-genetic covariance between the two traits observed in the same individuals, which is not properly accounted for in the prediction model. This scenario, however, differs from the cross-validation scenario studied in our case, since here the two ‘traits’ are actually the same biological trait observed in the same genotype, but in two completely separated environments. This setting should not give rise to non-genetic correlations between the two ‘traits’, e.g. caused by identical weather conditions affecting both traits simultaneously. Runcie and Cheng (2019) observed no systematic bias when the individuals do not share the same source of non-genetic variation, and thus, we assume that this source of bias is not relevant in our study.

In our scenario, the prediction accuracy can be reduced, though, if the second environment has a smaller number of phenotyped lines than the target environment indicating less overlap of genotypes with the target environment. In tendency this was confirmed in our study when GOL or TOM were modeled as the second environment for predicting the unobserved lines in other environments (e.g. BBG), since these two environments have around half the number of lines compared to other environments.

It has been shown that prediction accuracy is positively correlated to trait heritability. For instance, it was reported that prediction accuracies for resistance to yellow and stem rust in wheat was related to their heritability, respectively (Zhao *et al*., 2013; Momen *et al*., 2018). Similarly it was reported in maize that grain yield with low heritability has a prediction accuracy of 0.58, while grain moisture with high heritability has a prediction accuracy of 0.90 (Technow *et al*., 2014; Momen *et al*., 2018). Across the multitude of traits, environments, lines, and prediction methods, we could also find a substantial positive correlation between the heritabilities and the predictive abilities (see Fig. 12 – 13), which was more pronounced for sERRBLUP compared to GBLUP. The smallest correlation between heritability and the predictive ability was obtained with the univariate GBLUP model, while the maximum correlation of these two quantities was obtained with the bivariate sERRBLUP model.

Predictive ability in each environment can be increased by borrowing information especially from environments which have high phenotypic and genetic correlations to the target environment, which should be related to geographic and climatic similarities of the environments. In our study, the maximum predictive ability in each environment was obtained by sERRBLUP selecting interaction terms in environments with similar geographic features. This was observed for the majority of environments across series of phenotypic traits. The PCA based on environmental features (Fig. S12) also confirms that the environments which were closely located shared more common geographical and climatic features resulting in higher accuracy for prediction across the respective environments. To illustrate this, BBG and EIN which are closely located in Germany are close to each other in the PCA as well. This was also observed for ROG and OLI in Germany. GOL and TOM which are also closely located in Spain are close to each other in the PCA, too. For instance, our results indicate that in most cases the highest gain in phenotype prediction of e.g. BBG was obtained for sERRBLUP when borrowing information from EIN and vice versa. This was also observed in the majority of cases in bivariate models such that if the additional environment (i.e. EIN) which was added to the model shared more common geographical and climatic features with the target environment (i.e. BBG), the obtained predictive ability was higher compared to adding other environments as an additional environment (i.e. GOL) joint with target environment (i.e. BBG).

Rather than using a single environment to select SNP interactions for univariate sERRBLUP prediction, we also used an approach in which SNP interaction selection was based on all five environments jointly to evaluate the model in the sixth environment. This was also done for bivariate GBLUP, ERRBLUP and sERRBLUP such that the additional environment was the combination of all the other five environments instead of a single environment. In both univariate and bivariate models, it was shown that the obtained predictive ability across multiple environments jointly was mostly equivalent or higher than the maximum predictive ability obtained based on a single environment. Using an average across all other environments appears to be a robust alternative which in most cases will yield a result that is as good or even better than the best single environment, at the same time avoiding the risk of compromising prediction quality by choosing the ‘wrong’ training environment.

Overall, our results demonstrate that bivariate models can outperform univariate models and epistatic interactions can substantially increase the predictive ability. In the context of univariate models, it was shown that selecting a suitable subset of interactions based on other environments where phenotypic data of the full set of lines are available can substantially increase the predictive ability. Additionally, selecting a suitable subset of interactions based on all the other environments jointly performs as well as selecting a suitable subset of interactions based on the single environment with highest phenotypic correlation with the target environment.

The presented approach can be useful in cases, where different lines are grown in multiple environments, which can be either simultaneously, i.e. in the same season, or in subsequent seasons. We have shown, that ‘borrowing’ information, however just in the selection of the most relevant interactions, can substantially improve the phenotype prediction accuracy in another environment. This can be useful in sparse testing designs, e.g. where not all lines are grown in all environments. The suggested approach can be used to ‘impute’ missing phenotypes with a much increased accuracy compared to conventional approaches.

## Declaration

### Funding

This work was funded by German Federal Ministry of Education and Research (BMBF) within the scope of the funding initiative “Plant Breeding Research for the Bioeconomy” (MAZE – “Accessing the genomic and functional diversity of maize to improve quantitative traits”; Funding ID: 031B0195)

### Conflict of interest

On behalf of all authors, the corresponding author states that there is no conflict of interest.

### Ethics approval

The authors declare that this study complies with the current laws of the countries in which the experiments were performed.

### Consent to participate

Not applicable

### Consent for publication

Not applicable

### Availability of data and materials

All data and material are available through material transfer agreements upon request.

### Code availability

Not applicable

### Authors’ contributions

EV derived the results, analyzed the data, wrote the manuscript; TP proposed epistasis relationship matrices. JWRM proposed epistasis interaction selection; ACH, MM and CCS prepared the material; ACH proposed cross validation strategy in bivariate model; HS proposed the original research question, guided the structure of the research.TP JWRM ACM MM CCS HS read, revised and approved the manuscript.

## Acknowledgements

We are thankful to the technical staff at KWS SAAT SE, Misión Biológica de Galicia, Spanish National Research Council (CSIC), Technical University of Munich, and University of Hohenheim for providing the extensive phenotypic evaluation. We are grateful to the German Federal Ministry of Education and Research (BMBF) for the funding of our project within the scope of the funding initiative “Plant Breeding Research for the Bioeconomy” (MAZE – “Accessing the genomic and functional diversity of maize to improve quantitative traits”; Funding ID: 031B0195).

## Notes

### Competing Interest Statement

The authors have declared no competing interest.

