## Supplementary Material for "Accounting for epistasis improves genomic prediction of phenotypes with univariate and bivariate models across environments"

Theoretical and Applied Genetics

### Supplemental tables

**Table S1** The mean, minimum, maximum and standard deviation of phenotypic traits in each location for KE (blue numbers) and PE (red numbers).

| Trait | Location | Mean | Minimum | Maximum | Standard deviation |
|---|---|---|---|---|---|
| | BBG | 4.10\4.68 | 0.78\1.00 | 7.28\7.55 | 1.28\1.18 |
| | EIN | 4.11\4.70 | 0.86\1.03 | 9.00\9.03 | 1.31\1.18 |
| EV_V3 | OLI | 5.31\6.17 | 1.22\3.30 | 8.05\8.74 | 1.15\0.86 |
| | ROG | 5.35\5.84 | 1.71\2.90 | 7.90\7.92 | 0.95\0.75 |
| | GOL | 6.31\6.67 | 4.07\5.49 | 8.49\7.98 | 0.69\0.51 |
| | TOM | 5.51\6.15 | 1.93\3.84 | 7.34\8.45 | 0.99\0.67 |
| | BBG | 3.85\4.65 | 0.67\0.93 | 8.29\8.49 | 1.48\1.48 |
| | EIN | 4.24\4.82 | 0.94\1.52 | 7.07\7.46 | 1.11\0.98 |
| EV _V4 | OLI | 5.27\6.07 | 0.80\2.99 | 7.52\8.36 | 1.08\0.75 |
| | ROG | 5.44\5.85 | 2.65\2.88 | 7.86\7.94 | 0.92\0.78 |
| | GOL | 5.71\5.98 | 3.37\3.91 | 7.89\7.89 | 0.81\0.83 |
| | TOM | 5.26\5.75 | 2.59\3.92 | 6.89\7.35 | 0.83\0.61 |
| | BBG | 3.92\4.64 | 0.74\0.84 | 8.75\8.22 | 1.39\1.41 |
| | EIN | 5.03\5.54 | 0.97\1.51 | 8.05\8.39 | 1.24\1.06 |
| EV _V6 | OLI | 5.30\6.07 | 0.54\3.56 | 7.17\8.09 | 0.96\0.74 |
| | ROG | 5.55\5.91 | 1.02\2.52 | 8.07\7.76 | 0.95\0.77 |
| | GOL | 6.24\6.24 | 3.90\3.81 | 8.45\7.94 | 0.85\0.85 |
| | TOM | 5.58\5.86 | 2.96\3.90 | 7.66\7.91 | 0.92\0.68 |
| | BBG | 35.86\41.48 | 9.27\16.38 | 60.40\62.85 | 8.43\7.93 |
| | EIN | 34.49\38.73 | 6.90\20.43 | 53.14\57.94 | 7.24\6.17 |
| PH_V4 | OLI | 18.43\22.55 | 7.35\11.89 | 31.11\35.75 | 3.93\3.87 |
| | ROG | 25.50\28.10 | 9.23\13.63 | 42.29\41.54 | 4.60\4.53 |
| | GOL | 62.88\68.98 | 34.30\38.39 | 88.24\95.30 | 9.79\10.96 |
| | TOM | 41.60\47.45 | 11.98\25.37 | 63.89\72.12 | 8.71\8.27 |
| | BBG | 61.75\69.08 | 19.41\30.36 | 93.84\100.39 | 11.80\11.12 |
| | EIN | 62.40\69.36 | 21.41\36.53 | 95.54\98.80 | 11.89\9.62 |
| PH_V6 | OLI | 36.74\45.35 | 8.34\14.78 | 58.40\72.48 | 8.69\8.53 |
| | ROG | 61.46\68.91 | 32.17\30.35 | 89.74\94.77 | 9.34\9.52 |
| | GOL | 94.21\98.30 | 37.28\54.75 | 127.54\130.51 | 15.05\15.29 |
| | TOM | 83.86\92.35 | 48.46\57.81 | 119.07\124.98 | 14.41\12.79 |
| | BBG | 142.65\120.60 | 95.85\59.78 | 210.08\179.03 | 20.53\19.11 |
| | EIN | 159.18\141.35 | 100.84\69.01 | 228.96\211.14 | 21.57\21.10 |
| PH_final | OLI | 118.17\112.46 | 58.74\58.55 | 175.81\173.15 | 21.95\20.48 |
| | ROG | 137.04\122.25 | 74.25\63.56 | 211.14\201.92 | 22.32\20.56 |
| | GOL | 115.68\102.69 | 49.27\30.21 | 167.58\149.14 | 21.73\23.59 |
| | TOM | 157.99\144.61 | 81.92\79.28 | 245.00\195.36 | 24.82\18.95 |

| | | | | | |
|---|---|---|---|---|---|
| | **BBG** | 82.10\82.08 | 69.45\69.78 | 95.74\92.04 | 4.31\4.18 |
| | **EIN** | 82.55\81.78 | 70.36\68.86 | 102.02\101.50 | 5.23\5.17 |
| **FF** | **OLI** | 83.32\82.41 | 72.60\69.46 | 92.13\91.54 | 3.76\3.68 |
| | **ROG** | 73.06\71.91 | 62.45\59.10 | 91.22\88.03 | 4.82\4.47 |
| | **TOM** | 76.88\74.16 | 63.93\62.13 | 93.28\92.17 | 5.58\4.64 |
| | **BBG** | 5.02\3.03 | 0.59\0.03 | 9.58\9.22 | 2.78\2.39 |
| | **EIN** | 3.48\2.23 | 0.63\0.76 | 9.21\8.08 | 2.29\1.54 |
| **RL** | **OLI** | 2.59\1.80 | 0.59\0.52 | 9.15\7.65 | 1.64\1.19 |
| | **ROG** | 2.39\1.50 | 0.96\0.95 | 9.01\8.50 | 2.21\1.13 |

**Table S2** The combinations which did not converged in bivariate sERRBLUP with the number of not converged folds in 5-fold cross validation with 5 replicates for trait PH-V4 in KE (blue numbers) and PE (red numbers). The blue and red zeros in the parentheses represent the non-convergence of pre estimated variance components based on the full set in KE and PE, respectively.

| Predicted Environment | Additional Environment | Top 5 | Top 1 | Top 0.1 | Top 0.01 | Top 0.001 |
|---|---|---|---|---|---|---|
| BBG | EIN | - | - | - | - | (0) |
| BBG | GOL | (0)25 | 23(0) | 22 | - | |
| EIN | BBG | - | - | - | - | (0) |
| EIN | GOL | (0) | (0) | 18(0) | - | - |
| OLI | GOL | 21(0) | 21(0) | (0) | - | - |
| ROG | BBG | - | - | - | - | (0) |
| ROG | GOL | (0) | (0) | (0) | - | - |
| TOM | BBG | (0) | (0) | (0) | (0)25 | (0) / (0) |
| TOM | EIN | - | (0) | (0) | 1 | (0) |
| TOM | GOL | (0) | (0) | (0) | 9 | (0) |

**Table S3** The combinations which did not converged in bivariate sERRBLUP with the number of not converged folds in 5-fold cross validation with 5 replicates for trait EV-V3 in KE (blue numbers) and PE (red numbers). The blue and red zeros in the parentheses represent the non-convergence of pre estimated variance components based on the full set in KE and PE, respectively.

| Predicted Environment | Additional Environment | Top 5 | Top 1 | Top 0.1 | Top 0.01 | Top 0.001 |
|---|---|---|---|---|---|---|
| BBG | EIN | - | - | - | - | 1(0) |
| BBG | GOL | 1 | - | - | - | - |
| BBG | TOM | - | - | - | - | 3 |
| EIN | ROG | - | - | 2 | 1 | 2 |
| EIN | BBG | - | - | - | - | 3(0) |
| EIN | TOM | - | - | 1 | 1 | 1 |
| OLI | ROG | - | - | - | 2 | 1 |
| OLI | EIN | - | - | - | 2 | 25(0) / 1 |
| OLI | BBG | - | - | 1 | 1 / 3 | (0) / 6 |
| OLI | GOL | 1 | 1 | 3 | 1 | 2 |
| OLI | TOM | 2 | 1 | 7 | 7(0) | 10(0) |
| ROG | EIN | - | - | 1 | 12(0) | 25(0) / 6 |
| ROG | BBG | - | - | - | 1 / 2 | 25(0) |
| ROG | GOL | - | - | 1 | 2 | 1 |
| ROG | TOM | - | - | - | 4 | - |
| GOL | ROG | - | - | - | 2 | - |
| GOL | EIN | - | - | - | - | 25(0) |
| GOL | BBG | - | - | 1 | 1 | 5(0) / 1 |
| GOL | OLI | 1 | 2 | 1 | 2 | 3 |
| GOL | TOM | - | - | - | 3 / 5 | 5 / 1 |
| TOM | EIN | 1 | 2 | 9 | 14(0) | 25(0) |
| TOM | BBG | - | - | - | 1 | 5(0) |
| TOM | OLI | 1 | - | - | - | 2 |

| | | | | | | |
|---|---|---|---|---|---|---|
| TOM | GOL | 25(0) | 1 | - | 5 | 1 / 1 |

**Table S4** The combinations which did not converged in bivariate sERRBLUP with the number of not converged folds in 5-fold cross validation with 5 replicates for trait EV-V4 in KE (blue numbers) and PE (red numbers). The blue and red zeros in the parentheses represent the non-convergence of pre estimated variance components based on the full set in KE and PE, respectively.

| Predicted Environment | Additional Environment | Top 5 | Top 1 | Top 0.1 | Top 0.01 | Top 0.001 |
|---|---|---|---|---|---|---|
| BBG | EIN | - | - | - | - | 22(0) |
| BBG | GOL | - | - | 1 | 8 | 11 |
| EIN | OLI | - | 1 | - | - | - |
| OLI | EIN | 3 | 3 | 1 / 21(0) | 1 / 24(0) | 23(0) / 23(0) |
| OLI | BBG | - | - | 1 | - | - |
| OLI | GOL | - | - | - | 1 | - |
| OLI | TOM | - | - | - | 1 | 1 |
| ROG | EIN | - | - | - | - | 25(0) |
| GOL | EIN | - | - | - | - | 25(0) |
| GOL | TOM | - | 1 | - | - | - |
| TOM | ROG | - | 1 | - | 2 | - |
| TOM | EIN | - | 1 | 3 | - | - |
| TOM | BBG | - | 1 | 18(0) | 15(0) | 5 |
| TOM | GOL | 5 | 5 | 1 / 13(0) | 1 | 5 |

**Table S5** The combinations which did not converged in bivariate sERRBLUP with the number of not converged folds in 5-fold cross validation with 5 replicates for trait EV-V6 in KE (blue numbers) and PE (red numbers). The blue and red zeros in the parentheses represent the non-convergence of pre estimated variance components based on the full set in KE and PE, respectively.

| Predicted Environment | Additional Environment | Top 5 | Top 1 | Top 0.1 | Top 0.01 | Top 0.001 |
|---|---|---|---|---|---|---|
| BBG | ROG | - | - | - | - | 5(0) |
| BBG | EIN | - | - | 1 | 2 | 25(0) |
| BBG | GOL | 25(0) / 18(1) | 25(0) / 13(0) | 19(0) / 19(0) | 25(0) | - |
| EIN | ROG | - | - | 1 | 5(0) / 1 | 12(0) / 1 |
| EIN | GOL | 24(0) / 25(0) | 24(0) / 3(0) | 13(0) / 25(0) | 17 | - |
| OLI | EIN | - | - | 3 | 3 | 24(0) / 4 |
| OLI | ROG | - | - | - | - | 25(0) |
| OLI | GOL | 18(0) / 25(0) | 7 / 1 | 19(0) / 24(0) | 22(0) | - |
| ROG | EIN | - | - | 2 | 5 | 15(0) / 3 |
| ROG | BBG | 1 | 1 | - | 1 / 4(0) | 1 / 12(0) |
| ROG | OLI | - | - | 1 | - | 1 |
| ROG | GOL | 18 / 25(0) | 25(0) / 1 | 25(0) / 16 | 25 | - |
| GOL | ROG | - | - | - | - | 1(0) |
| GOL | EIN | - | - | - | - | 5(5) |
| TOM | ROG | - | 1 | 19(0) | 10(0) / 1 | 25(0) /7 |
| TOM | EIN | - | - | 7 | 7 | 25(0) |
| TOM | BBG | - | 6 | 6 / 1 | 4 | 4 |
| TOM | OLI | - | - | - | - | 1 |
| TOM | GOL | 25(0) / 25 | 25(0) / 4 | 25(0) / 25 | 21(0) / 25(0) | 20(0) / 10(0) |

**Table S6** The combinations which did not converged in bivariate sERRBLUP with the number of not converged folds in 5-fold cross validation with 5 replicates for trait PH-V6 in KE (blue numbers) and PE (red numbers). The blue and red zeros in the parentheses represent the non-convergence of pre estimated variance components based on the full set in KE and PE, respectively.

| Predicted Environment | Additional Environment | Top 5 | Top 1 | Top 0.1 | Top 0.01 | Top 0.001 |
|---|---|---|---|---|---|---|
| BBG | ROG | - | - | - | 1 | 5(0) |
| BBG | EIN | 2 | - | - | - | - |
| BBG | OLI | - | - | - | - | 16 |
| BBG | GOL | - | - | 1 | 1 | 2(0) |
| EIN | ROG | - | - | - | - | 25(0) |
| EIN | BBG | - | - | 5 | - | 17(0) |
| EIN | OLI | - | - | - | - | 1 |
| EIN | GOL | - | - | - | - | (0) |
| OLI | ROG | - | - | 1 | 8 | 21(0) |
| OLI | EIN | - | - | - | - | 2 |
| OLI | BBG | - | - | - | - | 11(0) |
| OLI | GOL | - | - | - | 1 | 25(0) |
| ROG | EIN | - | - | - | - | 2 |
| ROG | BBG | - | - | - | 1 | 4(0) |
| ROG | OLI | - | - | - | - | 25(0) |
| ROG | GOL | - | - | - | - | 23(0) |
| GOL | ROG | - | - | - | - | 1(0) |
| GOL | BBG | - | - | - | - | 25(0) / 1 |
| GOL | OLI | - | - | - | - | 25(0) |
| TOM | ROG | - | - | 2 | 3 | 15(0) |
| TOM | EIN | - | - | - | - | - |
| TOM | BBG | - | - | 17 | 2 | 25(0) |
| TOM | OLI | - | - | - | 1 | 24(0) |

| | | | | | | |
|---|---|---|---|---|---|---|
| TOM | GOL | 12(1) | 8(0) / 3 | 7(0) / 2 | 2 | 21(0) / 1 |

**Table S7** The combinations which did not converged in bivariate sERRBLUP with the number of not converged folds in 5-fold cross validation with 5 replicates for trait PH-final in KE (blue numbers) and PE (red numbers). The blue and red zeros in the parentheses represent the non-convergence of pre estimated variance components based on the full set in KE and PE, respectively.

| Predicted Environment | Additional Environment | Top 5 | Top 1 | Top 0.1 | Top 0.01 | Top 0.001 |
|---|---|---|---|---|---|---|
| BBG | ROG | 12(0) / 25(0) | 25(0) / 25(0) | 25(0) / 25(0) | 25(0) | - |
| BBG | EIN | - | - | - | - | 1 |
| BBG | TOM | 16(0) | 24(0) | 25(0) | 2 | - |
| EIN | ROG | 25(0) / 18 | 25(0) / 25(0) | 25(0) / 25(0) | 1 / 25(0) | 2 |
| EIN | BBG | 23(0) | 24(0) | 25(0) | 1 | 1 |
| EIN | OLI | - | - | - | - | 2 |
| EIN | TOM | 9 | 24(0) | 25(0) | - | - |
| OLI | ROG | 25(0) / 24(0) | 25(0) / 25(0) | 25(0) / 25(0) | 25(0) | 1 |
| OLI | EIN | - | 1 | - | - | - |
| OLI | BBG | 24(0) | 25(0) | 25(0) | - | 1 |
| OLI | GOL | - | - | - | - | - |
| OLI | TOM | 18(0) | 22(0) | 25(0) | 1 / 4 | 2 |
| ROG | EIN | - | - | - | - | 1 |
| ROG | BBG | 20(0) | 25(0) | 25(0) | - | - |
| ROG | TOM | 19(0) | 23(0) | 24(0) | 3 | - |
| GOL | ROG | 23(0) / 25(0) | 24(0) / 24(0) | 23(0) / 24(0) | 25(0) | - |
| GOL | BBG | 23(0) | 24(0) | 24(0) | - | - |
| GOL | TOM | 25(0) | 25(0) | 24(0) | 3 | - |
| TOM | ROG | 25(0) / 25(0) | 25(0) / 25(0) | 25(0) / 25(0) | 16(0) / 25(0) | 13(0) / 1 |
| TOM | EIN | - | - | 4 | 6 | 15(0) / 1 |
| TOM | BBG | 23(0) | 25(0) | 25(0) | 3 | 12(0) |
| TOM | OLI | - | - | 2 | 2 | 1 |
| TOM | GOL | - | - | 1 | - | 1 |

**Table S8** The combinations which did not converged in bivariate sERRBLUP with the number of not converged folds in 5-fold cross validation with 5 replicates for trait FF in KE (blue numbers) and PE (red numbers). The blue and red zeros in the parentheses represent the non-convergence of pre estimated variance components based on the full set in KE and PE, respectively.

| Predicted Environment | Additional Environment | Top 5 | Top 1 | Top 0.1 | Top 0.01 | Top 0.001 |
|---|---|---|---|---|---|---|
| BBG | EIN | 24(0) | 24(0) | 25(0) | 2 / 17 (0) | 5 |
| BBG | ROG | 25(0) | 25(0) | 25(0) / 25(0) | 5 / 25(0) | 8 |
| EIN | BBG | 1 / 16(0) | 2 / 1 | 2 / 25(0) | 3(0) / 25(0) | 3 / 1 |
| EIN | ROG | 25(0) | 25(0) | 25(0) / 25(0) | 1 / 25(0) | 1 / 4 |
| OLI | ROG | 25(0) | 2 / 25(0) | 21(0) / 25(0) | 10 / 25(0) | 2 / 6 |
| OLI | BBG | - | 4 | 25(0) | 24(0) | 1 / 17(0) |
| OLI | EIN | 9(0) | 20 | 25(0) | 12(0) | 14(0) |
| ROG | EIN | 25(0) | 25(0) | 25(0) | 5 | 12(0) |
| ROG | BBG | 15 | | 25(0) | 25(0) | 1 |
| TOM | EIN | 18(0) | 20(0) | 19(0) | 15(0) | 7 / 22(0) |
| TOM | BBG | 7 | - | 24(0) | 23(0) | 1 |
| TOM | OLI | - | - | 1 | 1 | 1 |
| TOM | ROG | 22(0) | 1 / 25 | 25(0) / 24(0) | 20(0) / 24(0) | 24(0) / 15(0) |

**Table S9** The combinations which did not converged in bivariate sERRBLUP with the number of not converged folds in 5-fold cross validation with 5 replicates for trait RL in KE (blue numbers) and PE (red numbers). The blue and red zeros in the parentheses represent the non-convergence of pre estimated variance components based on the full set in KE and PE, respectively.

| Predicted Environment | Additional Environment | Top 5 | Top 1 | Top 0.1 | Top 0.01 | Top 0.001 |
|---|---|---|---|---|---|---|
| BBG | EIN | - | - | - | - | 1 |
| EIN | ROG | - | - | 1 | 1 | 1 |
| EIN | BBG | - | 7 | 10(0) | 20(0) | 1 / 21(0) |
| EIN | OLI | 2 | 2 | 2 | 1 | 8 |
| ROG | EIN | - | - | 1 | 3 | 8 |
| ROG | BBG | - | - | - | 1 | 1 |
| ROG | OLI | - | - | 1 | - | - |

**Table S10a** The predictive ability of univariate GBLUP within each environment (blue numbers) and the maximum predictive ability of univariate sERRBLUP across each environment (red numbers) in Kemater

| Location | BBG | EIN | OLI | ROG | GOL | TOM |
|---|---|---|---|---|---|---|
| EV_V3 | 0.448\0.638 | 0.369\0.587 | 0.400\0.610 | 0.372\0.548 | 0.419\0.683 | 0.388\0.659 |
| EV_V4 | 0.460\0.658 | 0.371\0.618 | 0.375\0.569 | 0.459\0.610 | 0.352\0.629 | 0.318\0.590 |
| EV_V6 | 0.418\0.609 | 0.327\0.574 | 0.416\0.590 | 0.314\0.533 | 0.466\0.623 | 0.247\0.454 |
| PH_V4 | 0.470\0.707 | 0.455\0.679 | 0.439\0.623 | 0.469\0.665 | 0.519\0.694 | 0.346\0.617 |
| PH_V6 | 0.472\0.714 | 0.473\0.703 | 0.321\0.596 | 0.450\0.659 | 0.480\0.728 | 0.413\0.662 |
| PH_final | 0.677\0.812 | 0.628\0.783 | 0.541\0.679 | 0.675\0.812 | 0.509\0.628 | 0.502\0.682 |
| FF | 0.537\0.735 | 0.414\0.691 | 0.400\0.527 | 0.580\0.720 | NA | 0.456\0.589 |
| RL | 0.497\0.603 | 0.459\0.571 | 0.432\0.570 | 0.533\0.653 | NA | NA |

**Table S10b** The predictive ability of univariate GBLUP within each environment (blue numbers) and the maximum predictive ability of univariate sERRBLUP across each environment (red numbers) in Petkuser

| Location | BBG | EIN | OLI | ROG | GOL | TOM |
|---|---|---|---|---|---|---|
| EV_V3 | 0.324\0.590 | 0.255\0.542 | 0.249\0.525 | 0.363\0.547 | 0.450\0.591 | 0.334\0.545 |
| EV_V4 | 0.381\0.632 | 0.294\0.564 | 0.238\0.502 | 0.325\0.467 | 0.567\0.646 | 0.293\0.422 |
| EV_V6 | 0.432\0.626 | 0.307\0.559 | 0.216\0.446 | 0.332\0.516 | 0.655\0.714 | 0.336\0.445 |
| PH_V4 | 0.424\0.648 | 0.412\0.652 | 0.324\0.558 | 0.449\0.584 | 0.604\0.664 | 0.370\0.528 |
| PH_V6 | 0.458\0.669 | 0.465\0.692 | 0.320\0.548 | 0.471\0.653 | 0.670\0.718 | 0.465\0.600 |
| PH_final | 0.628\0.761 | 0.634\0.779 | 0.558\0.707 | 0.682\0.828 | 0.704\0746 | 0.583\0.671 |
| FF | 0.582\0.795 | 0.539\0.764 | 0.443\0.591 | 0.521\0.773 | NA | 0.414\0.550 |
| RL | 0.432\0.537 | 0.265\0.377 | 0.355\0.509 | 0.308\0.387 | NA | NA |

### Supplemental figures

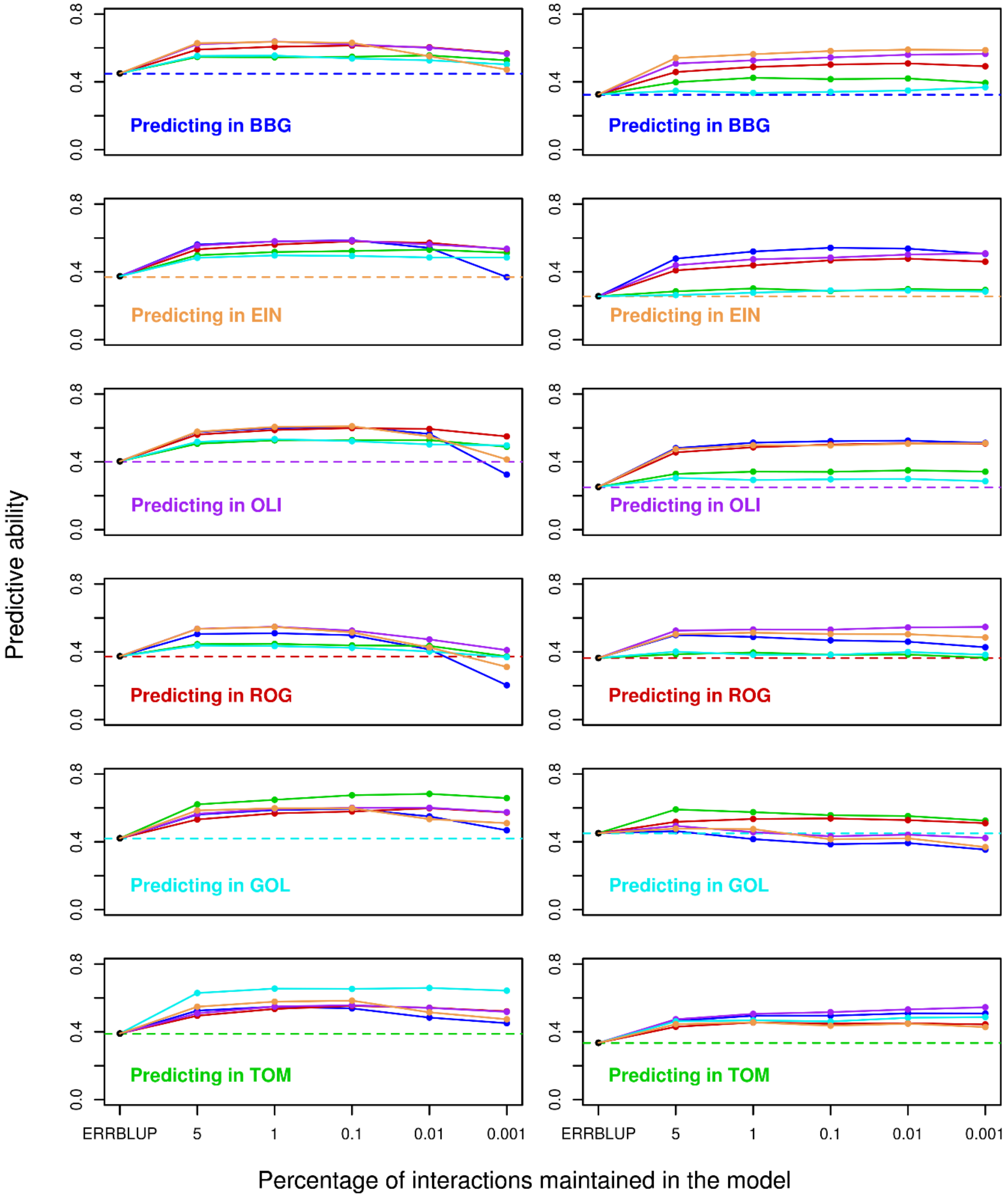


**Fig. S1a** Predictive ability for univariate GBLUP within environment (dashed horizontal line), univariate ERRBLUP within environment (black filled circle) and univariate sERRBLUP across environments when the SNP interaction selections are based on estimated effects variances (solid colored lines) for trait EV_V3 in KE (left side plots) and PE (right side plots). In each panel, the solid lines' color indicates the environment in which the relationship matrices determined by variable selection.

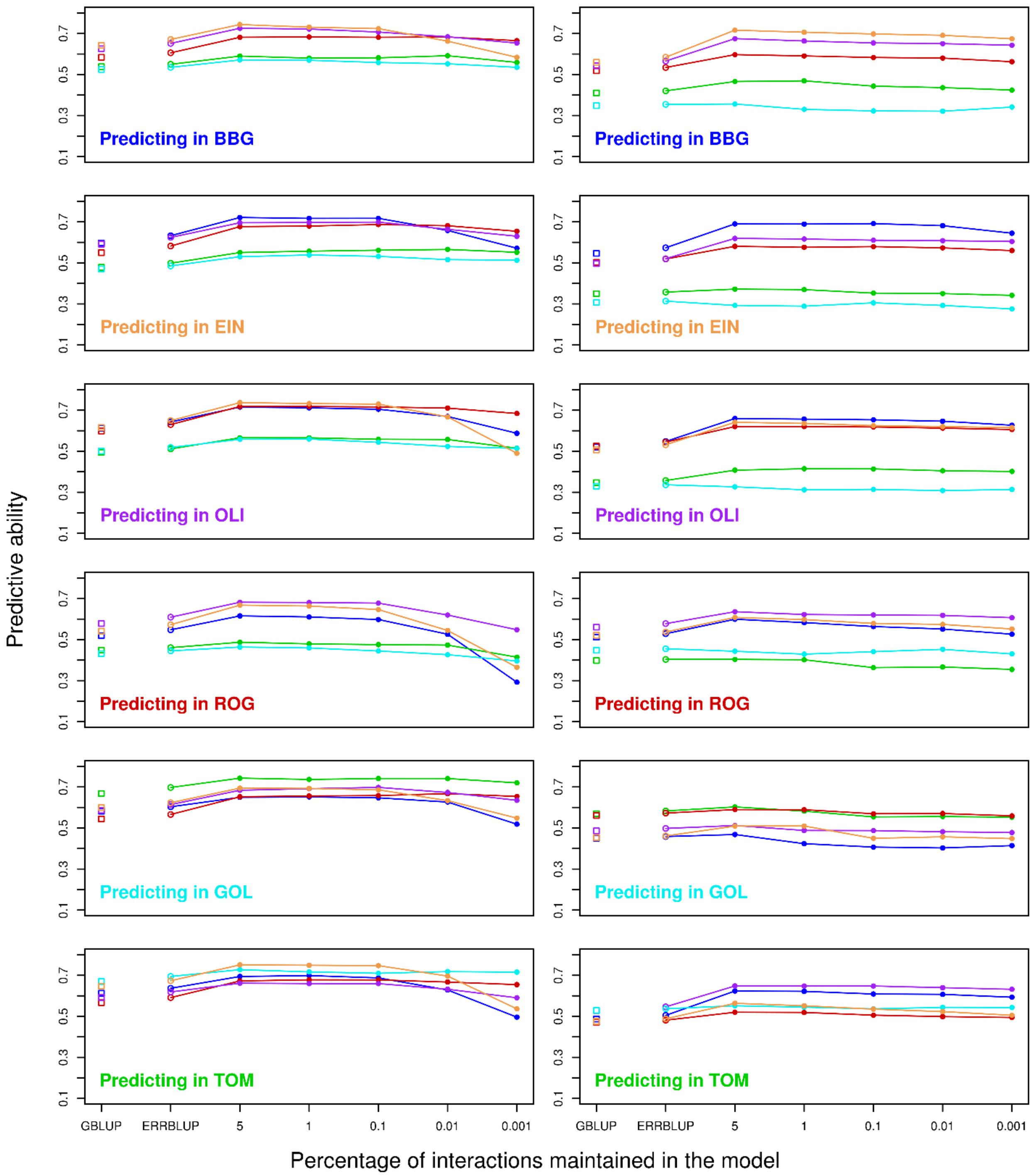


**Fig. S1b** Predictive ability for bivariate GBLUP (open squares), bivariate ERRBLUP (open circles) and bivariate sERRBLUP (filled circles and solid lines) when SNP interaction selections are based on estimated effects variances in Kemater (left side) and Petkuser (right side) for trait EV-V3. In each panel, the solid lines' color indicates the additional environment used to predict the target environment.

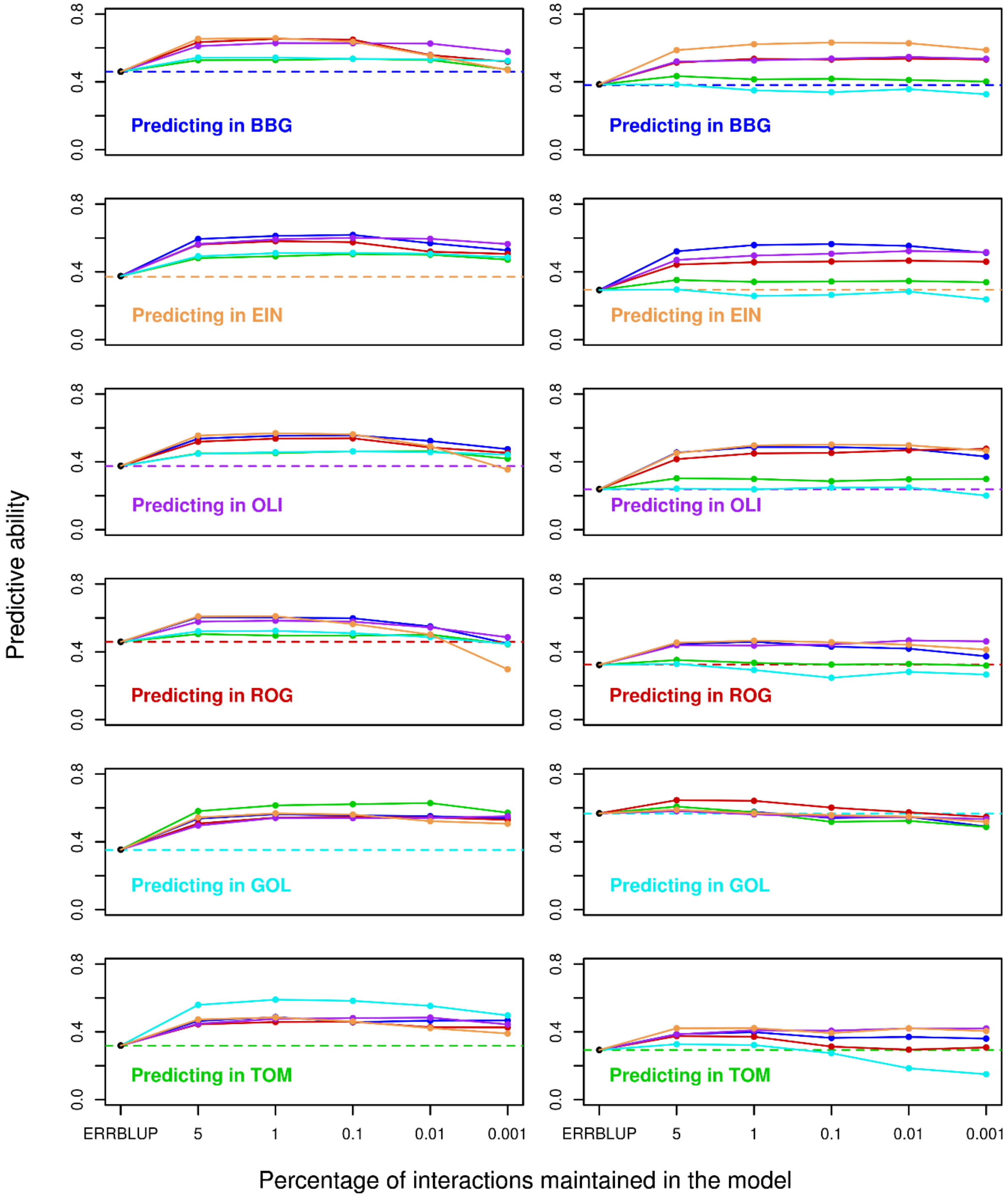


**Fig. S2a** Predictive ability for univariate GBLUP within environment (dashed horizontal line), univariate ERRBLUP within environment (black filled circle) and univariate sERRBLUP across environments when the SNP interaction selections are based on estimated effects variances (solid colored lines) for trait EV_V4 in KE (left side plots) and PE (right side plots). In each panel, the solid lines' color indicates the environment in which the relationship matrices determined by variable selection.

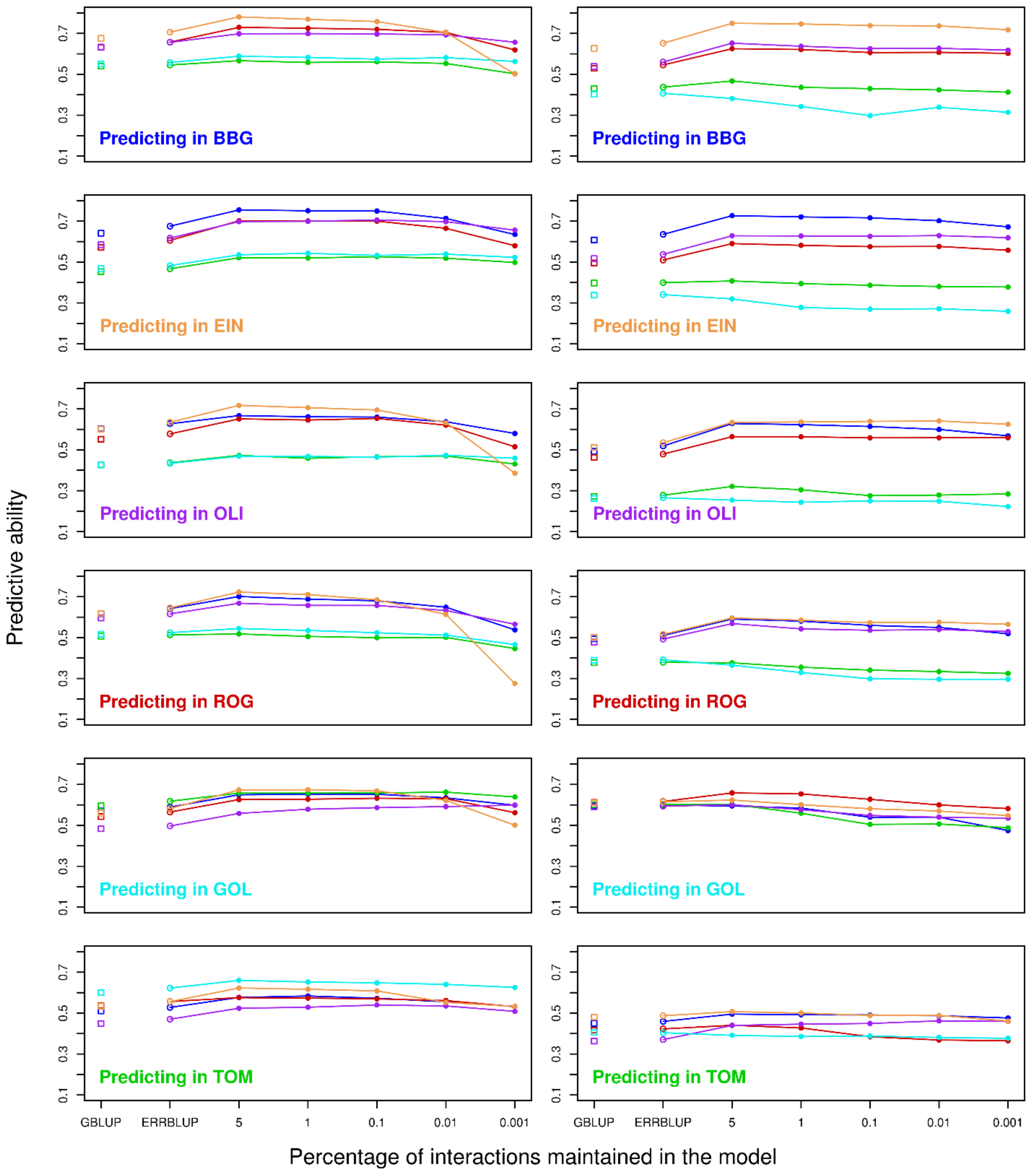


**Fig. S2b** Predictive ability for bivariate GBLUP (open squares), bivariate ERRBLUP (open circles) and bivariate sERRBLUP (filled circles and solid lines) when SNP interaction selections are based on estimated effects variances in Kemater (left side) and Petkuser (right side) for trait EV-V4. In each panel, the solid lines' color indicates the additional environment used to predict the target environment.

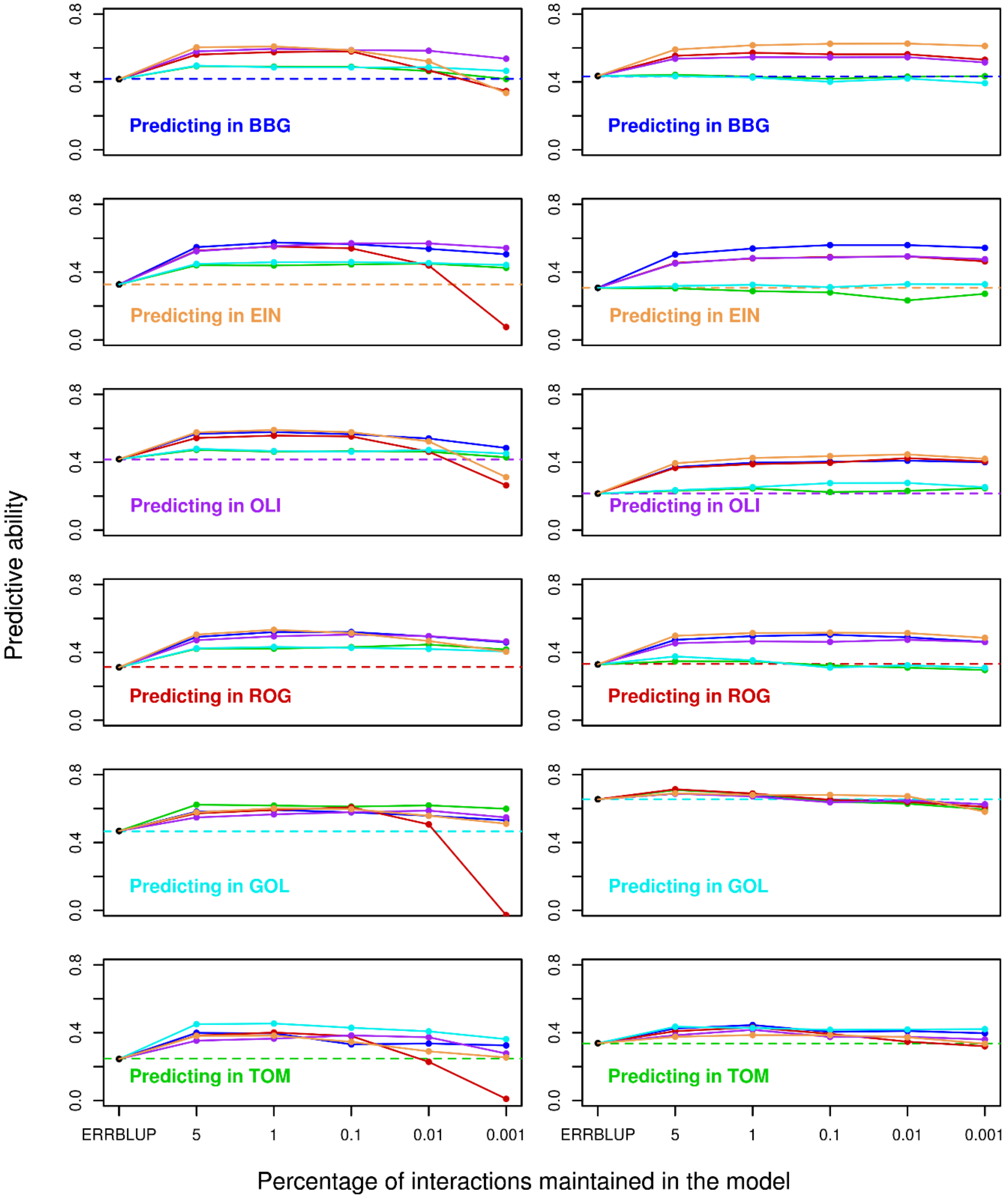


**Fig. S3a** Predictive ability for univariate GBLUP within environment (dashed horizontal line), univariate ERRBLUP within environment (black filled circle) and univariate sERRBLUP across environments when the SNP interaction selections are based on estimated effects variances (solid colored lines) for trait EV_V6 in KE (left side plots) and PE (right side plots). In each panel, the solid lines' color indicates the environment in which the relationship matrices determined by variable selection.

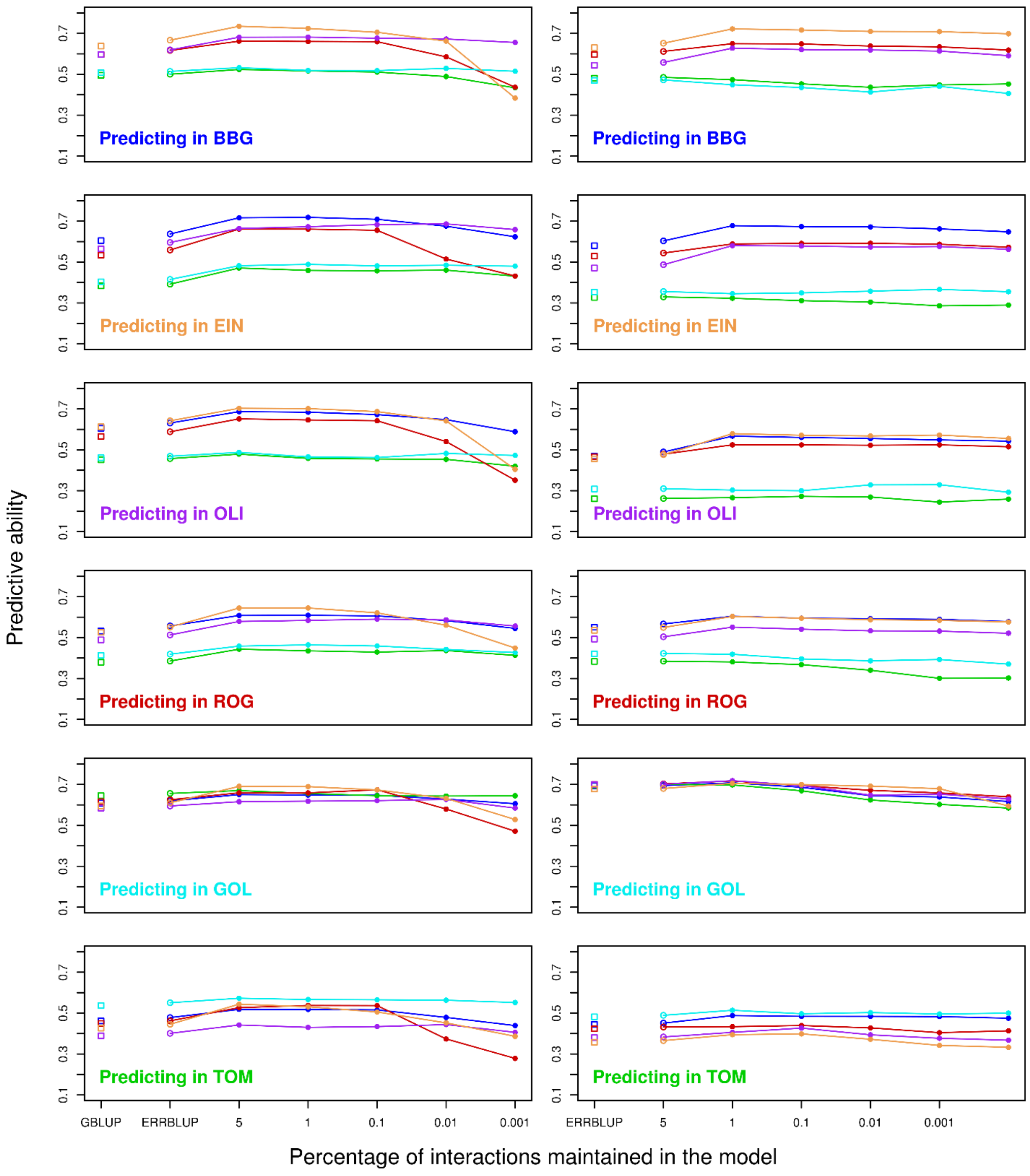


**Fig. S3b** Predictive ability for bivariate GBLUP (open squares), bivariate ERRBLUP (open circles) and bivariate sERRBLUP (filled circles and solid lines) when SNP interaction selections are based on estimated effects variances in Kemater (left side) and Petkuser (right side) for trait EV-V6. In each panel, the solid lines' color indicates the additional environment used to predict the target environment.

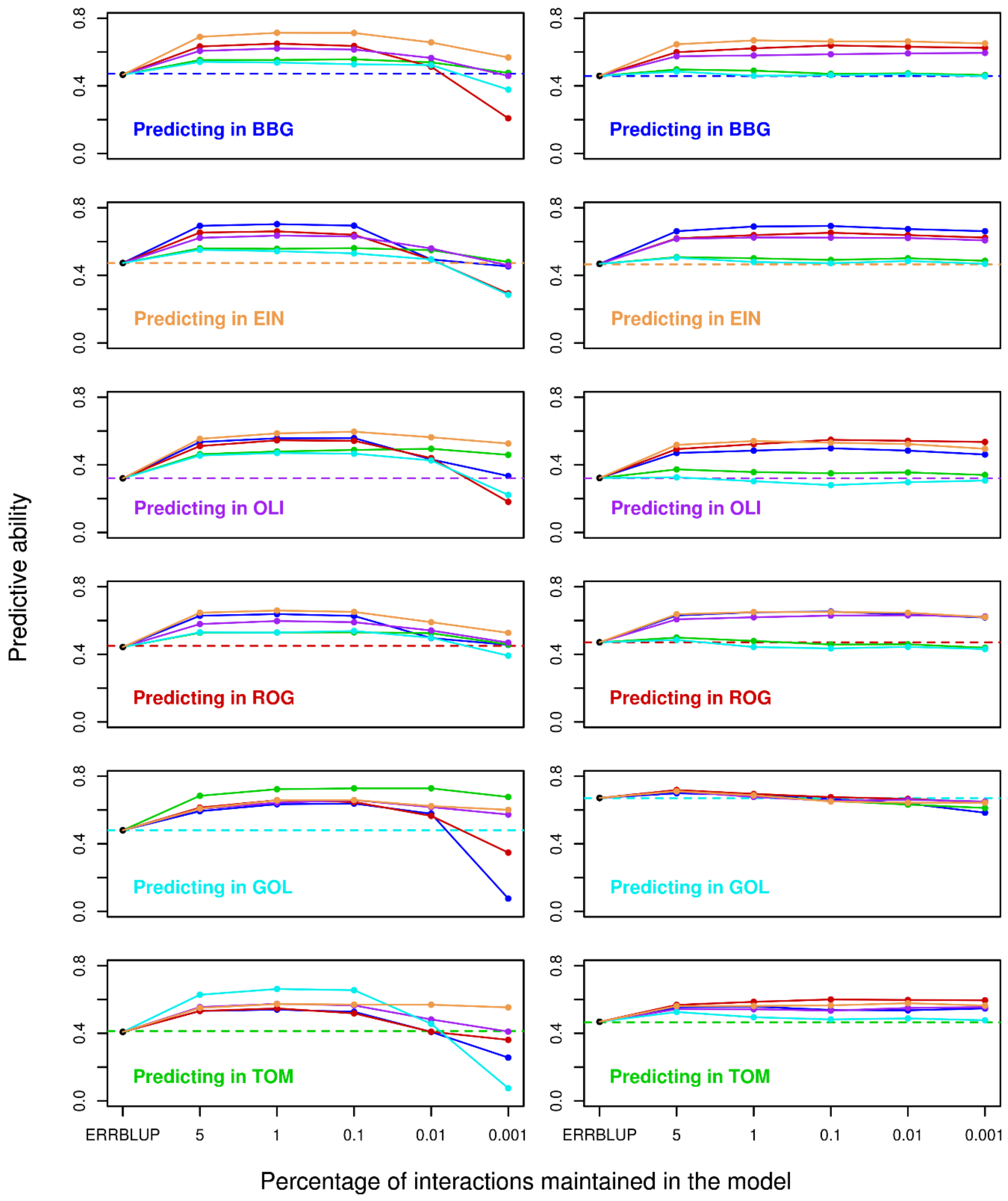


**Fig. S4a** Predictive ability for univariate GBLUP within environment (dashed horizontal line), univariate ERRBLUP within environment (black filled circle) and univariate sERRBLUP across environments when the SNP interaction selections are based on estimated effects variances (solid colored lines) for trait PH-V6 in KE (left side plots) and PE (right side plots). In each panel, the solid lines' color indicates the environment in which the relationship matrices determined by variable selection.

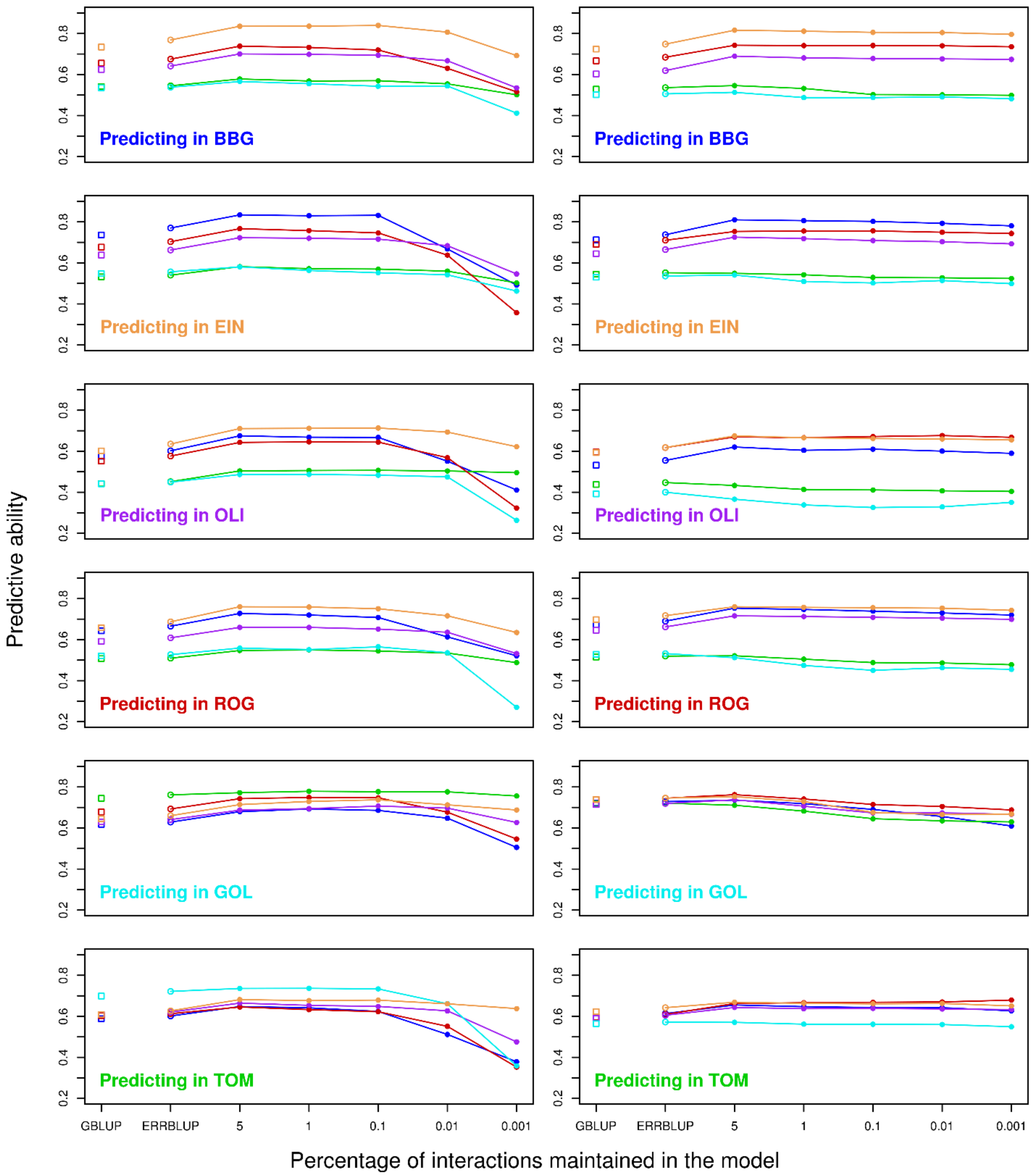


**Fig. S4b** Predictive ability for bivariate GBLUP (open squares), bivariate ERRBLUP (open circles) and bivariate sERRBLUP (filled circles and solid lines) when SNP interaction selections are based on estimated effects variances in Kemater (left side) and Petkuser (right side) for trait PH-V6. In each panel, the solid lines' color indicates the additional environment used to predict the target environment.

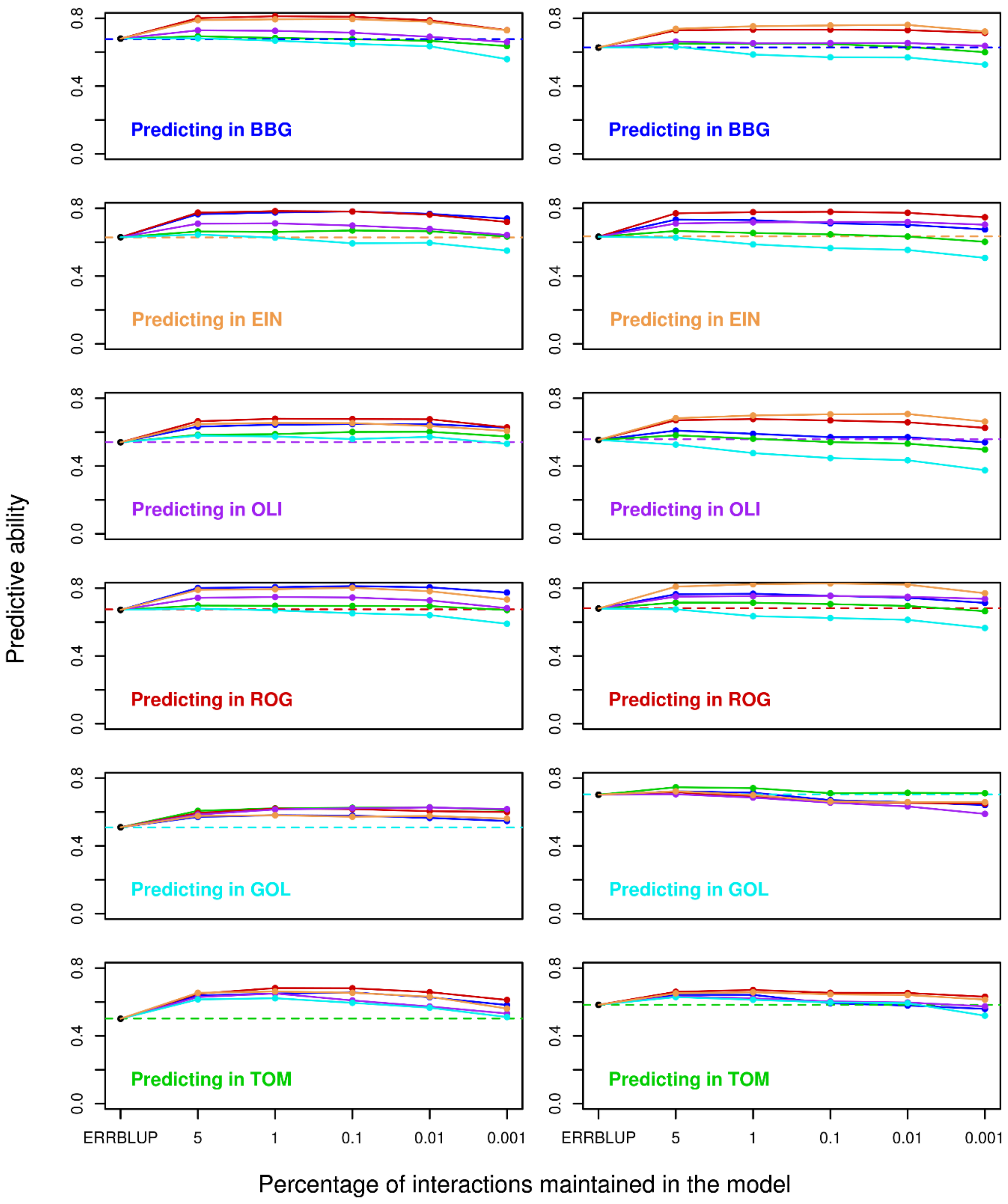


**Fig. S5a** Predictive ability for univariate GBLUP within environment (dashed horizontal line), univariate ERRBLUP within environment (black filled circle) and univariate sERRBLUP across environments when the SNP interaction selections are based on estimated effects variances (solid colored lines) for trait PH-final in KE (left side plots) and PE (right side plots). In each panel, the solid lines' color indicates the environment in which the relationship matrices determined by variable selection.

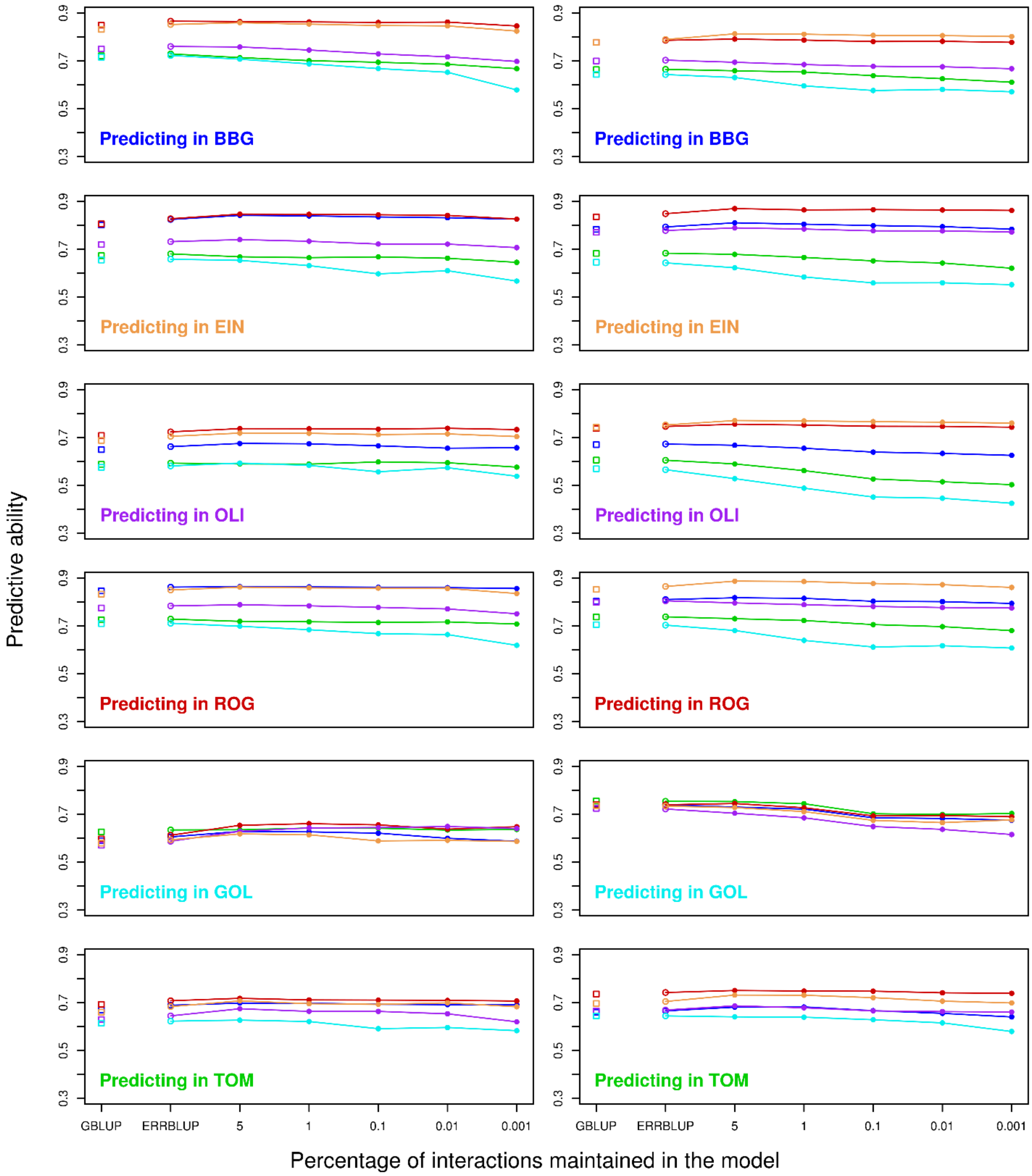


**Fig. S5b** Predictive ability for bivariate GBLUP (open squares), bivariate ERRBLUP (open circles) and bivariate sERRBLUP (filled circles and solid lines) when SNP interaction selections are based on estimated effects variances in Kemater (left side) and Petkuser (right side) for trait PH-final. In each panel, the solid lines' color indicates the additional environment used to predict the target environment.

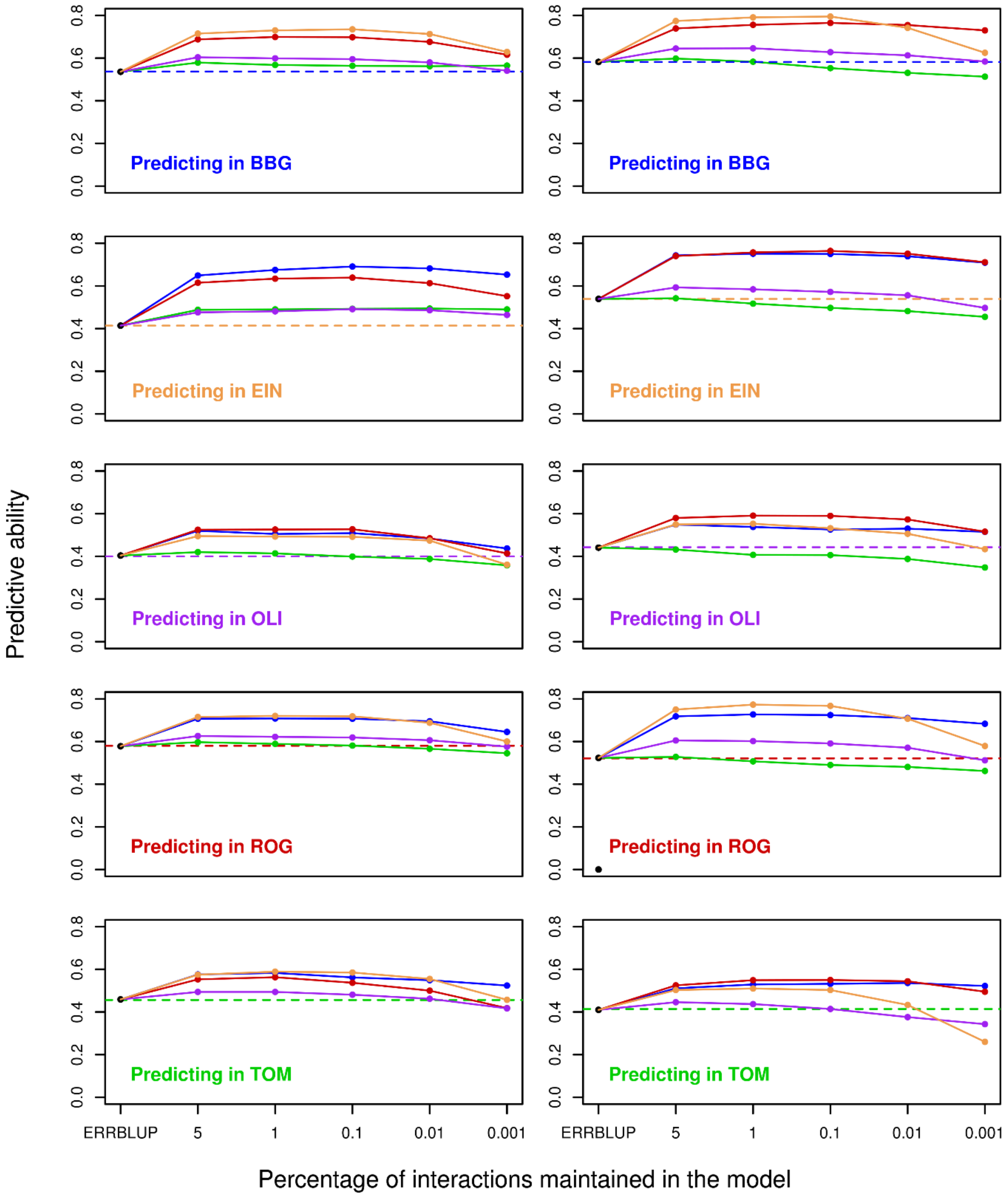


**Fig. S6a** Predictive ability for univariate GBLUP within environment (dashed horizontal line), univariate ERRBLUP within environment (black filled circle) and univariate sERRBLUP across environments when the SNP interaction selections are based on estimated effects variances (solid colored lines) for trait FF in KE (left side plots) and PE (right side plots). In each panel, the solid lines' color indicates the environment in which the relationship matrices determined by variable selection.

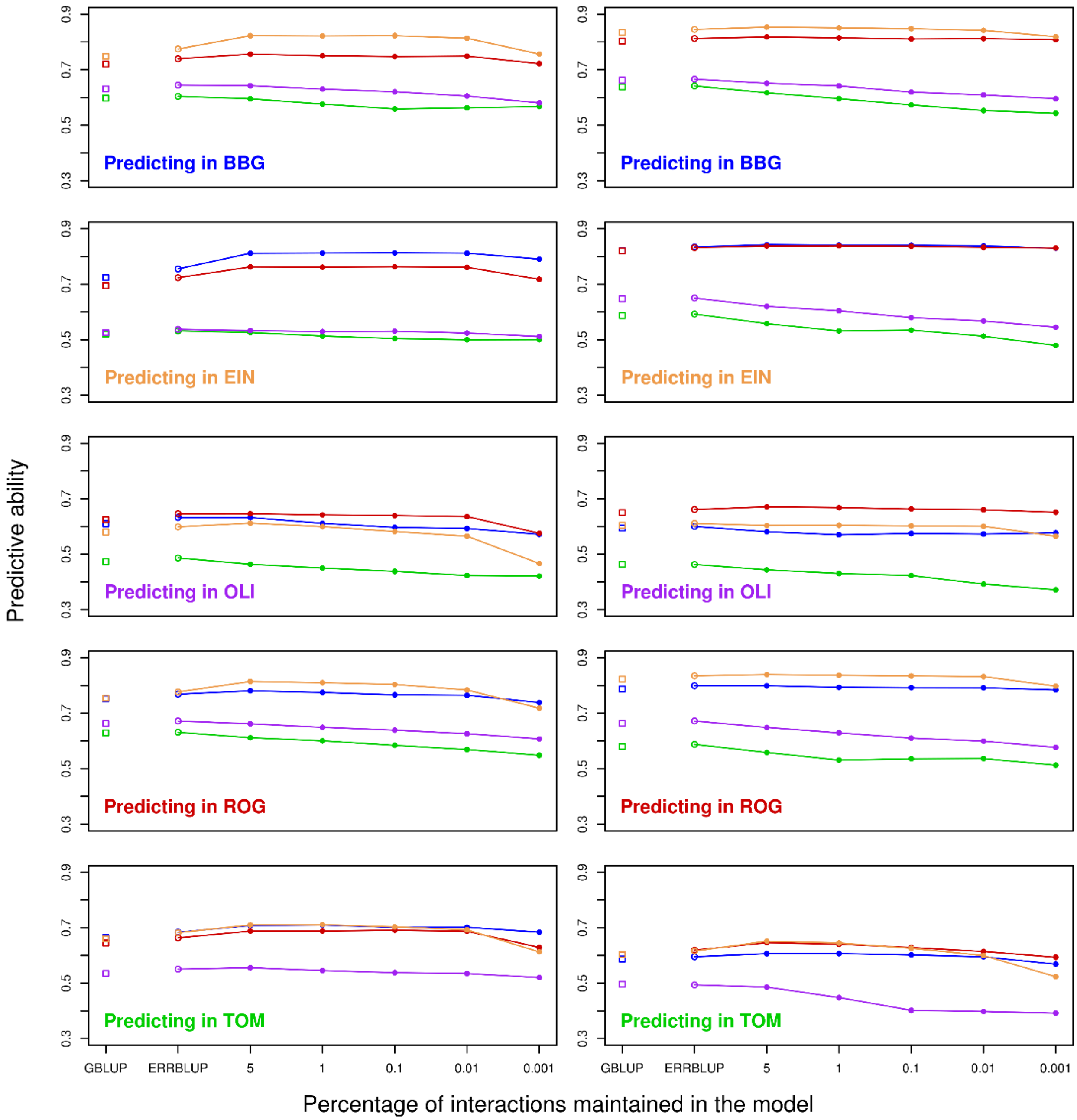


**Fig. S6b** Predictive ability for bivariate GBLUP (open squares), bivariate ERRBLUP (open circles) and bivariate sERRBLUP (filled circles and solid lines) when SNP interaction selections are based on estimated effects variances in Kemater (left side) and Petkuser (right side) for trait FF. In each panel, the solid lines' color indicates the additional environment used to predict the target environment.

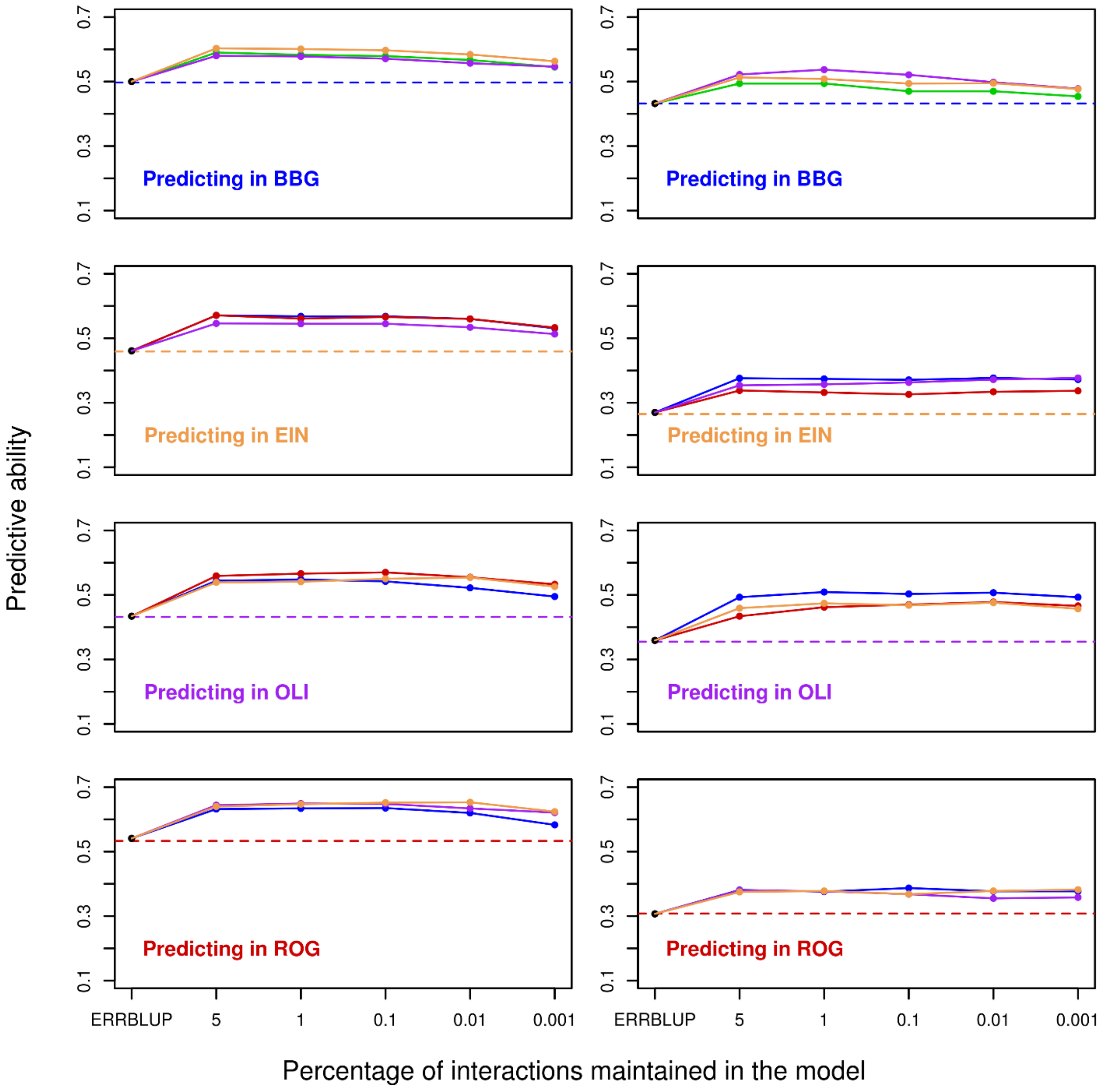


**Fig. S7a** Predictive ability for univariate GBLUP within environment (dashed horizontal line), univariate ERRBLUP within environment (black filled circle) and univariate sERRBLUP across environments when the SNP interaction selections are based on estimated effects variances (solid colored lines) for trait RL in KE (left side plots) and PE (right side plots). In each panel, the solid lines' color indicates the environment in which the relationship matrices determined by variable selection.

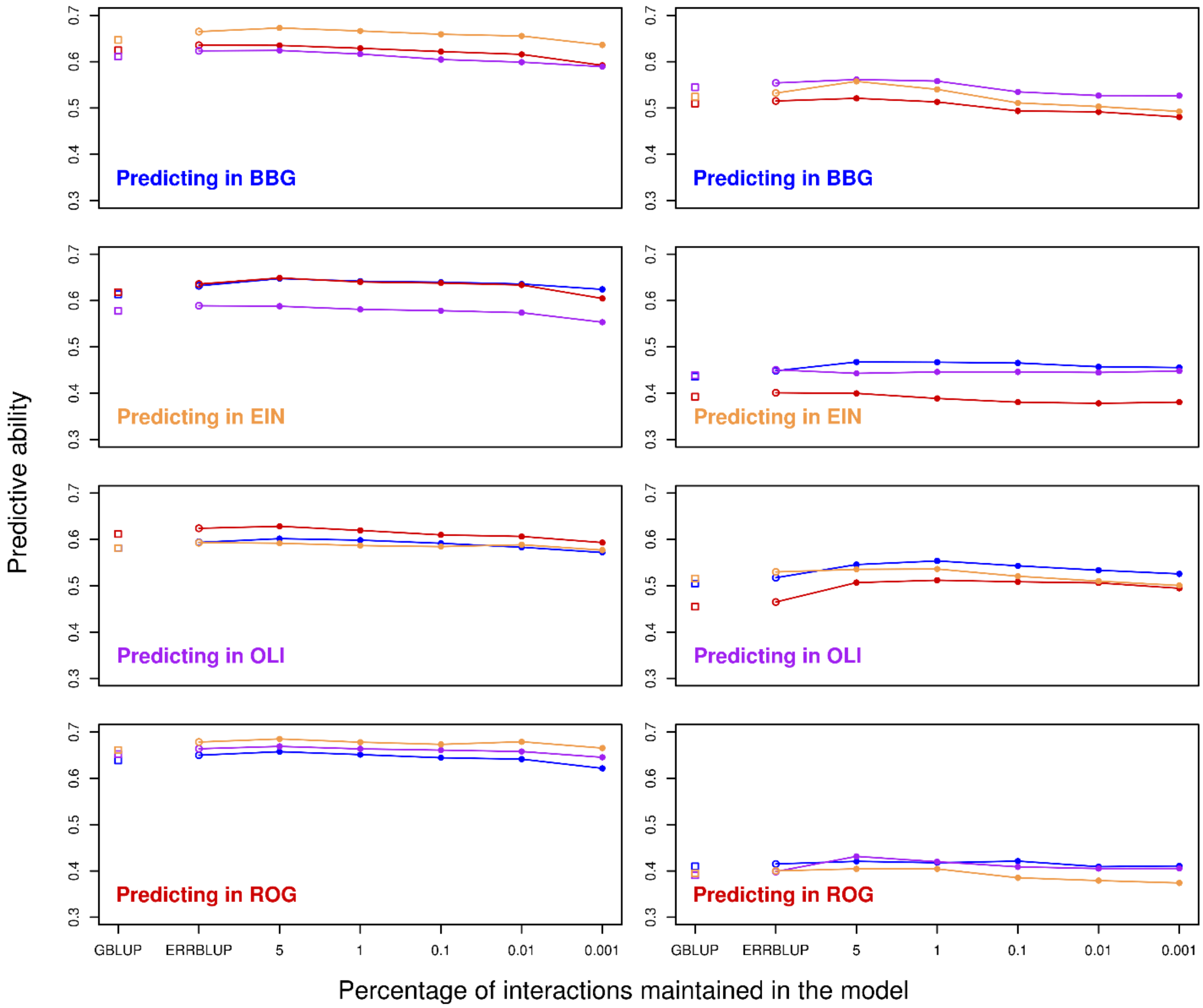


**Fig. S7b** Predictive ability for bivariate GBLUP (open squares), bivariate ERRBLUP (open circles) and bivariate sERRBLUP (filled circles and solid lines) when SNP interaction selections are based on estimated effects variances in Kemater (left side) and Petkuser (right side) for trait RL. In each panel, the solid lines' color indicates the additional environment used to predict the target environment

.

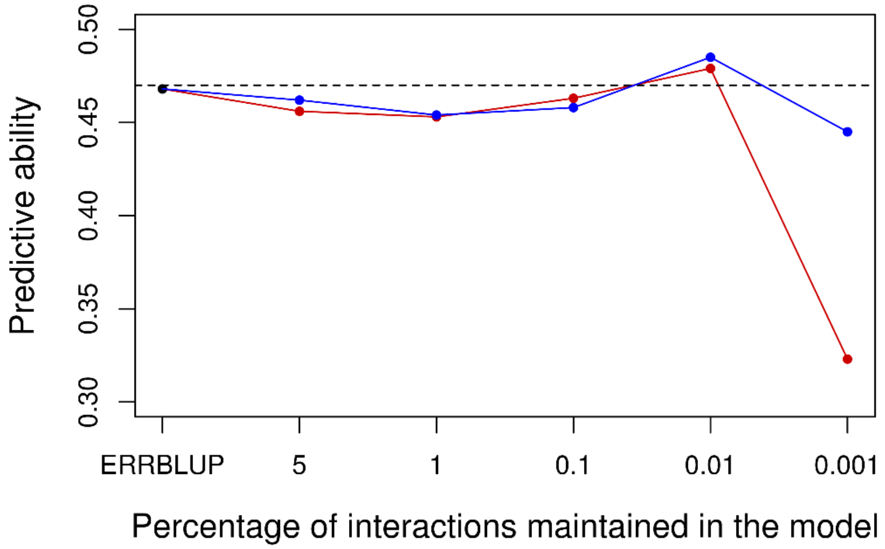


**Fig. S8** Predictive ability for univariate GBLUP within Bernburg (dashed horizontal line), univariate ERRBLUP within Bernburg (black filled circle) and univariate sERRBLUP when the SNP interaction selections are based on estimated effects variances (blue solid line) and estimated effect sizes (red solid line) within Bernburg for trait PH-V4 in Kemater.

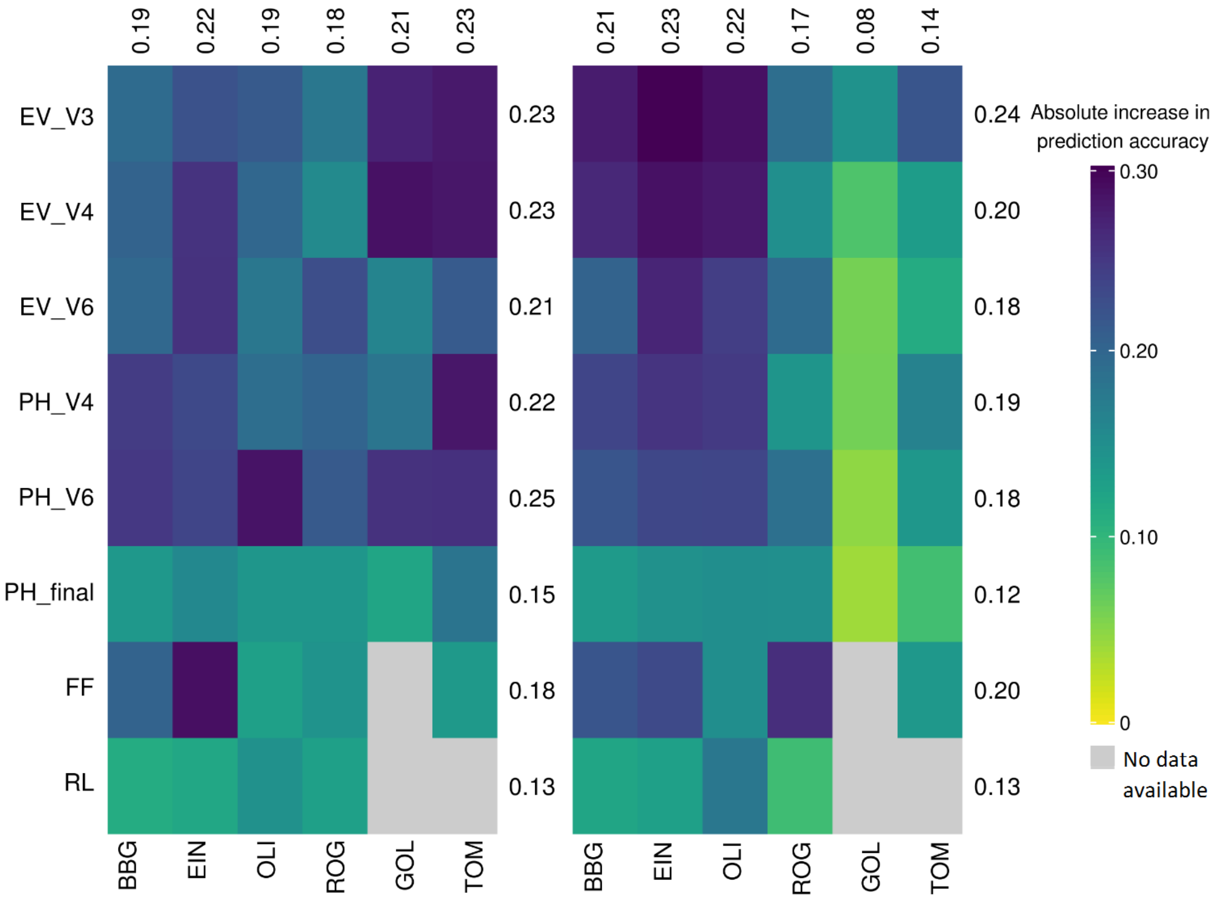


**Fig. S9a** Absolute increase in prediction accuracy from univariate GBLUP within environments to the maximum prediction accuracy of univariate sERRBLUP across environments when the SNP interaction selections are based on estimated effects variances in Kemater (left side plot) and in Petkuser (right side plot). The average of absolute increase in prediction accuracy for each trait and environments are display in rows and columns, respectively.

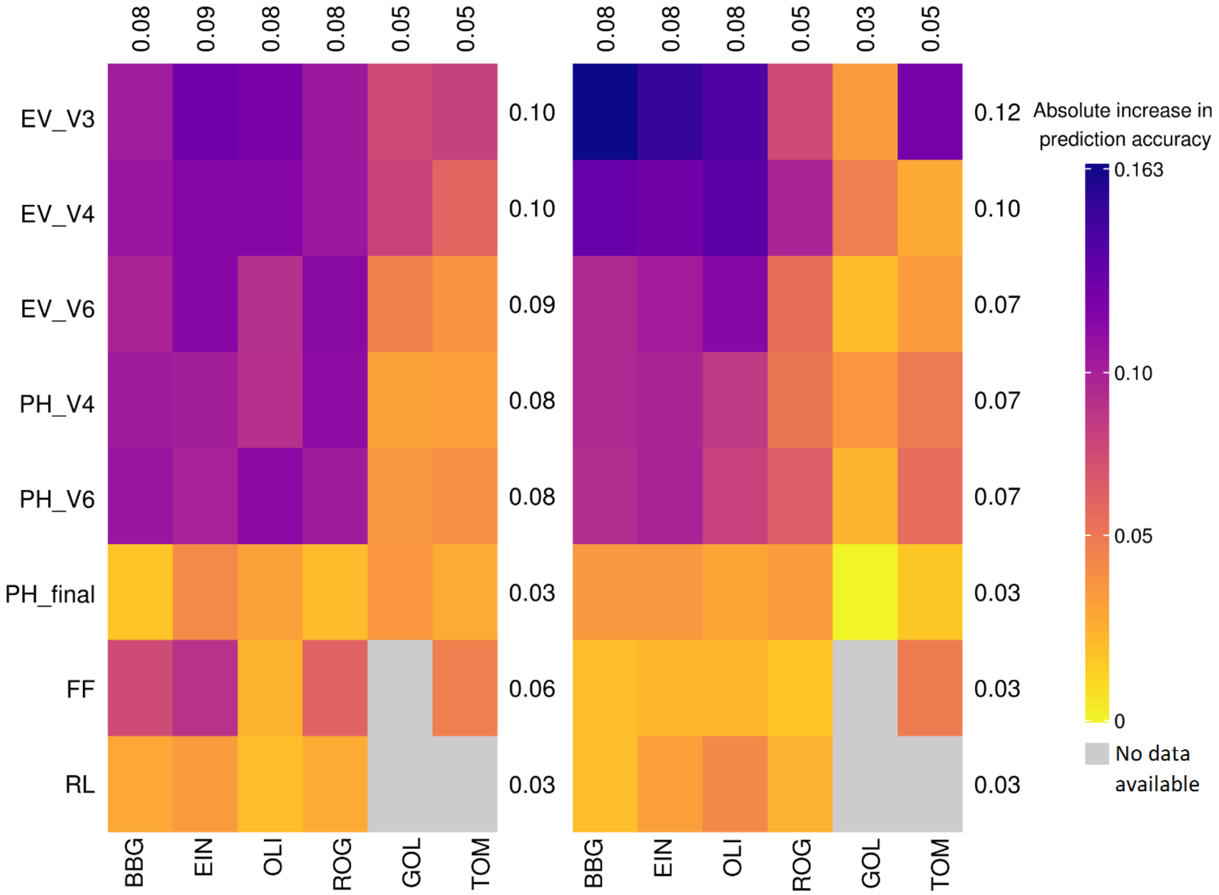


**Fig. S9b** Absolute increase in prediction accuracy from maximum bivariate GBLUP the maximum prediction accuracy of bivariate sERRBLUP when the SNP interaction selections are based on estimated effects variances in Kemater (left side plot) and in Petkuser (right side plot). The average of absolute increase in prediction accuracy for each trait and environments are display in rows and columns, respectively.

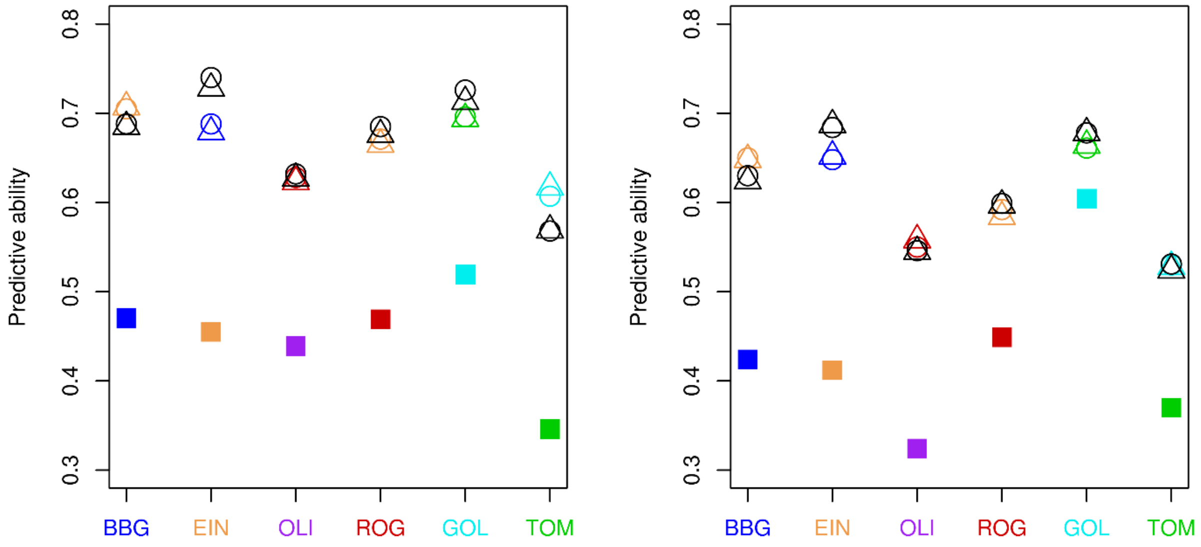


**Fig. S10** Comparison of predictive ability of univariate GBLUP within environments (filled squares) and with maximum predictive ability of univariate sERRBLUP when the SNP interaction selections are based on estimated effects sizes (circles) and estimated effect variances (triangles) for trait PH-V4 in Kemater (left side plot) and in Petkuser (right side plot). The colors dark blue, green, red, purple, light blue and orange represent the environments BBG, TOM, ROG, OLI, GOL and EIN respectively for prediction across a single environment. The color indicates the environment which had the maximum predictive ability for this respective target environment. However, the black symbols refer to prediction across the respective environment when the relationship matrices determined by variable selection in all the other five environments jointly.

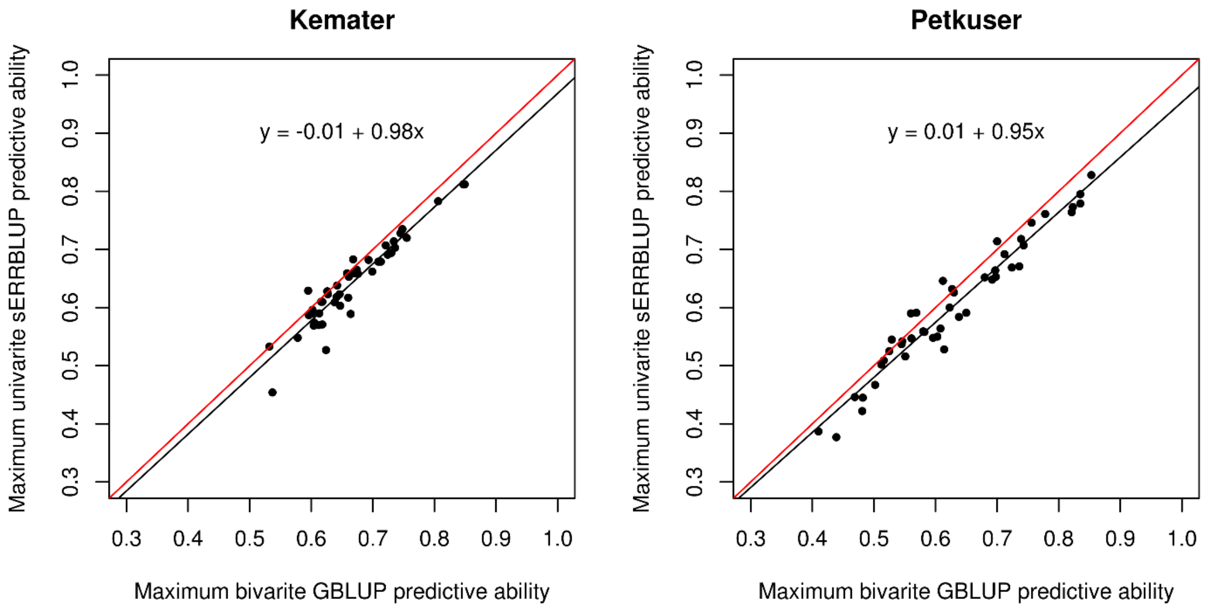


**Fig. S11** The comparison between the maximum predictive abilities of bivariate GBLUP within all six environments and the maximum predictive abilities of univariate sERRBLUP across environments for all traits in KE (left side plot) and PE (right side plot). In each plot, the diagonal line (red line) and the overall linear regression line (black line) with the regression formula are shown.

A PCA (Principal Component Analysis) graph in Fig. S12 demonstrates which environments are similar and which are relatively different from each other in terms of their weather data which are provided by Hölker *et al.* (2019).

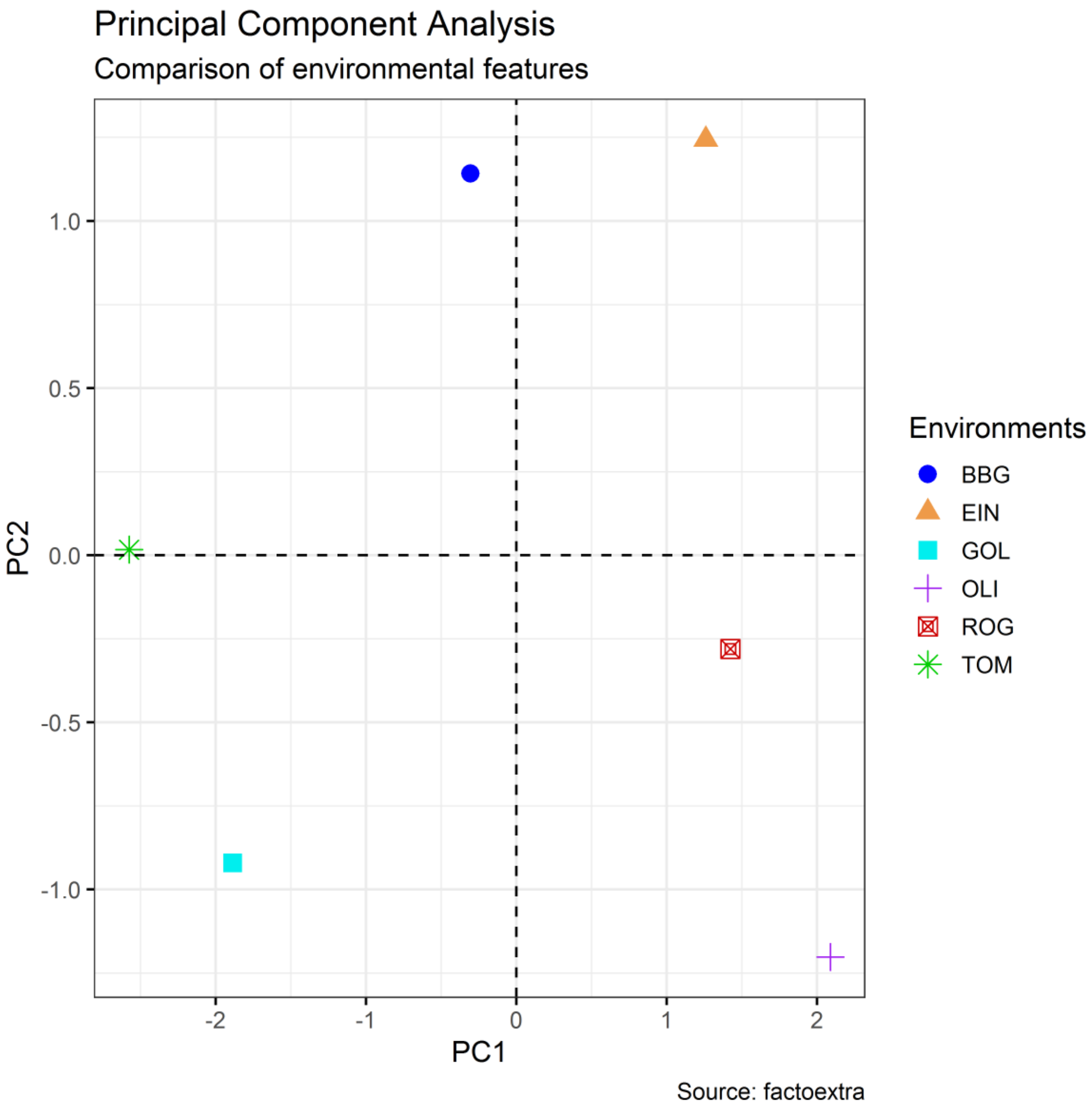


**Fig. S12** The position of each environment in the graph is obtained through five environmental features provided by Hölker *et al.* (2019) (altitude, total precipitation, and average, minimum and maximum daily temperature).
